# Cryo-EM studies of the four *E. coli* paralogs establish ABCF proteins as master plumbers of the peptidyl-transferase center of the ribosome

**DOI:** 10.1101/2023.06.15.543498

**Authors:** Shikha Singh, Riley C. Gentry, Ziao Fu, Nevette A. Bailey, Clara G. Altomare, Kam-Ho Wong, Matthew J. Neky, Chi Wang, Korak Kumar Ray, Ying-Chih Chi, Zhening Zhang, Robert A. Grassucci, Joachim Frank, Grégory Boël, Ruben L. Gonzalez, John F. Hunt

## Abstract

The genomes of most mesophilic organisms encode multiple <u>A</u>TP-<u>B</u>inding <u>C</u>assette <u>F</u> (ABCF) proteins. EttA, one of four *E. coli* paralogs, regulates synthesis of the first peptide bond on the ribosome dependent on ATP/ADP ratio, while <u>A</u>ntibiotic <u>Re</u>sistance factors (AREs), paralogs in other organisms, both regulate and directly mediate resistance to ribosome-targeted antibiotics. However, the physiological functions remain unclear for most paralogs, and the mechanism-of-action has yet to be rigorously established for any paralog. We herein present single particle cryogenic electron microscopy structures of ribosome complexes of all four *E. coli* ABCF paralogs (EttA, Uup, YbiT, and YheS), which, together with previously determined ARE structures, show that ABCFs control the binding geometry of the tRNA in the peptidyl-tRNA-binding (P) site on the ribosome. They modulate the position of its acceptor stem relative to the peptidyl transferase center (PTC) in a manner that can either promote (EttA and Uup) or disrupt (YbiT, YheS, and the AREs) proper catalytic geometry. The YbiT/70S reconstructions include a conformation with no density for ribosomal protein bL33, and structural analyses support the exchange of this sub-stoichiometric ribosomal protein being functionally related to conformational changes in YbiT controlled by sequence variations in the strongly non-canonical Signature Sequence in its first ABC domain. Our studies establish general structural/enzymological principles by which the ATPase activity of ABCF proteins controls translation elongation coupled to modulation of conformation and stereochemistry in the catalytic core of the ribosome.

---

The <u>A</u>TP-<u>B</u>inding <u>C</u>assette (ABC) Superfamily contains a vast number of mechanoenzymes that contain pairs of stereotyped ATPase domains that function as ATP-dependent mechanical clamps^1,2^. While most protein families within the ABC Superfamily function as transmembrane transporters, many families contain soluble proteins that mediate a diversity of biochemical functions ranging from DNA repair^3–5^ to metabolic cofactor biosynthesis^6,7^ and control of mRNA translation by ribosomes^8–12^. The ABCF family within the ABC Superfamily itself contains a large number of paralogous soluble protein subfamilies likely to have different molecular functions related to translation^8,13–17^ (**Fig. 1a**). We previously demonstrated that one ABCF subfamily, which we called “<u>E</u>nergy-dependent translation throttle A” or EttA, controls progression of the ribosome from the initiation phase to the elongation phase of translation dependent on ATP/ADP ratio^8^, a central physiological parameter that depends on cellular energy status^18,19^. Subsequent studies showed that a variety of ABCF subfamilies, referred to as antibiotic resistance elements (AREs), mediate resistance to ribosome-targeting antibiotics either directly^16,20–24^ or by regulating translation of other proteins that mediate resistance^9^. The genomes of most mesophiles encode multiple widely diverged ABCF paralogs, typically sharing only ∼30% sequence identity (**Fig. 1a-b**). For example, the human genome encodes three, the *Arabidopsis thaliana* genome encodes five, the *Saccharomyces cerevisiae* genome encodes two, and the *E. coli* K12 genome encodes four, called EttA, Uup, YheS and YbiT. The physiological functions remain unknown for the last three of these, as it does for the majority of ABCF subfamilies.

**Figure 1.**
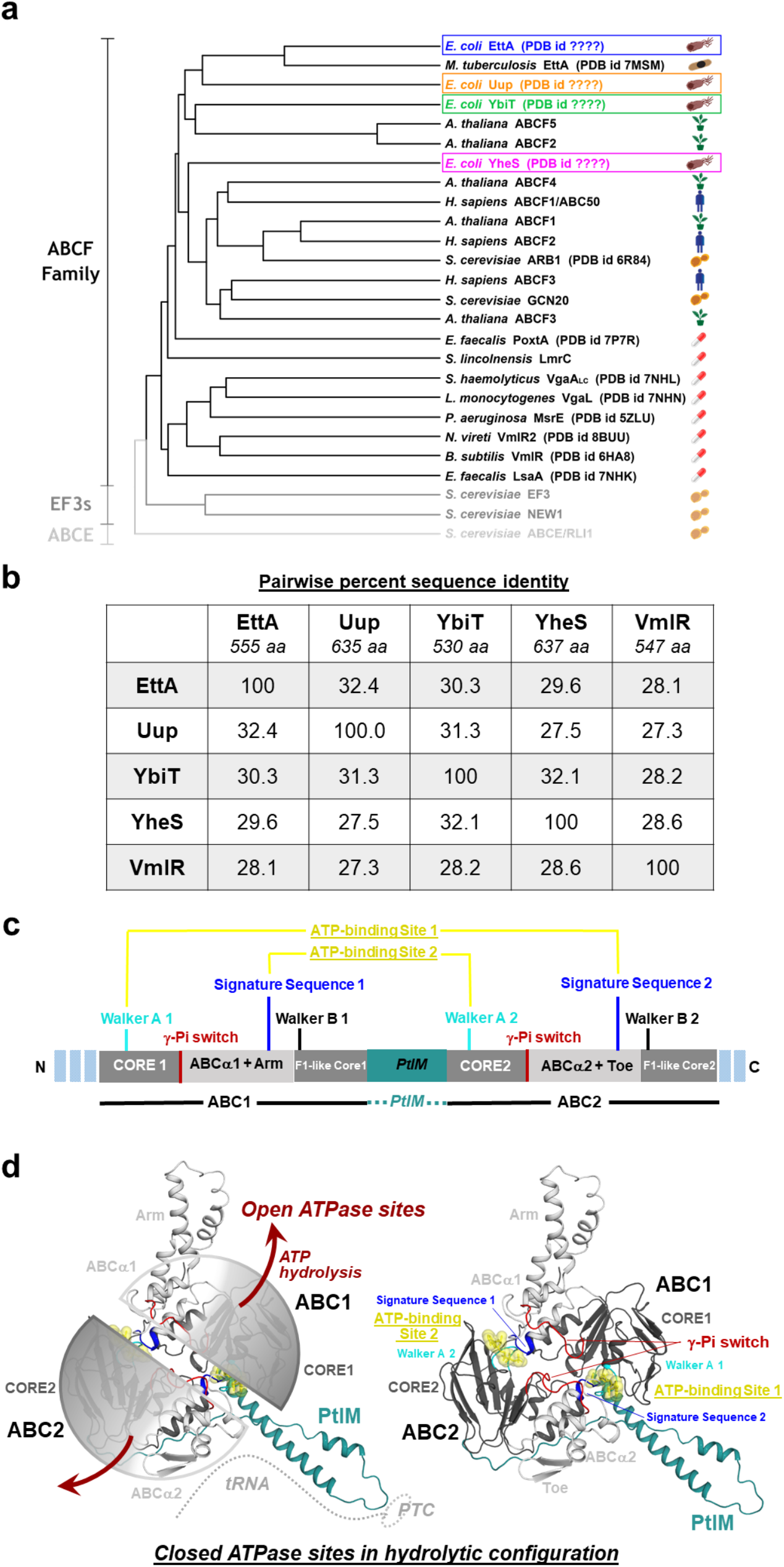
ABCF protein phylogeny and structural organization. **(a)** Cladogram depicting the evolutionary relationship between the four ABCF paralogs from *E. coli* (colored boxes), the EttA ortholog from *M. tuberculosis*, representatives of a diverse set of different ABCF paralog groups, the ARE ABCF proteins of known structure, and, for reference, three ABC Superfamily translation factors that are not in the ABCF family because they do not contain the PtIM domain (the yeast proteins EF3, New1, and ABCE. (**b)** Pairwise sequence identity matrix for the *E. coli* ABCF paralogs and VmlR, an ARE ABCF from *B. subtilis*. (**c)** Linear schematic showing the sub-domain organization of ABCF proteins, highlighting the sequence motifs involved in the formation of the conserved ATP sandwich dimer and hydrolysis of ATP. (**d)** Cartoon representations of EttA in its ATP-bound state, with ATP shown in yellow ball and stick representation. The two semi-transparent hemispheres in the image on the left outline the two ABC domains, ABC1 and ABC2, which bind two ATP molecules at their mutual interface. The image on the right shows the same ribbon diagram of EttA but without the semi-transparent hemispheres and with sequence motifs involved in ATP binding/hydrolysis highlighted (the Walker A in cyan, the Signature Sequence in blue, and the γ-P_i_ switch in red). Each ABC domain comprises an ATP-binding core subdomain (CORE 1 and CORE 2 in ABC1 and ABC2, respectively) and an α-helical subdomain (ABCα1 and ABCα2, respectively) that move as rigid bodies in different conformations of the protein. The scarlet arrows on the left indicates the general direction of domain rotation upon ATP hydrolysis as inferred from comparison of the nucleotide-free crystal structure^8^. The PtIM (<u>P</u>-site tRNA <u>I</u>nteraction <u>M</u>otif, shown in teal) connects the two ABC domains. Its interaction sites with the PTC (<u>P</u>eptidyl <u>T</u>ransferase <u>C</u>enter) of the ribosome and the P-site tRNA are schematized in the image on the right. The Arm motif (light gray) is an α-helical hairpin inserted in ABCα1, while the toe is a β-hairpin inserted at the same site in ABCα2.

The distinguishing sequence/structure feature of ABCF proteins is an ∼80 residue α-helical hairpin called the <u>P</u>-site tRNA <u>I</u>nteraction <u>M</u>otif (PtIM) that links the two tandem ABC ATPase domains encoded in a single polypeptide chain (**Fig.1c**). Our initial low resolution single particle cryogenic electron microscopy (cryo-EM) structure of EttA^25^ and subsequent higher resolution structures of the EttA ortholog from *Mycobacterium tuberculosis*^26^ (*Mtb*) as well as eight^20–24,27^ different ARE proteins all show the ABCF proteins bound in the tRNA Exit (E) site of the 70S ribosome, and these structures all have the PtIM interacting with the P-site tRNA proximal to the peptidyl transferase center (PTC) in the large (50S) ribosomal subunit. Furthermore, a cryo-EM structure of a ribosome recycling complex from *Saccharomyces cerevisiae* containing the ABCF protein Arb1 shows its PtIM bound to the large ribosomal subunit in a similar geometry^28^.

The PtIMs from different ABCF proteins share a relatively low level of sequence identity, but they are nonetheless reliably identified by sequence-profiling algorithms including the algorithm used to construct the PFAM database^29^, which identifies the PtIM as a conserved domain called PF12848. The tip of the PtIM, which connects its two α-helices, varies in sequence and length in different ABCF proteins, and these variations influence the antibiotic specificity of different ARE proteins^16,20–24,27,30–32^. For this reason, some papers have used the alternative name “antibiotic resistance domain” (ARD) for the PtIM. However, it interacts with the ribosome in an equivalent geometry in ARE proteins and in ABCF proteins that have no detectable activity mediating antibiotic resistance^8,9,12,16,20–22,24,25,33,34^, and it has not yet been possible to explain the qualitative features of the antibiotic specificity profiles of different AREs based on specific sequence variations in the tip or other regions of the PtIM^9,12,16,20–22,24,33,34^. Furthermore, the results presented below suggest that ABCF proteins with different primary functions could also mediate some degree of antibiotic resistance, which would imply functional continuity between the ARE protein and other ABCF proteins in the course of evolution.

ABC domains (**Fig. 1c-d**) contain two mostly rigid structural modules, an ATP-binding core subdomain (herein called the CORE subdomain) and an α-helical subdomain (herein called the ABCα subdomain) that are connected by a flexible ∼6 residue linker called the <u>γ</u>-<u>P</u>hosphate <u>S</u>witch (γP_i_-Switch) or alternatively the Q-Loop^5^. The catalytically active ATP-bound conformation of ABC Superfamily enzymes, which generally show 2-fold positive cooperativity^35^, comprises an “ATP-sandwich” complex^36–40^ in which two ATP molecules are bound in composite active sites, each formed between the Walker A/B motifs in the CORE subdomain in one ABC domain and the “Signature Sequence” in the ABCα subdomain in a second ABC domain (**Fig. 1c-d**). The Signature Sequence, which is the hallmark of ABC Superfamily ATPase domains, has a canonical sequence of LSGGQ but for reasons that remain unclear is most commonly LSGGE in ABC domains in ABCF proteins. The amino acids and sequence motifs involved in ATP binding and hydrolysis, which represent the most strongly conserved sequence features within ABC domains (**Fig. 1c-d**), are described in greater detail in the ***Supplementary Discussion*** section in the ***Supplementary Information*** (***SI***) for this paper. The clamp-like closure of the ABC domains during formation of the ATP-sandwich complex, which is driven by the energy of ATP binding^38^, represents the mechanochemical “power-stroke” that drives transmembrane transport in a wide variety of ABC Superfamily proteins^38,41–50^. We previously proposed that a second mechanochemical power-stroke could be driven by transient electrostatic forces generated in the active site during the ATP hydrolysis reaction^38,51^ (dark red arrows in **Fig. 1d**), and we recently verified this prediction for a widely studied vitamin B_12_ transporter in the ABC Superfamily^52,53^.

The function of ABC domains as ATP-dependent mechanical clamps was established primarily via studies in which the catalytic glutamate at the end the Walker B motif in the CORE subdomain is replaced by glutamine^5,35,38,41,54^, an isosteric amino acid that is incapable of performing general base catalysis and therefore blocks ATP hydrolysis. Isolated ABC domains from homodimeric transmembrane transporters in the ABC superfamily purify as monomers in the absence of nucleotides and remain monomeric in the presence of ADP, ATP, and non-hydrolyzable analogs of ATP (including in the presence of vanadate or aluminum/beryllium fluoride). However, in the presence of the glutamate-to-glutamine (EQ) mutation in the catalytic base, addition of ATP leads to highly efficient dimerization due to encapsulation of two molecules of ATP at the interface between the two ABC subunits in the ATP-sandwich complex^5,35,38,41,54^ (**Fig. 1d**). Non-hydrolyzable ATP analogs fail to drive formation of this complex because they generally have reduced binding energy compared to ATP due to substitution of one of the five atoms in its terminal phosphate group by an atom with different chemical characteristics. In contrast, EQ mutations in the catalytic base have proven effective in stabilizing structures modeling the ATP-bound pre-hydrolysis conformations of a very wide variety of ABC Superfamily ATPases^8,35,38,44,55–57^. In the case of ABCF proteins, the EQ mutation has been introduced into both of the tandem ABC domains, resulting in “EQ_2_” constructs that have been broadly successful in kinetically trapping their functional ATP-bound conformations^8,12,15,20–22,24,25,33,34,38,58^.

Formation of the catalytically active ATP-sandwich complex requires proper alignment of the two subdomains within both of the two interacting ABC domains in order to enable simultaneous binding of ATP at the two composite active sites formed at their mutual interface (**Fig. 1d**). Disruption of this alignment due to changes in the relative interaction geometry of the CORE and ABCα subdomains in one of the ABC domains prevents formation of the active ATP-sandwich complex in which two ATPase active sites are configured in proper catalytic geometry at its interface with the other ABC domain^59^. Such CORE/ABCα inter-subdomain movements within an ABC domain frequently result in the withdrawal of an invariant, catalytically important glutamine residue at the N-terminus of the γP_i_-Switch from the ATPase active site in that domain. These coupled conformational effects modulating active site formation are exploited for allosteric control of ATPase activity in some ABC Superfamily enzymes^36,42,59,60^.

We herein employ cryo-EM image reconstruction^25,61–64^ to elucidate near-atomic resolution structures of 70S ribosome complexes containing ATP-bound EQ_2_ mutants of all four *E. coli* ABCF paralogs. Comparison of these structures to a dozen previously determined structures of ribosome-bound complexes of different ABCF proteins indicates that the ATP-bound conformation controls the location and orientation of the acceptor stem of the P-site tRNA relative to the PTC in the catalytic core of the ribosome. Depending on the structure of the PtIM, different ABCF proteins either promote adoption of the proper geometry for peptide bond formation or pull the acylated acceptor stem of the P-site tRNA away from the PTC into non-catalytic geometries. ABCF proteins will perform surveillance of ribosome complexes with an empty E site, and the specificity for different complexes is likely to be determined at least in part by the activation energy required to move the acceptor stem of the P-site tRNA and covalently bonded amino acid or polypeptide, which will be strongly influenced by the sequence/structure of the PtIM.

## Results and Discussion

### Cryo-EM reconstructions of 70S ribosome complexes of EQ_2_ mutants of the E. coli ABCFs

To gain insight into the principles underlying ABCF protein function, we determined structures of hydrolysis-deficient “EQ_2_” mutants of all four *E. coli* ABCF paralogs bound to *in-vitro*-assembled *E. coli* 70S ribosome complexes. Biochemical and physiological studies indicate that EttA functionally interacts with both translation Initiation <u>C</u>omplexes (ICs) harboring fMet-tRNA^fMet^ in the P site of the ribosome and translation elongation complexes (ref.^8^ and manuscripts in preparation^65,66^), although the exact configuration of the ribosome in the latter complexes has not been established. Given this ambiguity combined with the observation that overexpression of YbiT partially complements the effects of knocking out the *ettA* gene in *E. coli* (accompanying paper^67^ and manuscript in preparation^66^), we prioritized structural characterization of the EQ_2_ mutants of all four *E. coli* ABCF paralogs bound to 70S ICs (**Figs. 2 & S1-S4** and **Tables S1-S2**), which are readily assembled *in vitro* in their physiological state using purified components. We also determined two reference 70S IC structures without a bound ABCF protein using ribosomes imaged in one of our ABCF-containing samples (**Table S3**). We furthermore probed the dependency of the EQ_2_-EttA and EQ_2_-YbiT interactions on the identity of the tRNA and its acylation state by determining their structures bound to 70S <u>E</u>longator tRNA <u>C</u>omplexes (ECs) harboring deacylated tRNA^Val^ in the P site of the ribosomes (**Figs. S5-S7** and **Table S1-S2**). These “PRE^-A^_Val_” complexes represent a <u>pre</u>-translocation configuration of the ribosome <u>minus</u> a peptidyl-tRNA in the <u>A</u> site, which is not a physiological state during the normal elongation cycle but has been a powerful tool for characterization of ribosome dynamics^68–70^.

**Figure 2.**
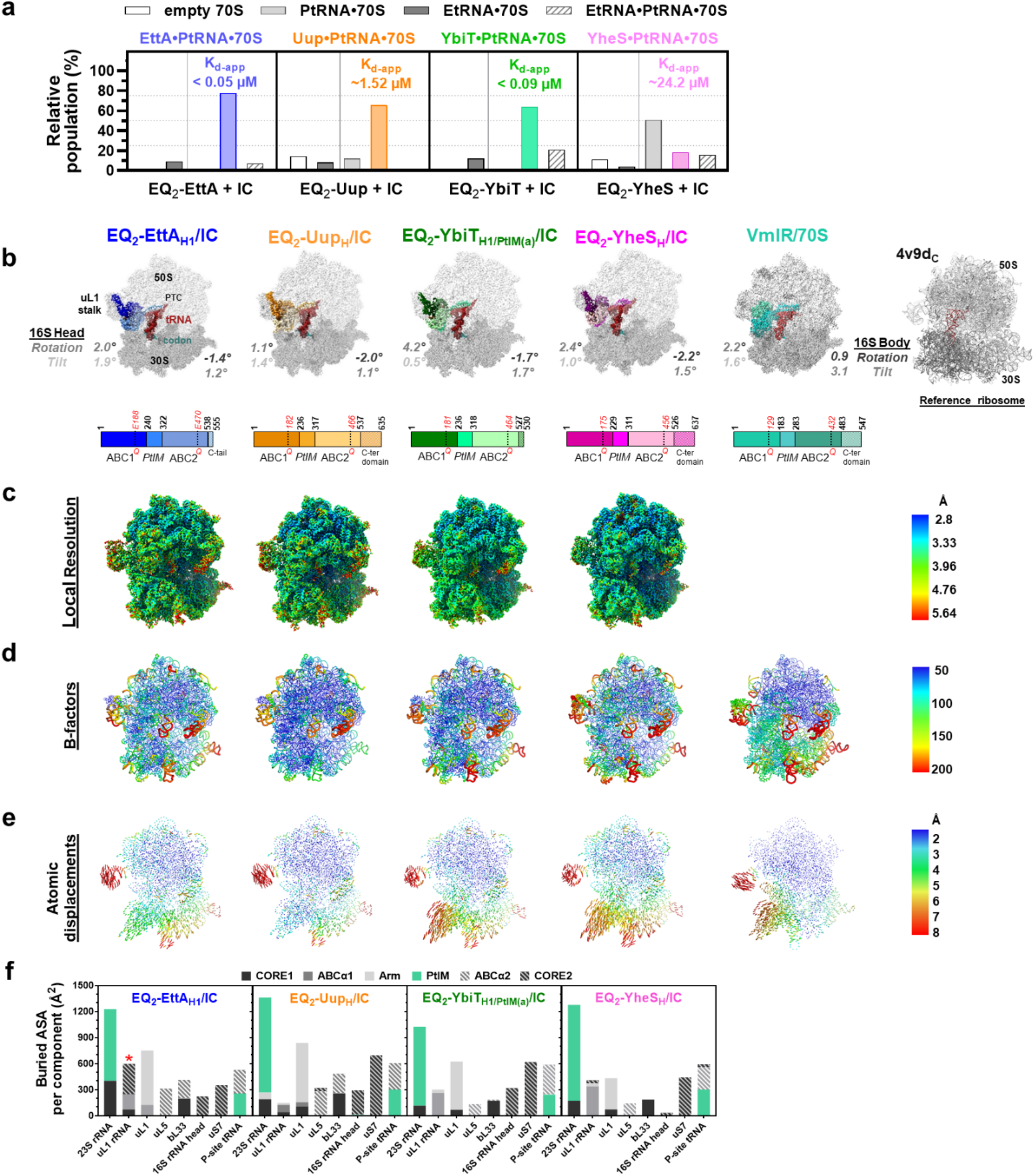
Cryo-EM structures of ATP-bound hydrolysis-deficient EQ_2_ mutants of E. coli EttA, Uup, YbiT, and YheS bound to homologous 70S ribosomal initiation complexes (ICs). **(a)** Population distribution of 70S particles in each dataset with *E. coli* EQ_2_-ABCF paralogs added to 70S ICs. **(b)** Density maps of the *E. coli* EQ_2_-ABCF-bound 70S ICs showing 30S subunit in dark gray, 50S subunit in light gray, P-site tRNAs in crimson red, mRNA in teal, and each of the *E. coli* ABCFs and the ARE ABCF (VmlR) color coded as in the bar below. Color bars show the general sub-domain organization of the four *E. coli* ABCFs and VmlR with the residue numbers demarcating each subdomain and the position harboring the E to Q mutation in each subdomain shown in red. Values calculated for the intersubunit rotations in these complexes with respect to the reference ribosome (4v9d_C_ (PDB ID 4v9d) shown in cartoon representation) are indicated adjacent to each complex. **(c)** Density maps of the EQ_2_-ABCF-bound 70 ICs colored according to local resolution. **(d)** Cartoon putty representation of the EQ_2_-ABCF-bound 70 ICs, high B-factor corresponds to red thicker regions of the tube, the color coding varies blue to red for low to high B-factor values as shown in the bar on the right. **(e)** Displacement (Å) of the phosphates in the 16S and 23S rRNA in the EQ_2_-ABCF-bound 70 ICs relative to those in the 4v9d_C_ complex (aligned on 23S rRNA), color coded blue (smallest) to red (largest). For calculating displacements in VmlR bound complex P-site-tRNA-bound *B. subtilis* ribosome (PDB ID 6ha1) was taken as reference. (f) Bar graphs showing buried surface area between the subdomains in *E. coli* EQ_2_-ABCF paralogs and the ribosomal elements and the P-site tRNA, in the EQ_2_-ABCF bound 70S ICs. The red asterisk (*) in the EQ_2_-EttA panel indicates unique interaction between the C-terminal tail extending from its CORE2 to the uL1 stalk not observed in the other *E. coli* ABCF paralogs.

While our cryo-EM samples yielded a single near-atomic-resolution reconstruction for the 70S ICs of EQ_2_-Uup and EQ_2_-YheS and the 70S ECs of EQ_2_-EttA and EQ_2_-YbiT, they yielded multiple near-atomic-resolution reconstructions for the 70S ICs of EQ_2_-EttA and EQ_2_-YbiT (**Figs. 2 & S1-S7** and **Tables S1-S2,**). In the case of EttA, the two conformational classes show a modest oscillation in the orientation of the ABC domains of EttA coupled to tilting the of uL1 stalk in the large (50S) ribosomal subunit and a small rotation of the “head” domain of the small (30S) ribosomal subunit (**Fig. S1j-k**). These conformational classes show the acceptor stem of the P-site tRNA making nearly identical interactions with the PTC of the ribosome as each other and the three previously determined structures of *Mtb*EttA^26^ showing density for the terminal CCA in the acceptor stem (**Fig. S9**). They seem likely to represent a continuous oscillation in the binding geometry of ATP-bound EttA in the E site of the ribosome (**left in Movie S1**). In contrast, the multiple classes reconstructed for YbiT show substantial conformational differences that are described in detail below.

The single classes reconstructed for the 70S ECs of EQ_2_-EttA (**Fig. S5f-g**) and EQ_2_-YbiT (**Fig. S6f-g**) are both very similar in conformation to largest class reconstructed for the 70S IC complex of the same ABCF protein complex except locally near the elbow region of the P-site tRNA, which differs in structure in tRNA^fMet^ *vs.* tRNA^Val^. These results demonstrate that the structure of the tRNA bound in the P site and its acylation state do not significantly alter the structural characteristics of ribosome complexes of these two EQ_2_-ABCF proteins, consistent with the results of biochemical and biophysical assays presented in the accompanying paper^67^ showing these proteins have similar affinities and functional interactions with ICs and PRE^-A^_Val_ complexes.

In this paper, we label the observed ABCF protein conformations using a subscript that encodes the configuration of the tandem ATP-binding sites (H for <u>H</u>ydrolytic, N for <u>N</u>on-hydrolytic, or I for Intermediate) and also in the case of YbiT the different binding modes of its PtIM on the 23S rRNA in the 50S subunit of the ribosome (PtIM(a) or PtIM(b)). Finally, the configuration of the 70S ribosome in each structure is designated as IC or EC, representing the initiator tRNA or PRE^-A^_Val_ complexes, respectively, with the addition of the subscript “L33^-^” for the YbiT complex structures in which no density is observed for ribosomal protein bL33, which are explained below. The complete set of 11 ABCF/70S ribosome complexes reported in this paper are thus designated as EQ_2_-EttA_H1_/IC, EQ_2_-EttA_H2_/IC, EQ_2_-EttA_H1_/EC, EQ_2_-Uup_H_/IC, EQ_2_-YheS_H_/IC, EQ_2_-YbiT_H1/PtIM(a)_/IC, EQ_2_-Ybi_TH2/PtIM(a)_/IC, EQ_2_-YbiT_I/PtIM(b)_/IC, EQ_2_-YbiT_N1/PtIM(a)_/IC_L33^-^_, EQ_2_-YbiT_N2/PtIM(b)_/IC, and EQ_2_-YbiT_N1/PtIM(a)/ECL33^-^_. The reasons underlying the Hydrolytic/Non-hydrolytic/Intermediate designations are explained in detail below, and the names and composition of all complexes are summarized in **Tables S1 and S2**.

The IC and EC structures reported in this paper were reconstructed at resolutions ranging from 3.0-3.3 Å and 3.0-3.6 Å (**Tables S1-S3**), respectively. The final maps from non-uniform refinement in cryoSPARC^71^ all have similar values for gold-standard <u>f</u>ourier <u>s</u>hell <u>c</u>orrelation (FSC) resolution, d_99_ resolution^72^, and also the d_model_ resolution for the corresponding coordinate model refined in PHENIX^72^. The coordinate models have generally excellent stereochemical characteristics (**Tables S1-S3**).

Our cryo-EM structures were determined in the presence of 2 mM Mg-ATP, and they all show density consistent with unhydrolyzed ATP being bound in both ATP-binding sites in all of the ABCF proteins (**Fig. S8a**), consistent with stabilization of pre-hydrolysis conformations by the EQ_2_ mutations^8,35,38,44,55–57^. They all show clearly resolved and continuous density from their N-terminal methionines through the C-termini of their ABC2 domains, although their Arm regions generally show somewhat diffuse density (**Figs. S1-S6**). EttA has an 18-residue C-terminal extension after ABC2 that can be traced except for its final three residues, and residues 446-551 in this segment show very clear density sandwiched between the ABCα1 subdomain of EttA and the uL1 stalk of the ribosome, as previously reported for *Mtb*EttA^26^. The final three residues in YbiT are disordered in some of its conformational classes but not others. Uup and YheS both have ∼100 residue C-terminal domains that are not visible in the density maps and therefore not modeled in our structures. There was clear density for all bases in the P-site tRNAs in all of the structures (**Figs. S1-S6**). However, only 6-9 bases of the 32-base model mRNA showed clear density, which included the codons in the P- and A-sites and in some cases the E-site but no interpretable density beyond that in the mRNA exit channel (**Fig. S8b-c**). The uL1 protein, uL1 stalk rRNA, and some peripheral helices in both the 23S and 16S rRNAs show diffuse density, which limited atomic modeling in these regions to rigid-body fitting of segments from reference structures.

### Estimation of ABCF affinity for 70S ICs and ECs based on cryo-EM population distributions

Following *in vitro* assembly, the complexes were applied to grids and imaged using direct electron detectors on 300 kV electron microscopes. (See Methods.) Systematic classification of the particles in the datasets using RELION/cryoSPARC^73,74^ revealed multiple structural classes in all of the datasets (**Figs. S1-S6**). The distribution of molecular complexes varied in the different datasets, but all included free 30S and 50S ribosome subunits as well as a substantial population of ABCF-bound 70S ribosomes harboring a P-site tRNA (**Figs. 2 & S7**). The datasets also showed varying content of other complexes, including empty 70S ribosomes, 70S ribosomes with a bound ABCF protein but no P-site tRNA, 70S ribosomes harboring a P-site tRNA but no ABCF protein, 70S ribosomes harboring an E-site tRNA and an empty P-site, and 70S ribosomes harboring both E-site and P-site tRNAs.

The populations assigned to different classes during cryo-EM data processing were used to estimate the apparent binding affinities of the ATP-bound conformations of the EQ_2_ -ABCF factors for the ribosome complexes in which they were visualized (**Figs. 2a & S7a**). The total concentration of the 50S ribosomal subunit during the *in vitro* assembly reaction was assumed to match the sum of all particles assigned to 50S-subunit containing classes, which provided an estimate for the absolute concentration of each ribosome complex in the cryo-EM sample. The free concentration of the EQ_2_-ABCF protein was calculated by subtracting the concentration of every ribosome complex containing it from its total concentration during the *in vitro* assembly reaction, enabling the apparent binding affinities for ribosomes with a P-site tRNA (top equation) and without one (bottom equation) to be calculated as follows:

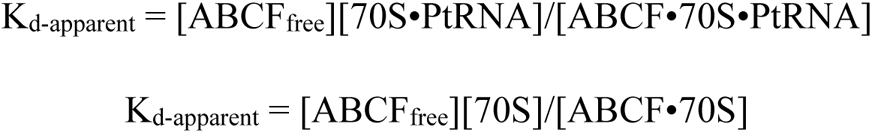

The symbol PtRNA represents a tRNA bound in the P site of the ribosome complex. When cryo-EM data processing did not yield a 70S•PtRNA or ABCF•70S class, the population of that class was assumed to be less than the minimum population of any class reconstructed in the same sample. Given the identity of the missing classes in the different samples, this assumption yields an upper bound for the apparent affinity for ribosomes harboring a P-site tRNA and a lower bound for the apparent affinity for empty ribosomes.

These analyses indicate that, in the presence of 2 mM ATP, the apparent binding affinities of the ATP-bound conformations of the ABCF proteins for the 70S IC span roughly two-orders of magnitude in the nanomolar-to-micromolar concentration range: EQ_2_-EttA < 0.05 µM, EQ_2_-YbiT < 0.09 µM, EQ_2_-Uup ∼1.5 µM, and EQ_2_-YheS ∼24 µM (**Fig. 2a**). The apparent binding affinities of EQ_2_-EttA and EQ_2_-YbiT for the 70S EC (< 0.33 and < 0.37 µM, respectively – **Fig. S7a**) are similar to their affinities for the IC. However, compared to their respective affinities for 70S ICs and ECs, the *E. coli* EQ_2_-ABCF proteins all show substantially weaker apparent binding affinities for 70S ribosomes without a bound tRNA: EQ_2_-EttA ∼62 µM, EQ_2_-YbiT ∼1.2 µM, EQ_2_-Uup > 12 µM, and EQ_2_-YheS > 19 µM (**Figs. 2a & S7a**). These affinities suggest that the *E. coli* ABCF proteins other than YbiT are likely to have little interaction *in vivo* with 70S ribosomes without a bound tRNA. Except for YheS, all of the apparent binding affinities inferred from our cryo-EM samples are consistent with the results of single-molecule fluorescence resonance energy transfer (smFRET) measurements of the interaction of the same EQ_2_-ABCF proteins with a PRE^-A^_Phe_ complex^67^, which is similar to the PRE^-A^_Val_ complex in our ECs. The significantly higher apparent binding affinity of EQ_2_-YheS exhibited in the smFRET experiment (< 1 µM) likely reflects specificity for the elongator tRNA used in that experiment compared to the initiator tRNA used to produce our EQ_2-_YheSH/IC structure. The physiological significance of the observed differences in the binding affinities of the four paralogs is discussed in the accompanying paper^67^.

### ABCF factors all bind in the E site of the ribosome

All four *E. coli* ABCF paralogs bind in the E site of the ribosome, and the same regions of the ABCF proteins make contacts with the same ribosomal features in all structures (**Figs. 2 & S7** and **Movie S1**), even though the specific molecular contacts at these sites are not phylogenetically conserved (**Fig. 3**) in these widely diverged (**Fig. 1b**) paralogs. The solvent-accessible surface area buried in the contacts between each of the subdomains in the ABCF proteins and specific components of the ribosome is quantified in **Figs. 2f & S7g**, and a verbal description of these interactions is provided in the ***Supplementary Discussion*** section in the ***SI*** for this paper. The PtIM, which connects the C-terminus of ABC1 to the N-terminus of ABC2, is the distinguishing feature of the ABCF protein family. It consistently makes extensive packing interactions with both 23S rRNA in the 50S subunit and the P-site tRNA (**Figs. 2b, 2f, 4, S7b, & S7g**). The magnitude of solvent-accessible surface area buried at the ABCF-ribosome contact sites varies significantly between the ribosome complexes of the different *E. coli* ABCF paralogs, again reflecting their phylogenetic divergence, but it is substantially more consistent for the PtIM than their other structural modules (**Figs. 2f & S7g**). The available structures of 70S ribosome complexes of ARE ABCF proteins^8, 20–25,27,33,34^ all show them binding in an equivalent geometry in the E site and make similar structural interaction to the *E. coli* ABC paralogs (**Figs. 2, 4, & S13**).

**Figure 3.**
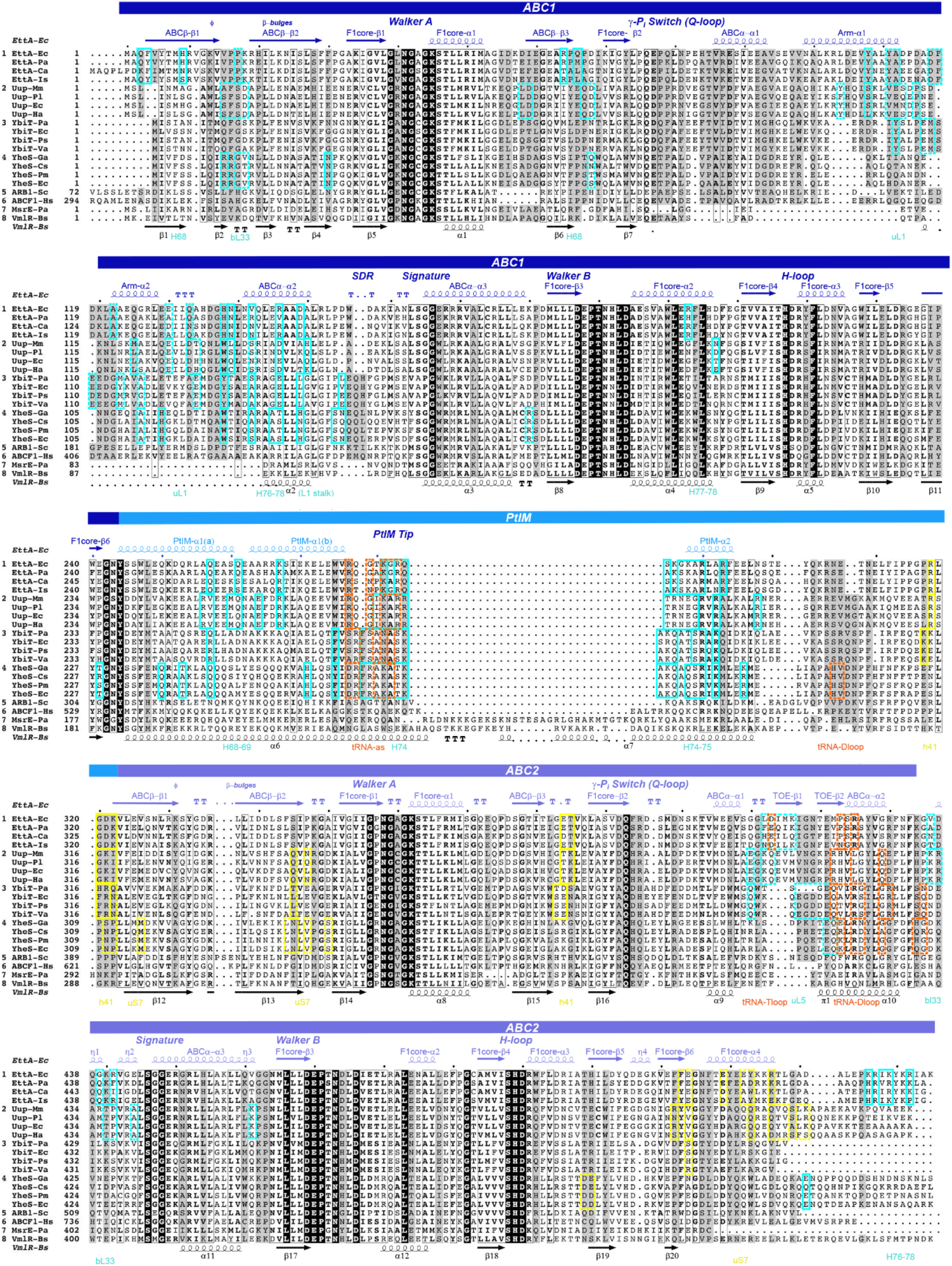
Sequence-structure schematic showing variations in ribosome-interacting residues in different ABCF family paralogs. Multiple sequence alignment of four *E. coli* ABCFs and their close homologs (*i.e.* showing ∼70% sequence similarity) from other prokaryotes. The names of the organisms have been abbreviated as: Ec *Escherichia coli*, Pa *Pseudomonas aeruginosa*, Ca *Candidatus Colwellia aromaticivoran*, Is *Idiomarina-sp.*, Mm *Morganella morganii*, Pl *Photorhabdus luminescens*, Ha *Hafnia alvei*, Ps *Photobacterium sanctipauli*, Va *Vibrio aquaticus*, Ga *Gilliamella apicola*, Cs *Candidatus Fukatsuia symbiotica*, Pm *Proteus mirabilis*, Sc *Saccharomyces cerevisiae*, Hs *Homo sapiens*, Bs *Bacillus subtilis.* Eukaryotic ABCFs for which structures have been reported along with two antibiotic resistance factors (ARE) ABCFs having longer PtIMs have been included to compare the sequence conservation and deviations from the *E. coli* ABCFs within the sub-domains. Secondary structural elements have been assigned using the ribosome bound conformation of EttA (shown in navy) and VmlR (shown in black). ABC1, PtIM and ABC2 sub-domains have been shown in navy, marine and slate blue shades respectively. Columns where the sequences remain 100% conserved are shaded in black with bold letters; while the columns that show synonymous mutations are shown in bounding black boxes with the letters in bold. Columns shaded in gray show sequence conservation within the homologs from different species but different between the paralogs. ABCF residues involved in interactions with components from 70S initiation complex have been enclosed in boxes, solid lines for rRNA and dashed lines for ribosomal proteins. Boxes in cyan are for 50S subunit, yellow for 30S subunit, and orange for tRNA.

**Figure 4.**
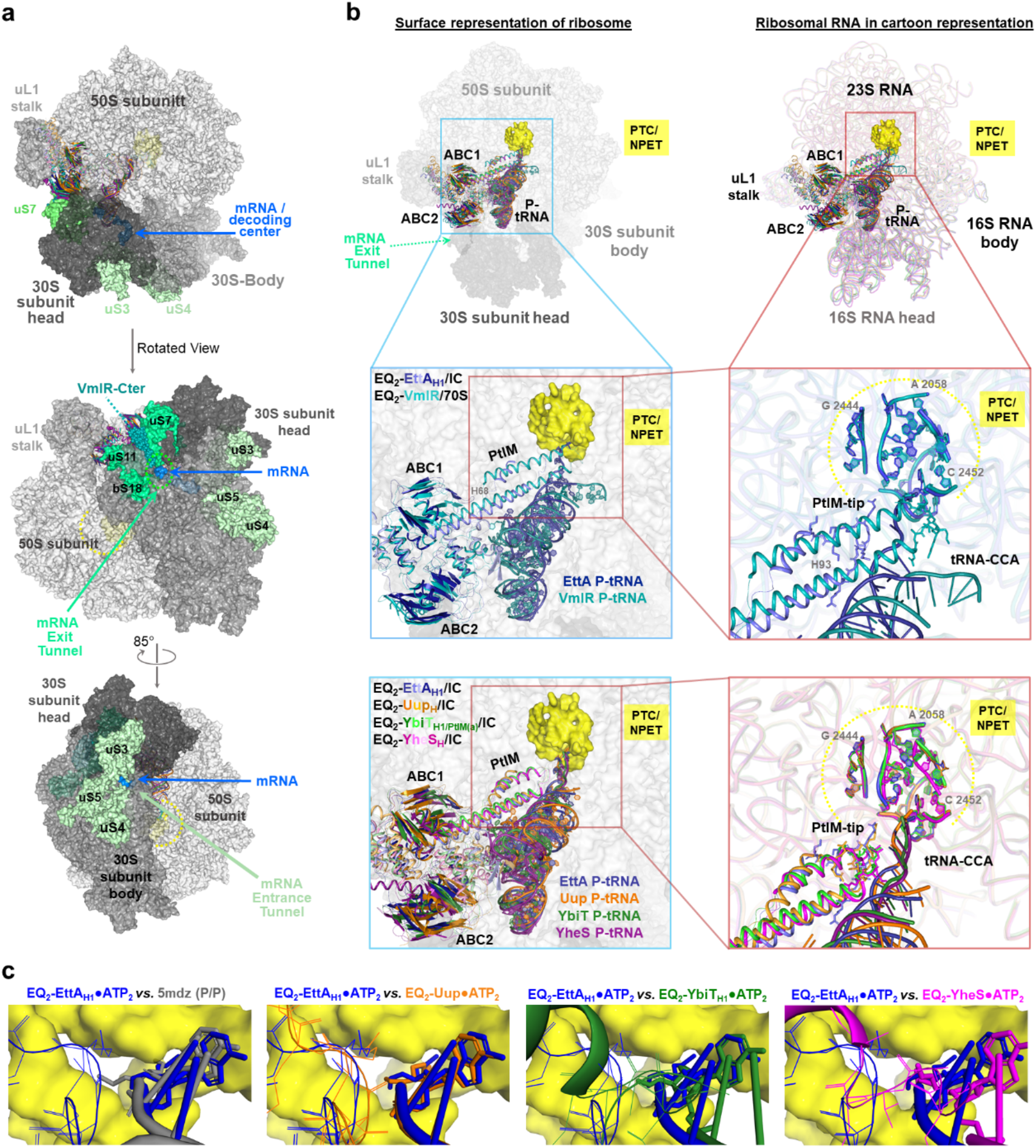
ABCF proteins bind in an equivalent geometry in the E site with the <u>P</u>-site tRNA <u>I</u>nteraction <u>M</u>otif (PtIM) making extensive contacts to the 23S rRNA and P-site tRNA. **(a)** Surface representation of four 23S rRNA-aligned *E. coli* EQ_2_-ABCF-bound 70S ICs and VmlR-bound 70S complex from *B. subtilis* (PDB ID 6ha8), showing the 30S head in dark gray, 30S body and uL1 stalk in medium gray, and 50S subunit in the lightest gray shades and the 30S subunit proteins (uS3, uS4 and uS5) lining the entry into the mRNA channel in dull green, the mRNA in the channel in marine blue, and the 30S subunit proteins (uS7, uS11, uS17) lining the exit site of mRNA channel in bright green. The C-terminal domain from VmlR is shown in teal spheres. **(b)** (Top left panel) Cartoon representation of the five complexes in **a** showing conserved positioning of PtIM in the 70S ribosome-bound complexes. Semi-transparent surface for only EQ_2_-EttA_H1_-bound 70S IC is shown with the 50S subunit in light gray and the 30S subunit in dark gray and residues lining the PTC/NPET are shown in yellow. *E. coli* ABCFs and their corresponding P-site tRNAs are color coded as: EttA in blue, Uup in orange, YbiT in green, and YheS in magenta with darkest shade for CORE subdomains and the tRNA, medium shade for PtIM, and lightest shade for ABCα and Arm helices. ARE ABCF VmlR is shown in light teal and the P-site tRNA from the complex is shown in dark teal. Top right panel shows the ribosomal 16S and 23S rRNA in cartoon representation for all the 70S ribosome complexes (color codes are same as the ABCFs). Middle left panel is the region from the inset showing zoomed in view of EttA and VmlR and the P-site tRNAs from their 70S-ribosome-bound complexes; middle right panel zooms in further showing only the PtIMs (tip of the PtIM shown in stick representation) and the terminal region of the acceptor stem of the P-site tRNA while bottom panel is the four *E. coli* ABCFs. Yellow dashes encircle the bases forming the PTC/NPET region, shown in cartoon representation on the right (middle and bottom panels). **(c)** Comparisons at the PTC between the four *E. coli* ABCF paralogs showing the PtIMs and acceptor stem of the P-site tRNA, EQ_2_-EttA_H1_ (shown in blue with the residues forming the PTC/NPET as yellow surface) is used as reference. First panel shows the comparison of the reference with the tRNA from an *E. coli* reference ribosome with a P-site tRNA in its classical configuration (PDB ID 5mdz).

The binding geometry of the ABCF proteins in the E site of the ribosome positions the C-termini of their ABC2 domain ∼45 Å from the mouth of mRNA exit channel in the 30S subunit (**Fig. 4a**). The structure of VmlR (PDB ID 6ha8)^22^, an ARE ABCF, shows that the 64-residue C-terminal extension after the end of ABC2 in that protein forms an α-helical hairpin bound on the surface of the 30S subunit in a geometry in which the turn joining the two helices in the hairpin protrudes into the mouth of mRNA exit channel (**Fig. 4a**). Therefore, the 5’ segment of the mRNA being translated by an ABCF-bound ribosome, including the 5’ untranslated region (5’-UTR) of the mRNA in a 70S IC, can potentially interact directly with the ABCF protein given its binding geometry in the E site.

### Overview of ribosome conformations in the ABCF complexes

The EQ_2_ variants of the *E. coli* ABCF proteins all stabilize the ribosome in a global conformational state, referred to a Global State 1 (GS1) or Macrostate-I^75,76^ in which the ribosomal subunits are in their “non-rotated” relative orientation and the P-site tRNA is in its “classical” configuration (**Figs. 2b, S7b, & S10**). These structures are consistent with the results of smFRET studies of their influence on the dynamics of PRE^-A^ complexes^8,25,67^. As described in detail in the ***Supplementary Discussion*** section in the ***SI*** for this paper, these structures, similar to all previously determined structures of the ATP-bound conformations of ARE proteins^8, 20–25,27,33,34^, show small counterclockwise rotations of the “body” domain of the 16S rRNA in the 30S subunit relative to the 50S subunit while remaining in the GS1 conformation. Using the angular conventions in a recent analysis of ribosome structures^77^, the body rotations range from −2.2° to −1.4°, and they are accompanied by rotations of the “head” domain of the 16S rRNA relative to its body ranging from +1.1° to +4.2° (**Figs. 2b, S7b, & S10**). These values all reside close to a dominant cluster of anticorrelated body/head rotation angles in a large ensemble of *E. coli* ribosome structures in the GS1 state^77^. The spread of the values in this analysis compared to the trend line suggests that the uncertainty in the determination of the rotation values is generally under ∼1°. In this context, the small negative rotations of the head of the 16S rRNA and small positive rotations of its body in the ABCF complex structures are likely to be statistically significant. The observed 16S rRNA body and head orientations are controlled by direct structural interactions with the bound ABCF factors (**Figs. 2f & S7g**) and possibly also by their modulation of the binding geometry of the P-site tRNA. (See below.) However, the functional importance of these small rotations is unclear.

The complexes show substantially larger variations in the orientation of the uL1 stalk in 23S rRNA in the large ribosomal subunit compared to the standard reference structure for GS1 with a P-site tRNA in the classical state^78^ (**Fig. S10**). The uL1 stalk rotates substantially inward towards the PTC compared to this reference structure, with the rotations of the stem and head of its rRNA varying from ∼9°- 17° and ∼16°- 26°, respectively, in the different ABCF complexes, compared to ∼45° and ∼48°, respectively, in the standard reference structure for Global State 2^78^ (GS2, alternatively called Macrostate-II). The specific rotation is determined by direct packing interactions between each ABCF protein and both the rRNA and uL1 protein in the uL1 stalk (∼700-1,400 Å^2^ of buried solvent-accessible surface area per component – **Figs. 2f & S7g**). The variations in the orientation of the uL1 stalk in the different ABCF complex structures are not correlated with the small rotations between the 30S and 50 S ribosomal subunits described above (**Fig. S10b**). This observation demonstrates that binding of ABCFs in the E site and their interactions with the uL1 stalk can override the coupling of its orientation to intersubunit rotation in ABCF-free 70S ribosome complexes with a deacylated P-site tRNA that spontaneously fluctuate between the GS1 and GS2 conformations^61,63,68,79,80^.

### ABCF protein orientation in the E site of the ribosome varies between paralogs due to variations in intermolecular contacts

Mapping the detailed atomic contacts in our 70S IC cryo-EM structures to a sequence-structure alignment of the *E. coli* ABCF paralogs (**Fig. 3**) shows that the specific residues mediating their binding to the ribosome are not conserved, and in many cases, they are not in homologous positions in the different ABCF factors even though the same subdomains consistently interact with the same structural components of the ribosome in all of the complexes (**Figs. 2f & S7g**). The variations in the positions of the interacting residues are attributable to the conserved subdomains within the ABC domains binding in different orientations relative to the conserved core of the 50S subunit of the ribosome in the 70S IC complexes with the different paralogs (left in **Fig. 4b** and **Fig. S11**). Following alignment of the 23S rRNA in the 50S subunit in these complexes, the orientations of the equivalent subdomains differ by 5°- 38°. (See the legend to **Fig. S11** for details.) These differences reflect significant variations in the relative orientations of the subdomains within each ABCF protein between the different paralogs. These subdomain rotations result in changes in the location and identity of the amino acids that make contacts to equivalent regions of the ribosome and the P-site (**Fig. 3**), and they likely contribute to the small differences in ribosome conformation described above (**Figs. 2b, S7b, & S10**) as well as the larger differences in P-site tRNA-binding geometry detailed below (**Figs. 4b, 5, S9, & S12-S14**).

**Figure 5.**
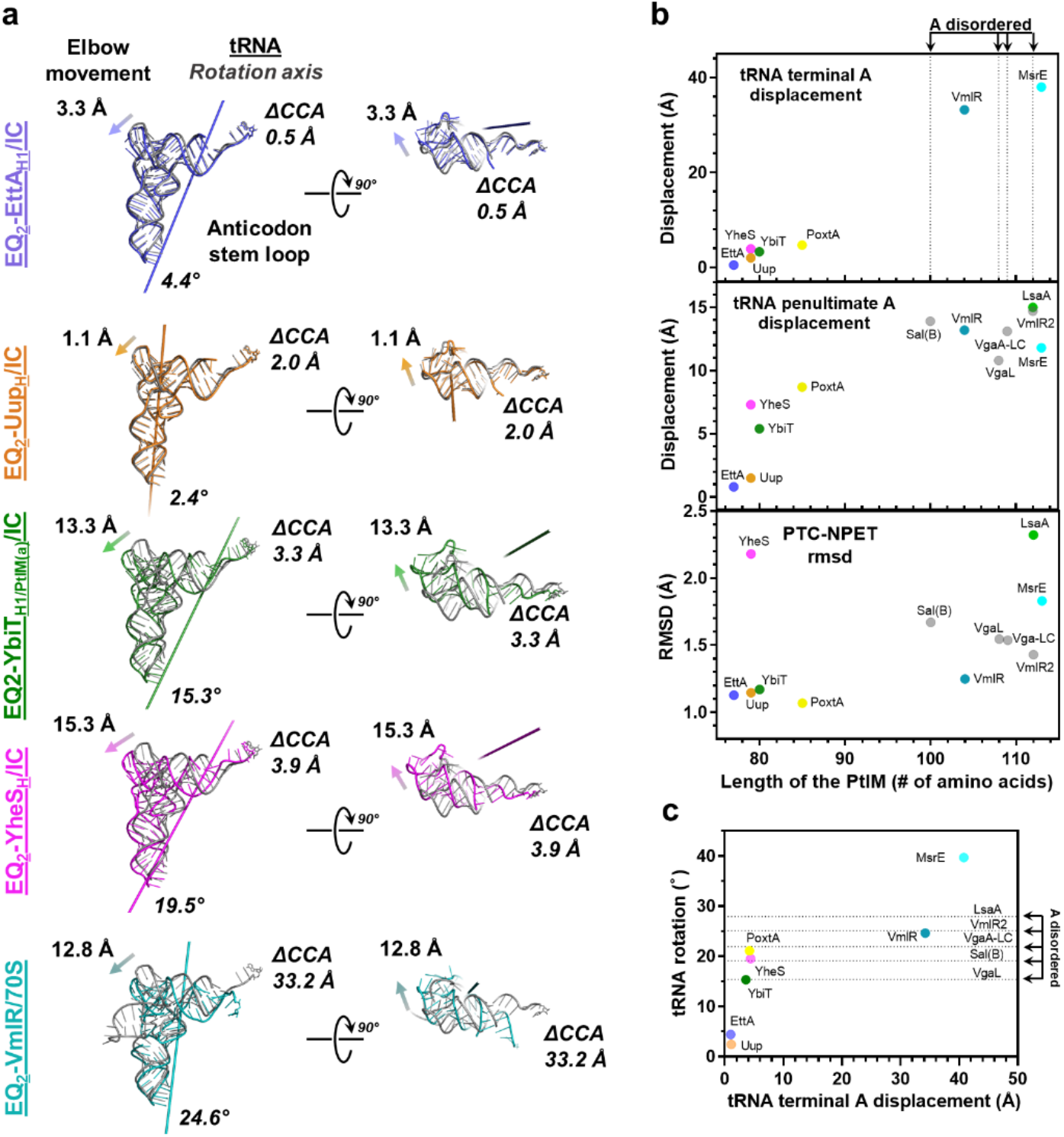
ABCF proteins control P-site tRNA binding geometry via the PtIM. **(a)** P-site tRNA from the four *E. coli* EQ_2_-ABCF (EttA in blue, Uup in orange, YbiT in green, and YheS in magenta)-bound 70S ICs, and VmlR-bound 70S (in teal), showing the rotation of the P-site tRNA relative to a P-site tRNA in its classical configuration (in gray) from the reference ribosome (PDB ID 4v9d). The axis running across the length of the tRNA represents the axis of rotation with values for the rotation mentioned at the bottom right of the axis. ΔCCA represents the shift in terminal A of the acceptor stem while the arrow at top left of each of the tRNAs shows the direction of elbow movement and value in (Å) shows its magnitude. Shift vectors for the ARE ABCF VmlR are calculated against the reference ribosome from *B. subtilis* with P-site tRNA (PDB ID 6ha1). **(b)** Graph showing the displacement of the acceptor stem of the P-site tRNA in the ABCF-bound 70S ribosome complexes (for the four *E. coli* ABCFs and the ARE ABCFs (PDB IDs: 5zlu, 7nhk, 7nhl, 7nhn, 7p48, 7p7r, 8buu and 6ha8)) relative to the length of the PtIM (number of amino acids in the PtIM). Top panel shows the displacement in (Å) for the terminal A in the acceptor stem, the dashed lines represent the ABCFs Sal(B), VgaL, VgaA-LC, and LsaA for which the terminal CCA in acceptor stem is disordered. Middle panel shows the displacement for the A before CCA relative to the P-site tRNA in reference ribosome. Lower panel shows the RMSDs of the bases at the PTC/NPET between the ABCF bound complexes relative to reference ribosome. **(c)** Graph plotted for rotation of P-tRNA (°) relative to displacement (Å) of the A in CCA. Four horizontal dashed lines are for the ARE ABCFs with terminal A unresolved.

Despite the observed variations in the relative orientations of the subdomains within the tandem ABC domains, there is tight conservation of the stereochemistry at the two ATPase active sites at their mutual interface in the ATP-bound hydrolytic conformations of all of the ABCF paralogs. (See **Fig. 7** below.) Therefore, evolution has selected for sequence changes that preserve the detailed stereochemical properties of their active sites while altering the inter-subdomain interaction geometries within the different ABCF paralogs to stabilize different intersubunit orientations and P-site tRNA configurations within 70S ribosome complexes in GS1 (**Fig. S11**). Notably, the different hydrolytic conformations of EQ_2_-EttA and EQ_2_-YbiT show significant oscillations of their tandem ABC domains within the E site of the ribosome (left and center in **Movie S1**). However, these oscillations maintain essentially the same packing geometries both in the interface between those domains as well as their interfaces with components of the ribosome, which produces coupled oscillations in the uL1 stalk in the 50S ribosomal subunit and the head of the 30S subunit. Therefore, the local packing interactions of each ABCF paralog in the E site of the ribosome remain stable during the observed rotations of their ABC domains even though those interactions vary between different paralogs.

The range of variation observed in the orientation of the ABCα subdomain relative to the CORE subdomain is substantially greater in ABC1 *vs.* ABC2 in the 70S IC complexes of the *E. coli* ABCF proteins (**Fig. S11f**). This difference is very likely attributable to restriction of the relative motion of CORE2 and ABCα2 by tight packing in their mutual interface of the polypeptide segment that follows the C-terminus of the PtIM, while no equivalent packing interaction exists in ABC1 (**Fig. S11b-e**). Studies of other ABC Superfamily proteins have established that reorientation of the ABCα subdomain relative to the CORE subdomain can play a central role in allosteric control of ATPase activity^36,42,59^ via modulation of the conformation of the γ-phosphate switch (Q-loop), the polypeptide segment connecting the two subdomains, which contains an invariant glutamine that hydrogen-bonds (H-bonds) to the γ-phosphate of ATP in the standard catalytic geometry for ABC ATPases. Restriction of movement between the ABCα2 and CORE2 subdomains in ABC2 by the C-terminus of the PtIM, which is a conserved feature in all ABCF structures determined to date, is likely an evolved feature that gives ABC1 the principal role in allosteric control of their ATPase activity. This inference is supported by structural studies of the regulation of the ATPase activity of wild-type (WT) EttA (manuscript in preparation^65^) as well as analyses of the different conformations of YbiT bound to the 70S IC. (See **Figs. 8 & S17** below).

### Conserved binding geometry of the ABCF PtIM on 23S rRNA

The binding geometry of the PtIM between the P-site tRNA and 23S rRNA in the 50S subunit is very strongly conserved in all ABCF complex structures determined to date, including the *E. coli* paralog structures presented here (**Figs. 4b-c**), all ARE structures (**Figs. 4b & S13**), and a structure of yeast Arb1 bound to the large (60S) ribosomal subunit in a ribosome recycling complex from that organism (**Fig. S14**). Despite this strong conservation in local binding geometry, the specific residues in the PtIM mediating its interactions with 23S rRNA are still not conserved (**Fig. 3**). Nonetheless, the central region in the N-terminal α-helix in the PtIM is always bound in a shallow groove formed by helix H68 and the linker between helices H69-H70 in 23S rRNA (*i.e.*, with G1839-G1840 and C1893- C1894 in H68 and U1926 in the H69-H70 linker making contacts to EQ_2_-EttA) (right in **Fig. 4b**). The tip of the PtIM, which connects its N-terminal and C-terminal α-helices, always interacts directly with the acceptor stem of the P-site tRNA while simultaneously making extensive interactions with 23S rRNA segments proximal to the PTC, including helices H74-H75 and H93 (*i.e.*, G2064 and A2435-G2437 in H74, U2076-A2077 and G2253-C2254 in H75, and C2594-U2596 and A2600 in H93 for EQ_2_-EttA). However, even though the tip of the PtIM varies in length by as much as 36 residues in different ABCF proteins, it does not penetrate into the active site of the PTC except in two of the 13 known ABCF complex structures, and even these two structures show very limited contacts between the PtIM and nucleotides in the PTC (*i.e.*, a single residue in LsaA^20^ and three in MsrE^27^). The longer tip sequences make more extensive packing interactions with the 23S rRNA segments proximal to the entrance to the PTC while producing significantly larger displacements in the acceptor stem of the P-site tRNA compared to the canonical catalytic geometry for peptide bond formation (**Figs. 5, S10a, & S12-S13** and **Table S4a**).

### ABCF proteins control the binding geometry of the P-site tRNA

While the PtIM in all of the ABCF proteins interacts with 23S rRNA in a conserved geometry and makes qualitatively similar interactions with the P-site tRNA (**Fig. 4b**), the different paralogs produce widely varying conformations and configurations of its acceptor stem (**Figs. 4b-c, 5, S10a, & S12-S13** and **Table S4a**) dependent on the sequence and length of the tip of the PtIM (**Fig. 5b**). Some ABCF proteins stabilize the acceptor stem in the proper catalytic geometry for peptide bond formation in the PTC, while others clearly disrupt this geometry, as detailed below. The P-site tRNA nonetheless remains in approximately the same region of the ribosome in all of the ABCF protein complexes as in the classical ribosome configuration in the non-rotated GS1 state (**Figs. 2, S7, S10, & S13**), and they all furthermore maintain proper codon-anticodon interactions within the P site of the 30S subunit in all of the ABCF complexes (**Fig. 6d-i**). These interactions are conserved despite varying perturbations in the acceptor-stem geometry via different rotations of the P-site tRNA around axes extending roughly from their acceptor stems to their anticodon loops (left in **Fig. 5a**). These rotations within the P site are mediated by interactions of the elbow region of the tRNA with the ABCα2 subdomain in ABC2 and in some cases also with residues in the C-terminus of the PtIM in the region linking it to the N-terminus of ABC2 (**Fig. 4b**). The solvent-accessible surface-area buried in the interface with the ABCα2 subdomain tends to be higher in the complexes with larger rotations of the P-site tRNA compared to the classical configuration^61,62,78^ (**Figs. 2f, 4b, 5, & S7g**), and larger rotations are correlated with larger displacements of the acceptor stem from the proper catalytic geometry for peptide bond formation (**Fig. 5c**).

**Figure 6.**
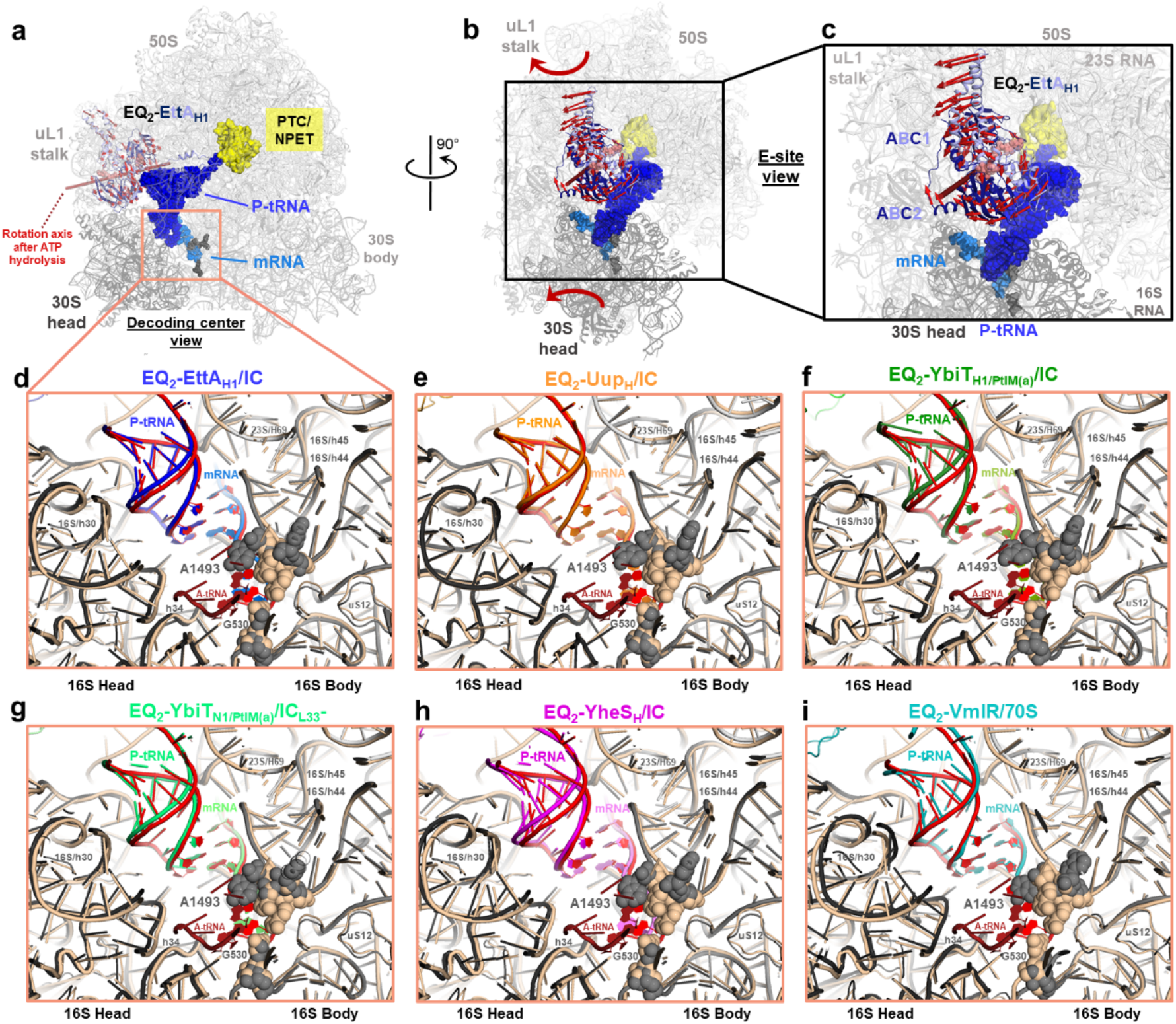
The ATP-bound conformations of the four E. coli ABCF paralogs conserve the stereochemistry of the P site anticodon/codon interactions in the 30S subunit, but ATP hydrolysis could potentially drive transient distortions. **(a)** Cartoon representation of EQ_2_- EttA_H1_/IC showing the 30S head in dark gray and the 30S body and 50S subunit in light gray, the P-site tRNA in blue and mRNA in marine blue, and the bases G530, A1492, and A1493 as dark gray spheres. EttA is shown in blue (CORE subdomain in navy and ABCα in light blue). Axis of rotation, of the ATP-bound EttA in the 70S ribosome-bound state relative to its *apo* form reported in the crystal structure (PDB ID 4fin) is shown as red line. Shift vectors between the Cα in the ABC domains, for the two EttA structures are shown as red arrows, based on a CORE alignment of the adjacent domain. **(b)** Rotated view of **a** showing the E site, curved red arrows depict the probable direction of motion of the uL1 stalk and 30S head subsequent to ATP hydrolysis and repulsion of the ABC dimer structure. **(c)** Zoomed in view of **b**. Panels **(d-i)** show a zoomed-in views of the A site of the EQ_2_-ABCF-bound 70S ribosome complexes relative to reference ribosome within a pre-translocation complex (PDB ID 7k50) with the 30S subunit in its “closed” state and the bases G530, A1492, and A1493 in the conformation where they can interact with the backbone of codon-anticodon helix. The reference ribosome is shown in beige, its P-site tRNA and mRNA segment are shown in red, and its A-site tRNA in crimson red. **(d)** EQ_2_-EttA_H1_/IC with its P-site tRNA in blue, mRNA in marine blue, 30S subunit in dark gray, 50S subunit in light gray, and bases G530, A1492 and A1493 as gray spheres. **(e)** EQ_2_-Uup_H_/IC with its P-site tRNA in dark orange, mRNA in light orange. **(f)** EQ_2_-YbiT_H1/PtIM(a)_/IC with its P-site tRNA in dark green, mRNA in bright green. **(g)** EQ_2_-YbiT_N1/PtIM(a)_/IC_L33_- with its P-site tRNA in green, mRNA in lighter shade of green. **(h)** EQ_2_-YheS_H_/IC with its P-site tRNA in magenta, mRNA in lighter shade of magenta**. (i)** EQ_2_-VmlR/70S with its P-site tRNA in teal, mRNA in light teal.

The EQ_2_ mutants of two of the *E. coli* ABCF paralogs, EttA and Uup, stabilize the acceptor stem of the P-site tRNA in a roughly canonical configuration for peptide bond formation (**Figs. 4c & 5a**), maintaining the position of its aminoacylated 3’-hydroxyl group 0.5 Å or 2.0 Å, respectively, from its position in a standard reference structure for that state (PDB id 5mdz^81^). The EQ_2_-EttA-bound and EQ_2_-Uup-bound ribosomal complexes similarly show approximate conservation both of the overall orientation of the P-site tRNA (4.4° and 2.4° rotations, respectively) and of the position of its elbow region (3.3 Å and 1.1 Å displacements, respectively) compared to reference structures (PDB ids 4v9d^78^ and 5mdz^81^). Our *E. coli* EQ_2_-EttA 70S IC and EC (PRE^-A^_Val_) structures all have extremely similar P-site tRNA configurations to each other and also to the structures of the 70S IC complexes of *Mtb*EttA^26^ in which both ATPase active sites are properly formed (**Figs. S5, S7, S9, & S10a** and **Table S4a**). The small differences in the stereochemistry of the PTC between these structures are within the effective range of uncertainty in the determination of atomic positions in structures determined at the operative resolution^82^. Notably, EttA and Uup both have a β-hairpin called the “toe” inserted in their ABCα2 subdomains (**Figs. 1 & 3**), while this structural feature is missing in the ABCF paralogs characterized to date that disrupt the catalytic geometry at the PTC. Given the role of this region of the ABCα2 subdomain in controlling the rotation of the P-site tRNA (**Figs. 2f, 4b, & S7g**), which is correlated with the magnitude of the displacement of its acceptor stem from proper catalytic geometry (**Fig. 5c**), future studies should examine whether ABCF proteins having the toe motif generally promote adoption of proper catalytic geometry at the PTC as observed for EttA and Uup.

Our inference that the ATP-bound conformations of EttA and Uup stabilize the acceptor stem of the P-site tRNA in the proper geometry for peptide bond formation is consistent with a variety of biochemical results. Di/tri-peptide synthesis assays have demonstrated that the ATP-bound conformation of EQ_2_-EttA slightly accelerates formation of the first peptide bond in a 70S IC and then traps the ribosome in that state^8,67^, while WT-EttA also alleviates ribosome stalling events at certain acidic residues^66^. WT-Uup slightly increases the yield of luciferase from *in vitro* translation reactions using purified ribosomes and translations factors^67^. Finally, the cryo-EM structure of a 70S ribosome complex with Elongation Factor P (EF-P), a smaller protein that binds in the E site to promote proper peptide bound formation in proline-rich sequences, shows an extremely similar P-site tRNA geometry to EQ_2_-EttA and EQ_2_-Uup (1.5° rotation, 0.9 Å displacement of the 3’-OH of the CCA, and 1.1 Å displacement of the elbow – **Fig. S13b**).

The EQ_2_ mutants of the other two *E. coli* ABCF paralogs, YbiT and YheS, displace the acceptor stem from its canonical geometry for peptide bond formation by modest distances of 3.3 Å and 3.9 Å, respectively, and they also produce larger changes in both the overall orientation of the P-site tRNA (15.3° and 19.5° rotations, respectively) and the position of its elbow region (13.3 Å and 15.3 Å displacements, respectively) (**Fig. 5**). The acceptor stem displacement by YheS produces an allosteric change in the conformation of the PTC and the nascent polypeptide exit tunnel (NPET) that will be described in a separate paper (manuscript in preparation^83^). The EQ_2_- YbiT 70S IC and EC structures all show extremely similar P-site tRNA configurations (**Figs. S6-S7, S10a, S12, & S15**). Di/tri-peptide synthesis assays have demonstrated that the ATP-bound conformation of YbiT-EQ_2_ inhibits formation of the first peptide bond in the 70S IC^67^, consistent with its ∼3 Å displacement of the acceptor stem of the P-site tRNA (**Fig. S15**) perturbing the geometry at the PTC sufficiently to significantly reduce the rate of peptide bond formation.

These results, in conjunction with equivalent analyses presented below of the available cryo-EM structures of ARE-bound 70S ribosome complexes, suggest that the primary function of ABCF proteins is to control the position of the acceptor stem of the P-site tRNA relative to the PTC in order to modulate the rate of peptide bond formation and the geometry of nascent polypeptide interactions with the PTC/NPET. We hypothesize that some ABCF factors, including EttA and Uup, stabilize the acceptor stem of formylmethionylated- or peptidylated-tRNAs in a catalysis-competent conformation, while others, including YbiT, YheS, and the AREs of known structure, push it into a catalytically incompetent conformation. The analyses in this paper suggest several possible functions for the latter activity including resolutions of clogs in the PTC/NPET when antibiotics or troublesome protein sequences produce adventitious stereochemical interactions inhibiting translation.

### PtIM length correlates with P-site tRNA acceptor stem displacement but not PTC conformation

We compared the positions and orientations of the P-site tRNA in our *E. coli* ABCF-bound 70S ribosome complexes to those observed in the eight ARE-bound 70S ribosome complex structures that have been deposited in the Protein Data Bank (center of **Fig. 4b** and **Figs. 5, S10a, & S13b**). While the PtIMs of the AREs interact with the same regions of both the 23S rRNA in the 50S subunit and the P-site tRNA as the *E. coli* ABCFs, they uniformly produce larger displacements of the terminal 3’-hydroxyl group in the acceptor stem of the P-site tRNA ranging from ∼5-38 Å, and they also produce larger shifts in the position of its elbow region ranging from ∼11-16 Å, as well as larger rotations in its orientation ranging from 21°- 30°. In several ARE-bound 70S ribosome complex structures, the terminal 3’-hydroxyl group in the acceptor stem is disordered, but the penultimate A in the acceptor stem in these complexes shows ∼13 Å displacements that are similar in magnitude to the movement of the equivalent nucleotide in the ARE-bound complexes showing the largest displacements of the terminal 3’-hydroxyl group in the acceptor stem (**Fig. 5b**). These large displacements of the acceptor stem will clearly disrupt the catalytic geometry at the PTC, which depends on the exact location of the terminal 3’-ribose group in the acceptor stem, in addition to directly pulling any nascent polypeptide attached to the P-site tRNA out of the PTC/NPET^21,34^.

The number of residues in the tip of the PtIM, which determines its overall length, clearly correlates with the magnitude of the displacement of the terminal 3’-hydroxyl and the penultimate A in the acceptor stem of the P-site tRNA in the ABCF-bound 70S ribosome complexes (upper panels in **Fig. 5b**). Notably, the size of the PtIM does not correlate with changes in the conformation of the PTC/NPET as assessed based on the root-mean-square deviation (RMSD) of the backbone phosphates of the 23S rRNA nucleotides forming these features (bottom panel in **Fig. 5b**). The lack of a consistent influence of the PtIM on PTC/NPET conformation is explained by the lack of any direct contacts between residues in the PtIM and nucleotides in the PTC/NPET in any complex structure except for the minor contacts made by LsaA^20^ and MsrE^27^ mentioned above. These observations support precise control of the conformation and position of the acceptor stem of the P-site tRNA being the dominant function of ABCF proteins, and they suggest that displacement of the nascent polypeptide chain attached to the acceptor stem is the major mechanism by which the AREs mediate resistance to antibiotics binding to the PTC/NPET^21,34^.

### Analysis of forces likely generated following ATP hydrolysis by the ABCFs

The availability of a nucleotide-free crystal structure of *E. coli* EttA^8^ enables modeling of the approximate trajectory of the conformational change that will take place concomitant with ATP hydrolysis when EttA is bound to 70S ribosomes (**Fig. 6a-c**). Aligning the ATP-binding CORE subdomain of each ABC domain in the nucleotide-free structure to the corresponding domain in the ATP-bound EQ_2_- EttA_H1_/IC structure shows the approximate direction of motion of the other ABC domain following ATP hydrolysis. Displaying the displacement vectors inferred in this manner suggests that hydrolysis likely generates forces that will tend to push the uL1 stalk, the P-site tRNA, and the head of the 30S subunit around an axis extending roughly from the PTC outward through the center of the E site (**Fig. 6b-c**). The magnitudes of the actual displacements in the different components of the ribosome and the P-site tRNA will depend on the relative strength of the restoring force opposing each motion, which is not clearly established. However, given the many observations showing that the uL1 stalk is flexibly attached to the rest of the 50S subunit^75,76,81,84^, the restoring force opposing its motion is likely to be weak, suggesting that the main movement in the ribosome following ATP hydrolysis is likely to be in the position of the uL1 stalk.

Nonetheless, the forces generated on the P-site tRNA and the 30S head subsequent to ATP hydrolysis (**Fig. 6b-c**) could potentially propagate to the anticodon loop of the P-site tRNA and interacting mRNA codon in the interface between the head and body of the 30S subunits. This codon-anticodon interaction and the conformation of the adjacent decoding center are approximately canonical in all of our cryo-EM structures with *E. coli* ABCF proteins compared to a standard pre-translocation reference structure^85^ (PDB id 7k50) with both P-site and A-site tRNAs (**Fig. 6d**), and the local stereochemistry in this region is maintained during the observed rotations of the ATP-bound pre-hydrolysis conformations of both EQ_2_-EttA and EQ_2_-YbiT in the E site of the ribosome because the P-site tRNA and the ‘platform’ region of the 30S subunit both rotate together with the 30S head (**Movie S1**). These observations demonstrate that, even when they occur, rotations of the P-site tRNA and 30S head do not necessarily perturb tRNA-mRNA interactions. While the force analysis above suggests ATP hydrolysis by ABCF proteins could potentially dynamically modulate codon-anticodon interactions and the conformation of the interacting ribosomal segments at the interface between the head and body of the 30S subunit, future studies will be required to determine whether there are any functionally significant effects of this kind associated with the ATPase cycles of ribosome-bound ABCF proteins.

### Canonical ATP encapsulation even by non-canonical ABC Signature Sequences

All of the 70S ribosome complexes we reconstructed for the EQ_2_ variants of the four *E. coli* ABCF proteins show their tandem ABC domains forming ATP-sandwich complexes with two unhydrolyzed ATP molecules (**Fig. S8a**) bound between the Walker A/B motifs in the ATP-binding Core of one ABC domain and the Signature Sequence in the ABCα subdomain of the other ABC domain (**Fig. 7**). Despite deviations from the canonical ABC Superfamily LSGGQ Signature Sequence at minimally one position in both domains in every *E. coli* ABCF protein (**Fig. 3**), their atomic interactions with the β/γ-phosphates of the bound ATP molecules are nonetheless essentially canonical^38,59^ in all of their composite ATP-binding sites except for those in YbiT (**Fig. 7c**), which show substantial variations between the different conformational classes reconstructed for its 70S ribosome complexes (described below). As analyzed in detail in the ***Supplementary Discussion*** section in the ***SI*** for this manuscript, the atomic interactions with ATP observed in our cryo-EM structures of *E. coli* EQ_2_-EttA, EQ_2_-Uup, and EQ_2_-YheS (**Fig. 7c**) suggest that all of their ATP-binding sites are likely to be catalytically competent and captured in a functional pre-hydrolysis conformation in the ATP-bound 70S ribosome complexes we have reconstructed. A key observation supporting this inference is they all of these sites show canonical values (**Table S4b**), for two fiducial distances that we call Separation and Slide, which describe the interaction geometry of the Walker A motif and Signature Sequence motifs forming each composite ATP-binding site^38,59^ (**Fig. 7a-b**). However, enzymological studies are needed to verify this inference and to establish the influence of the non-canonical residues in their Signature Sequences on their catalytic rate constants and the details of the chemical and conformational dynamics accompanying ATP hydrolysis.

**Figure 7.**
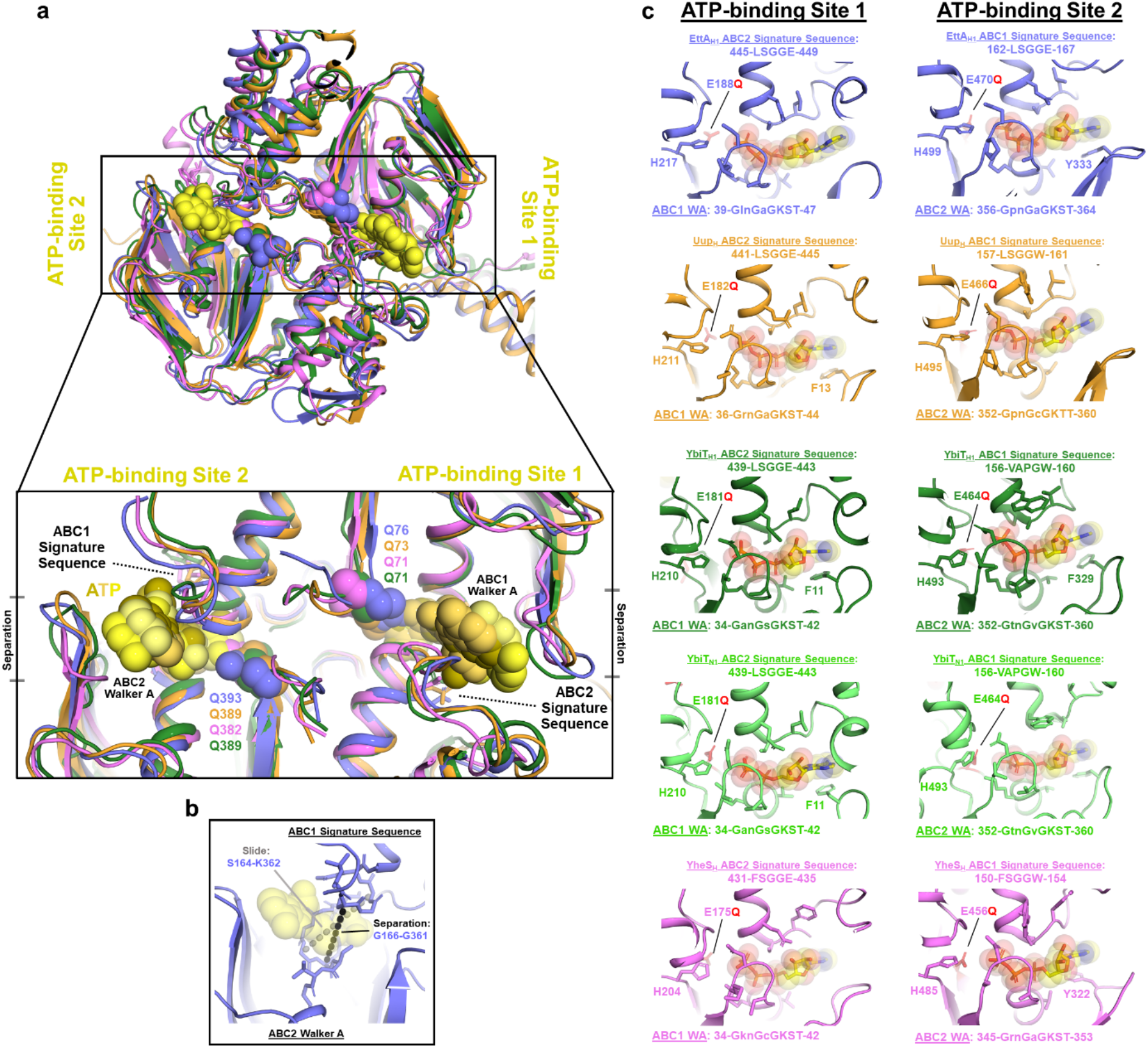
Stereochemistry of ATP binding in the composite active sites in the E. coli ABCF paralogs. **(a)** Cartoon representation of CORE1 aligned *E. coli* EQ_2_-ABCFs in their 70S IC-bound state. EttA is colored blue, Uup orange, YbiT green, and YheS magenta. ATP at the two ATP- binding sites on the ABCFs (shown in different shades of yellow) and catalytic glutamate from the two γ-Pi switch loops is shown as spheres. **(b)** ATP-binding site 2 in EttA showing the separation distance in black and slide distance in gray. **(c)** Zoomed in view of the ATP-binding sites 1 and 2 from *E. coli* EQ_2_-ABCF-bound 70S ICs.

### 70S ribosome complexes of EQ_2_-YbiT show both catalytic and non-catalytic ATPase active site conformations

Local three-dimensional classification of the particles in our EQ_2_-YbiT 70S IC sample using a mask encompassing YbiT, the P-site tRNA and the uL1 stalk yielded five conformational classes, which we have labeled according to conformational features as described above (**Figs. 2, 8, S3, S6, S12, & S15-S16** and **Tables S2 & S4**). The two Hydrolytic conformations (*i.e.*, EQ_2_-YbiT_H1/PtIM(a)_/IC and EQ_2_-YbiT_H2/PtIM(a)_/IC together representing ∼37% of the population. They are extremely similar except for a rigid-body rotation of the tandem ABC domains in the E site coupled to a small movement of the uL1 stalk mostly perpendicular to the plane of the 50S/30S interface and a small rotation of the head of the 30S subunit compared to its body that maintains the codon-anticodon interactions of the P-site tRNA in the 30S subunit (center in **Movie S1**). The two Non-hydrolytic conformations (*i.e.*, EQ_2_-YbiT_N1/PtIM(a)_/IC_L33_- and EQ_2_- YbiT_N2/PtIM(b)_/IC, representing ∼33% and 14% of the population, respectively) are also similar except for a rotation of ABC2 relative to CORE1 coupled to a modest shift in the PtIM binding geometry (**Fig. 8a**) that preserves the stereochemistry of the interaction between the acceptor stem of the P-site tRNA and the PTC (**Fig. S15a**). The Intermediate conformation (i.e., EQ_2_- YbiT_1/PtIM(b)_/IC, representing ∼16% of the population) is similar to the Hydrolytic conformations but has the alternative binding geometry of the PtIM and a small rotation of the ABCα1 subdomain in roughly the same direction as the larger rotations in that domain in the Non-hydrolytic conformations (that are explained in detail below). Notably, the stereochemical interactions of the P-site tRNA with the PTC are essentially the same in all our EQ_2_-YbiT structures (**Fig. S15b**) despite the repositioning of the PtIM (**Fig. 8a**).

**Figure 8.**
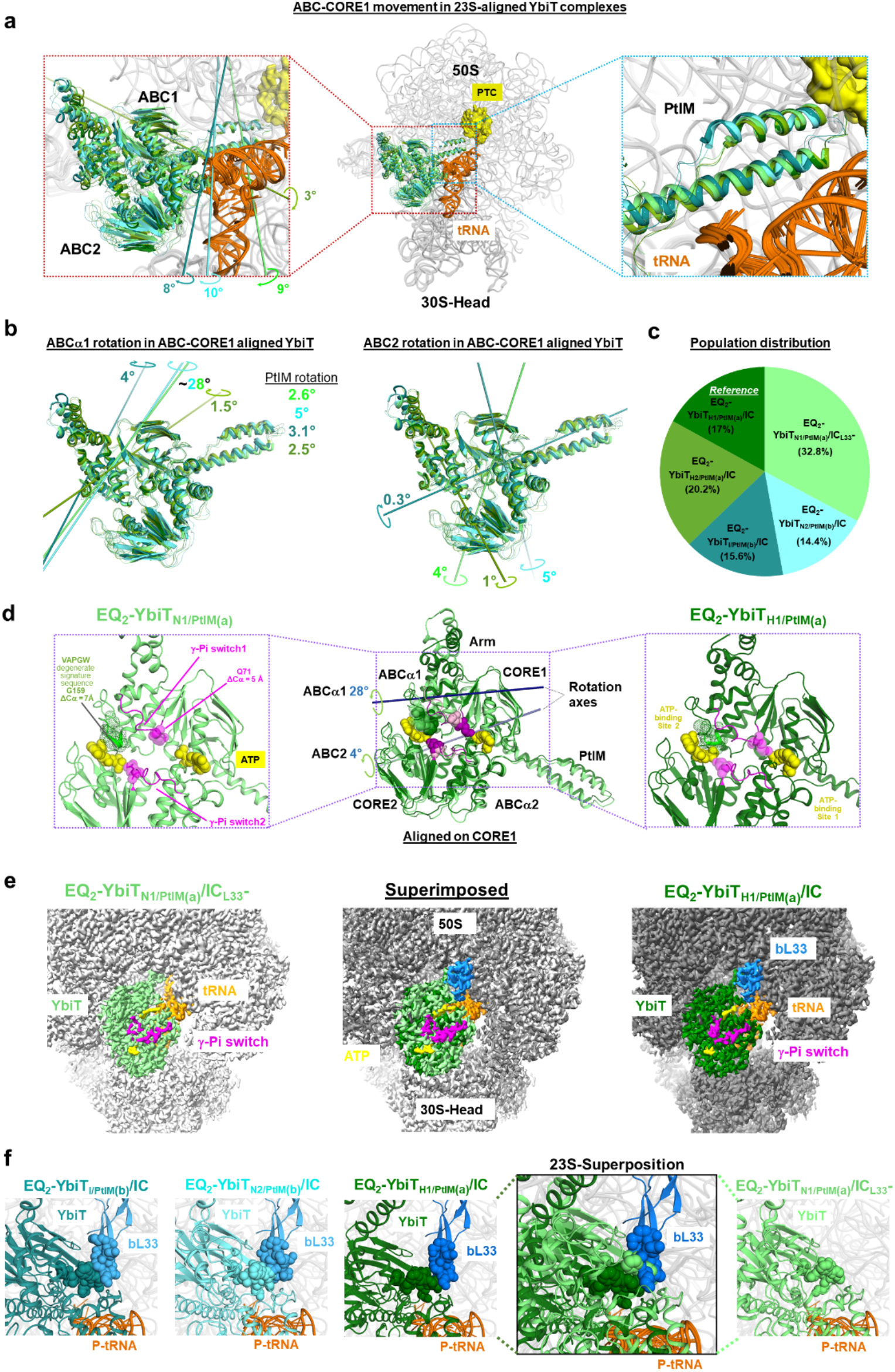
YbiT conformation and likely ATPase activity are linked to the presence/absence of the bL33 protein on the 50S subunit. **(a-f)** Comparison of the five classes resolved for EQ_2_-YbiT- bound 70S ICs showing the subdomain orientation/rotations in YbiT relative to each other. Population distribution of the five classes shown in panel **c**, the conformation of YbiT in each is designated as H for Hydrolytic and N for Non-hydrolytic, with the two different PtIM positions designated as “(a)” and “(b)” and absence of density for the ribosomal protein bL33 designated as L33**^-^**. The five complexes aligned on their 23S rRNA are shown in panel **a** (middle panel), with 16S rRNA (dark gray) and 23S rRNA (light gray) and tRNA (orange) from the ICs in cartoon representation along with YbiT (different shades of green). Positioning of the PtIMs in YbiT from each of the five complexes is shown in right panel and the left panel shows the relative rotations of the CORE1 domain in each relative to the reference structure EQ_2_-YbiT_H1/PtIM(a)_/IC shown in dark green. The lines running through the domain depict their axes of rotation (colors for each correspond to that of YbiT). **(b)** Sub-domain rotations in the 70S IC-bound YbiT relative to the reference structure, the axes on left show the rotation of ABCα1 on an ABC-CORE1 alignment, PtIM rotations are mentioned next to the PtIM, color coded for each YbiT; rotation of the ABC2 domains are shown in the right panel. **(c)** Population distribution of the five YbiT-bound 70S ICs. **(d)** Cartoon representation of the ABC-CORE1 aligned YbiT in the Hydrolytic and Non-hydrolytic conformations. The axes of rotation of the ABCα1 (dark blue line) and ABC2 (light blue) in the Non-hydrolytic state relative to the reference Hydrolytic state. The ATP molecules (yellow), the glutamate (magenta) from γ-Pi switch, and the degenerate signature sequence in ABC1 (VAPGW) are shown as spheres. Left and right panels show zoomed in view of the inset showing individual YbiT. ΔCα value show the linear displacement of G159 in the ABC1 signature sequence and the Q71 in ABC1 γ-Pi switch from the Non-hydrolytic conformation of YbiT, relative to the Hydrolytic conformation. **(e)** Density map for the two YbiT-bound 70S ICs, YbiT Non-hydrolytic (dull green) shows no density for bL33 has the γ-Pi switch loop (magenta) in ABC1 retracted away from the ATP, and YbiT Hydrolytic (dark green)-bound 70S IC showing density for bL33 (marine blue) has the γ-Pi switch loop in ABC1 contacting ATP (yellow) at ATP-binding site 1. **(f)** bL33 and YbiT in the different EQ_2_-YbiT-bound 70S ICs, inset shows the superposition of YbiT in the Non-hydrolytic and Hydrolytic conformations with residues from the Non-hydrolytic conformation clashing with bL33 present in the Hydrolytic conformation.

Applying the same classification procedure to the 70S EC (PRE^-A^_Val_) dataset yielded a single high-resolution class (*i.e.*, EQ_2_-YbiT_N1/PtIM(a)_/EC in **Fig. S6** and **Table S2**) that is extremely similar in conformation to EQ_2_-YbiT_N1/PtIM(a)_/IC_L33_-, which is the largest class in 70S IC dataset. The EC dataset, which is smaller and lower in overall quality than the 70S IC dataset, yields an aggregated intermediate resolution class (before separating the single high-resolution class) that shows diffuse density roughly matching the full ensemble of high-resolution classes reconstructed from the 70S IC sample. The proportion of EQ_2_-YbiT_N1/PtIM(a)_ particles is similar in the aggregated class in the EC dataset and the ensemble of classes in the 70S IC dataset, consistent with the YbiT having similar interaction properties with both the IC and EC.

A detailed analysis of the active-site sequence features and ATP interactions observed in the YbiT_H_ conformations (**Fig. 7c**) is presented in the ***Supplementary Discussion*** section in the ***SI*** for this paper. In brief, like the other *E. coli* ABCF proteins, both YbiT_H_ structures show essentially canonical contacts^38,59^ to β/γ-phosphates of the ATP molecule bound in ATP-binding Site 1 (*i.e.*, the composite site formed between the Walker A/B motifs in CORE1 in ABC1 and the Signature Sequence in the ABCα2 subdomain in ABC2), which suggests this site is likely to be catalytically competent. These structures show nearly, but not entirely, canonical contacts to β/γ-phosphates of the ATP in ATP-binding Site 2 (*i.e.*, the composite site formed between the Walker A/B motifs in CORE2 in ABC2 and the Signature Sequence in the ABCα1 subdomain in ABC1). Because of the substitution of the canonical LSGGQ Signature Sequence with the strongly degenerate sequence VAPGW (**Figs. 3 & 7c**), Site 2 in YbiT lacks a single conserved H-bond to the to the γ-phosphate of ATP, which is made by the otherwise invariant serine residue in the Signature Sequence in the standard catalytic geometry for ABC ATPases^38,59^. This perturbation seems likely to alter the kinetic properties of Site 2 compared to canonical ATPase active sites, although it could still potentially be catalytically competent, an issue that should be addressed in future experiments.

The Separation and Slide distances (**Fig. 7a-b**) observed at both composite ATP-binding sites in the YbiT_H_ structures (**Fig. 7c & Table S4b**) are consistent with these inferences and support the EQ_2_ mutations stabilizing a physiological pre-hydrolysis conformation of YbiT in the 37% of the population in the structures we have designated as hydrolytic (*i.e.*, EQ_2_-YbiT_H1/PtIM(a)_/IC and EQ_2_- YbiT_H2/PtIM(a)_/IC). While these structures show somewhat larger variations in both the Separation and Slide distances compared to our structures of the EQ_2_ mutants of the other *E. coli* ABCF paralogs, both distances in both composite ATP-binding sites are within 1 Å of the distances observed at the equivalent ATP-binding site in the other paralogs (**Fig. 7c & Table S4b**). These variations could reflect altered kinetic properties or potentially differences in the details of the mechanochemical reaction cycle in YbiT compared to the other ABCF paralogs, but they are nonetheless consistent with Site 1 and possibly also Site 2 being catalytically active.

In contrast, the Separation and Slide distances (**Fig. 7a-b**) observed at Site 2 show larger variations in the YbiT_N_ structures (**Fig. 7c & Table S4b**), and these variations together with additional structural analyses (**Figs. 8 & S16**) support both Site 1 and Site 2 being in catalytically incompetent conformations in the 47% of the population we have designated as non-hydrolytic (*i.e.*, EQ_2_-YbiT_N1/PtIM(a)_/IC_L33_- and EQ_2_-YbiT_N2/PtIM(b)_/IC). The two composite ATP-binding sites have different structural perturbations in the Non-hydrolytic conformations relative to the canonical catalytic configuration for ABC Superfamily ATPases, but the perturbations in both sites are driven by reorientation of the ABCα1 subdomain in ABC1, as explained below.

The YbiT_N_ structures have Separation and Slide distances at Site 1 within 1 Å of those observed in our structures of the EQ_2_ mutants of the other *E. coli* ABCF paralogs (**Table S4b**), indicating essentially proper alignment of the Walker A/B and Signature Sequence motifs encapsulating the phosphate groups of the ATP molecule bound to this composite site (**Figs. 7c, 8d, & S17a-b**). However, the Separation distance is up to 2 Å lower and the Slide distance is up to 5 Å higher at Site 2 in the YbiT_N_ structures compared to the other paralogs (**Table S4b**), indicating a major shift in the configuration of the Walker A/B and Signature Sequence motifs interacting with the ATP molecule bound at this composite site (**Figs. 7c, 8d, & S17a-b**). The larger Slide distance reflects a lateral movement of the degenerate Signature Sequence in the ABCα1 subdomain of ABC1 (in the orientation in the images in this paper) driven by an ∼28° rotation of that subdomain (**Fig. 8a-b**). The lateral movement of the Signature Sequence in the ABCα1 subdomain disrupts all of its standard atomic contacts to the γ-phosphate group of the ATP molecule bound to the Walker A/B motifs in CORE2 in ABC2, resulting in solvent exposure of the β/γ-phosphate groups in that ATP molecule (**Figs. 7c & 8d**). This shift in active-site configuration, which is likely to be promoted by the strong deviations in the ABC Superfamily Signature Sequence in the ABCα1 subdomain (VAPGW *vs.* LSGGQ), is overwhelmingly likely to render Site 2 catalytically inactive in the YbiT_N_ conformations.

The 28° degree rotation of the ABCα1 subdomain that drives the lateral movement of its Signature Sequence at Site 2 in the YbiT_N_ conformations (**Figs. 8d & S17a-b**) simultaneously drives a conformational change that is likely to also render Site 1 catalytically inactive (**Fig. 8a-b**). Because the ABCα1 subdomain is connected to the ATP-binding core of ABC1 (CORE1) by the γ-phosphate switch (Q-loop), the large rotation of the ABCα1 subdomain directly withdraws the invariant glutamine (Q71) in the γ-phosphate switch from ATP-binding Site 1 (**Fig. 8d**). The ∼5 Å shift in the position of Q71 driven by the rotation of the ABCα1 subdomain disrupts the conserved H-bond between the amide group of its sidechain and the γ-phosphate group of ATP in the canonical pre-hydrolysis conformation of EQ_2_ mutants of ABC Superfamily ATPases^44,46,59^, including our structures of EttA, Uup, and YheS (**Fig. 7c**). This disruption of a highly conserved active-site H-bonding interaction to the high-energy leaving group in ATP is very likely to stop ATP hydrolysis at Site 1. Therefore, the observed ∼28° rotation of the ABCα1 subdomain in ABC1 likely blocks ATP hydrolysis simultaneously at both Site 1 and Site 2 in the YbiT_N_ conformations.

Both ATP-binding sites in YbiT also seem likely to be catalytically inactive in the Intermediate (YbiT_I_) conformation, which shows smaller coupled shifts in the positions of the degenerate Signature Sequence in the ABCα1 subdomain and the conserved glutamine in the γ- phosphate switch in ABC1 (**Figs. S17a-b**). These shifts are driven by a qualitatively similar but much smaller rotation of the ABCα1 subdomain in the YbiT_I_ *vs.* YbiT_N_ conformations (4° vs. 28°— **Fig. 8b**). Details are described in the ***Supplementary Discussion*** in the ***SI*** for this paper.

In summary, the different conformations of EQ_2_-YbiT bound to 70S ribosome complexes show coupled changes in the configuration and likely catalytic activity of both of its ATP-binding sites driven simultaneously by reorientation of its ABCα1 subdomain. Movement of the ABCα subdomain has been shown to play an important role in allosteric regulation of other ABC Superfamily ATPases^5,59^, and based on the results presented here and a cryo-EM structure we have determined for wild-type EttA (manuscript in preparation^65^), ABCα1 is likely to play a central role in the functional control of 70S ribosome activity by ABCF family proteins.

The reconstruction of multiple YbiT conformations from a single 70S ribosome complex sample implies the different ribosome-bound conformations have relatively similar free energies. This behavior is likely to reflect differences in the inherent structural/energetic properties of the active sites in YbiT, which show more deviations from the canonical sequence features in ABC Superfamily ATPases than other ABCF proteins. The most significant deviations are in the Signature Sequence in the ABCα1 subdomain in ABC1, which shows the largest functional conformational perturbations (**Figs. 3 & 7c**). The distribution of conformations observed at the ATPase active sites in YbiT could potentially be influenced by energetic perturbations caused by the EQ_2_ mutations in the catalytic glutamate residue, although all other EQ_2_-ABCF•70S ribosome complex structures determined to date show exclusively hydrolytically active ATP active-site conformations that are very similar in all of the paralogs. Thermal motion will inevitably result in enzymes exploring catalytically inactive conformations locally in their active sites during the approach to the optimal stereochemistry for catalysis^86^. These conformations will influence the catalytic rate constant, and they could also represent important components of the mechanochemical reaction mechanism. The alternative configurations of the ATPase active sites observed in our YbiT ribosome complexes are all likely to be explored during its physiological ATPase reaction cycle, and, as explained below, they seem likely to play a role in its physiological function, although their relative populations in our samples could potentially be influenced by the presence of the EQ_2_ mutations. Following attainment of the catalytic conformation, ATP hydrolysis will lead to an opening of the mechanical clamp formed by the tandem ABC domains (*e.g.*, as illustrated in **Fig. 6a-c**), likely driven by local electrostatic forces in the active site generated during phosphodiester bond cleavage^51–53^, which will eject the ABCF protein from the ribosome (**Fig. 9**).

**Figure 9.**
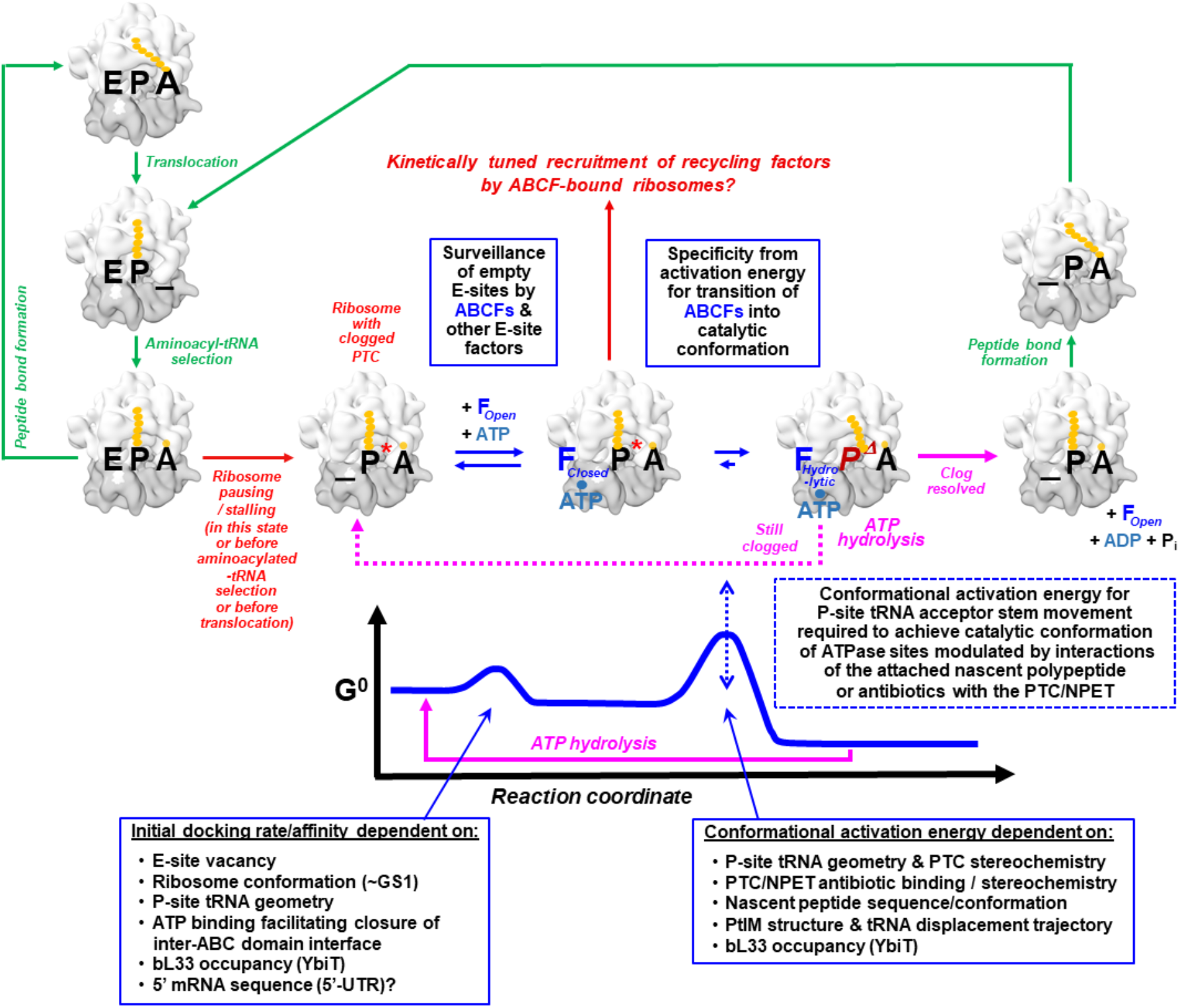
Schematic showing hypothesized ABCF mechanism-of-action based on analysis of available cryo-EM structures of paralogs bound to 70S ribosome complexes. First panel from left to right shows the steps in a normal elongating ribosome (top to bottom), E, P and A represent the tRNA, “**_**” represents an empty tRNA-binding site, and yellow spheres represent amino-acids in the nascent polypeptide. Green arrows show normal progression of one state into another and red arrows indicate ribosome pause/stall which could lead to states with empty E-site on the ribosome as shown in the second panel. ABCFs which are E-site binding factor can bind to ribosomes with an empty E-site in an energy dependent manner. Initial ATP bound ABCF undergoes a conformational change to attain a hydrolytic/catalytic conformation leading to release of energy (graph below shows the free energy landscape upon ABCF binding to the ribosome and its release post ATP hydrolysis). P* indicates a P-site tRNA in a stalled ribosome complex and *P^Δ^* the P-site tRNA that undergoes a change in configuration upon ABCF binding.

### Conformational changes in YbiT are coupled to the presence/absence of bL33

Clear and well-resolved density is observed for protein bL33 in the 50S ribosomal subunit in all of our *E. coli* ABCF-70S reconstructions (**Fig. S17a**) except for the non-hydrolytic EQ_2_-YbiT_N1/PtIM(a)_/IC_L33_- conformational class reconstructed from our 70S IC sample (**Fig. 8e**), which has the highest population among the five classes reconstructed from that sample, and the single high-resolution class reconstructed from our 70S EC sample (EQ_2_-YbiT_N1/PtIM(a)_/EC_L33_- in **Fig. S17a**). As indicated above, these classes contain a similar proportion of all YbiT-bound ribosomes in both samples, and the refined structures show YbiT in extremely similar Non-hydrolytic conformations. Note that all of our cryo-EM density maps and structures for *E. coli* ABCF-70S complexes were produced using the same local three-dimensional classification procedure employing a mask constructed from the ABCF protein, P-site tRNA, and uL1 stalk but not explicitly including protein bL33 (See Methods). Therefore, formation of a stable complex with ribosomes lacking bL33 is very likely to be a specific biochemical property of YbiT in the N1/PtIM(a) conformation.

Protein bL33 is known to be sub-stoichiometric in some *E. coli* ribosome preparations and also to exchange from ribosomes rapidly *in vivo* but not *in vitro*^87^. These observations suggest that YbiT_N1/PtIM(a)_ could have either selected ribosomes in our preparation already lacking bL33 or alternatively displaced bL33 during binding. To evaluate the first possibility, we performed local three-dimensional classification without alignment^74^ using a mask encompassing just bL33 on ribosomes harboring a P-site tRNA but without a bound ABCF protein (**Fig. S4**); these ribosomes came from the same preparation used to prepare the 70S IC sample that produced our EQ_2_-YbiT complex structures. This procedure yields equal populations of 70S ribosomes with and without density for bL33, which both yield 3.3 Å cryo-EM density maps (**Figs. S4 & S17b** and **Table S3**). Taking into account the other 70S ribosome classes reconstructed from the same sample, the 70S ribosome class harboring a P-site tRNA without density for bL33 corresponds to ∼26% of all 70S ribosomes in that sample. Using the same bL33 mask for three-dimensional classification of the particles in our IC structures with the EQ_2_ mutants of EttA, Uup, and YheS does not yield any evidence of ribosome missing bL33 in those complexes (data not shown). The failure to detect a 70S-ribosome-bound class without bL33 density for the EQ_2_ mutant of any *E. coli* ABCF protein other than YbiT suggests that their observed packing interactions with bL33 (**Figs. 2f, S7g, & S17a**) contribute significantly to their ribosome binding affinity. Notably, the other conformations of YbiT all make similar packing interactions with bL33 as EttA, Uup, and YheS (**Figs. S16c & S17a**), while the N1/PtIM(a) conformation exclusively positions the domains of YbiT so that the ATP-interacting loop in the N-terminal antiparallel β-sheet in CORE1 would have severe steric overlap with residues connecting the two β-hairpins in bL33 (**Fig. 8e**).

Based on these observations, it is possible that docking of YbiT in the ATP-bound N1/PtIM(a) conformation could directly displace bL33 from ribosomes before a spontaneous transition into the hydrolytic H1/2 conformations induces ATP-hydrolysis and YbiT release. However, the observation of a significant population without bL33 in our ribosome preparation indicates it is also possible that ATP-bound YbiT_N1/PtIM(a)_ docks preferentially to ribosomes lacking bL33 rather than displacing it. In this case, YbiT could sequester ribosomes in an inactive state until bL33 binding induces a transition into the YbiT_H1/2_ conformations to induce ATP hydrolysis and YbiT release. A three-dimensional variability analysis on combined particles from the YbiT_H2/PtIM(a)_ and YbiT_N1/PtIM(a)_ classes provides a graphical illustration of the coupling between bL33 binding and the conformational transition between Non-hydrolytic and Hydrolytic conformations of ATP-bound EQ_2_-YbiT mediated by the rotation/shift in its ABCα1 subdomain (right in **Movie S1**). Future studies will be need to distinguish between the divergent mechanistic hypotheses for the coupling between bL33 binding and YbiT binding to ribosomes. In either case, there would be an important functional role for the Non-hydrolytic conformation of YbiT observed in our cryo-EM reconstructions, which seems likely to be related to the sequence variations in its strongly degenerate Signature Sequence compared to other ABC ATPases^38,46^.

### Summary and conclusions

Comparing the 11 cryo-EM structures of 70S ribosome complexes of the four *E. coli* ABCF proteins presented in this paper to each other and to 13 previously determined ABCF complex structures establishes general structural and mechanistic principles underlying the function of the proteins in this pervasively distributed family of translation factors. These principles rationalize the great evolutionary diversity of the family and moreover the possibility that paralogs mediating and controlling resistance to different ribosome-targeting antibiotics may have repeatedly arisen within the family independently from one another in the course of evolution^9,33,34,88^ (**Fig. 1a**).

Our structural analyses demonstrate that ABCF proteins function as master plumbers of the PTC and the adjacent entrance to the NPET in the 50S subunit of the ribosome (**Figs. 4, S9, & S15**). The ABCFs directly control the stereochemistry in the PTC via interaction of their PtIM, the conserved central domain that is the defining feature of the ABCF protein family (**Fig. 1b-d**), with the tRNA bound in the P site of the ribosome (**Figs. 2, 5, S7, & S12-13**). The 3’-hydroxyl of the ribose group in the 3’-terminal adenosine of this tRNA, which is the final nucleotide in the acceptor stem, forms the acyl linkage to the nascent polypeptide chain, while the 2’ hydroxyl of this ribose group directly participates in shuttling a proton to its 3’-hydroxyl during cleavage of that acyl linkage when the nascent chain is elongated. The 3’-terminal adenosine in the P-site tRNA thus directly forms part of the active site of the PTC along with adenine 2451, uracil 2584, and adenine 2602 from the 23S rRNA of the ribosome. The PtIM in different ABCF proteins either promotes adoption of the proper catalytic geometry by the acceptor stem of the P-site tRNA (*e.g.*, in EttA) or alternatively anything from modest (*e.g.*, in YbiT) to extreme (*e.g.*, in MsrE) disruptions of that geometry (**Figs. 2, 5, S7, S9, S12, S13, & S15**).

While these conformational changes are associated with 2°-25° rotations of the P-site tRNA around axes roughly extending from its acceptor stem to its anticodon loop (**Figs. 2, 5, S7, S12-S13**), all available ABCF-70S complexes show the ribosome in GS1 (Macrostate-I) with small negative rotations of the body of the 30S subunit compared to the 50S subunit and small positive rotations of the head of the 30S subunit compared to its body^77^ (**Figs. 2b, S7b, S10, & S13**). They furthermore all have the P-site tRNA in approximately its classical configuration maintaining canonical codon-anticodon contacts within the P site of the 30S subunit (**Fig. 6**). Notably, the stereochemistry at this site as well as the specific interaction geometry of the acceptor stem of the P-site tRNA with the PTC produced by each paralog (**Figs. 8, S1, S3, S9, & S15**) are maintained during the oscillations of the ABC domains of the EttA (**Fig. S1**) and YbiT within the E site of the ribosome in their hydrolytic conformations (**Movie S1**). These observations are consistent with maintenance of the local structural interactions at those two sites being a critical factor controlling ABCF protein evolution.

Given that the nascent polypeptide is covalently bonded to the 3’-terminal adenosine of the P-site tRNA, movement of the acceptor stem of the tRNA by the PtIM in ABCF proteins will directly pull on the nascent polypeptide chain, causing conformational changes to propagate through the PTC and into the NPET. Therefore, ABCF protein binding will generally modulate adventitious stereochemical interactions throughout this region comprising the catalytic core of the ribosome, including both problematic nascent chain conformations as well as direct blockage by antibiotics targeting the PTC/NPET. Even the ABCF proteins promoting proper catalytic geometry at the PTC will tend to have some activity of this kind because adventitious stereochemical interactions anywhere in the PTC/NPET are likely to disrupt the proper catalytic geometry and thus be reversed by structural interactions driving adoption of that geometry. The free energy of binding of the ABCF paralogs in their ATP-bound hydrolytic conformations will provide the thermodynamic driving force to disrupt adventitious stereochemical interactions in the PTC/NPET and pull the ribosome out of the conformational/kinetic trap caused by such interactions (center right in **Fig. 9**). For the ABCF proteins disrupting proper catalytic geometry at the PTC, release of the ABCF protein from the ribosome following ATP hydrolysis will give the P-site tRNA and covalently bonded nascent polypeptide chain an opportunity to return to proper catalytic geometry. The fundamental activity of ABCF proteins transiently modulating PTC/NPET stereochemistry upon ribosome binding will enable paralogs to influence numerous molecular processes influencing nascent chain elongation by the ribosome. This activity, combined with the opportunity for modulation of the rate of ATPase hydrolysis by ABCF proteins to tune nascent chain elongation kinetics (discussed below), will enable facile evolutionary optimization of activity to achieve specific biochemical goals via sequence variations in ABCF proteins.

Sequence variations in the PtIM are likely to play a key role in such evolutionary optimization of ABCF protein activity given that it is the only segment that directly contacts the acceptor stem of the P-site tRNA (**Figs. 3-4, S9, & S15**), and it sometimes also directly contacts nucleotides forming the PTC. However, these sequence variations are unlikely to have a simple relationship to activity because modulation of PTC/NPET stereochemistry occurs almost entirely indirectly through the movement of the acceptor stem of P-site tRNA by the PtIM (manuscript in preparation^83^). Analysis of the 13 different ABCF proteins for which 70S ribosome complex structures are available shows the length of the PtIM is strongly correlated with the displacement distance of the acceptor stem of P-site tRNA but not with the magnitude of conformational variation in the PTC/NPET **(Fig. 5b)**. Amino acids from the PtIM only penetrate into the PTC at all in two of these structures (just one residue in LsaA^20^ and three in MsrE^27^). Therefore, the vast majority of sequence variations in the PtIM do not directly influence PTC/NPET conformation but instead control the position of the terminal peptidylated adenosine in the P-site tRNA and thereby the direction and distance of displacement of the nascent polypeptide chain that is covalently bonded to that adenine. This effect will strongly modulate PTC/NPET conformation whenever a nascent polypeptide is bound to the P-site tRNA^33,34^, meaning in all physiological ribosome complexes except the 70S IC prior to the formation of the first peptide bond.

The disruption of nascent chain interactions with the PTC/NPET during ABCF-induced movement of the acceptor stem of the P-site tRNA is therefore likely to involve substantial conformational activation energy with the magnitude depending on the exact trajectory of that movement, which is controlled by the sequence of the PtIM and its structural interactions with that tRNA and the 50S subunit. We hypothesize that this activation energy effect (center right in **Fig. 9**) is a major determinant of ABCF protein specificity and therefore a key driver of evolutionary diversification in the sequence of the PtIM. To test this hypothesis, future research should prioritize kinetic and structural studies of ABCF protein interaction with ribosomal complexes with peptidylated P-site tRNAs, ideally including complexes with antibiotics or adventitious polypeptide sequences clogging the PTC/NPET.

The mechanochemical reaction cycle mediating ATP binding/hydrolysis by ABCF proteins ultimately controls the activity of their PtIM’s modulating PTC/NPET stereochemistry, meaning sequence variations controlling the kinetics of their mechanochemical reaction cycle are likely to represent another key contributor to the evolution of ABCF activity. The two available structures of nucleotide-free ABCF proteins^8,26^ both show a wide opening of the two ATP-binding sites located in the interface between the tandem ABC domains that flank the PtIM (**Fig. 1d**). This opening is incompatible with binding in the E site of the ribosome in the configuration observed in all of the available ABCF-70S ribosome complex structures, consistent with biochemical observations showing high-affinity ribosome binding is dependent on ATP binding to ABCF proteins^20–26^. Therefore, interfacial ATP binding closing the mechanochemical clamp formed by the ABC domains is effectively cooperatively coupled to the initial binding of an ABCF protein in the E site of the ribosome (center left in **Fig. 9**). Release of the ABCF protein from this site, which is driven by ATP hydrolysis (right in **Fig. 9**), is required for nascent polypeptide elongation to continue because tRNA-mRNA translocation requires the P-site tRNA to move into the space occupied by the ABCF protein. The rate of the initial interaction of the ATP-bound ABCF protein with the ribosome and its rate of ATP hydrolysis therefore provide two potential points for kinetic control of functional ABCF protein interaction with ribosomes dependent on sequence variations influencing ATP binding or hydrolysis.

Our structural analyses support the existence of a third potential point of kinetic control of ABCF protein function related to the mechanochemistry of ATP binding, specifically during the transition from a closed, but catalytically inactive, conformation of the ATPase active sites into the proper catalytically active conformation for ATP hydrolysis (center right in **Fig. 9**). A subset of 70S ribosome complex structures with both *Mtb*EttA^26^ (**Fig. S9**) and *E. coli* YbiT (**Figs. 8, S15, & S16**) demonstrate that, while ribosome binding requires ATP-dependent closure of the clamp formed by the ABC domains, it does not require them to adopt the proper conformation for ATP hydrolysis. These structures furthermore directly demonstrate that transition from such a closed but catalytically inactive conformation into the catalytically active conformation can be coupled to substantial changes in the binding geometry of the ABC domains and the PtIM (**Figs. 8, S9, & S16**), which means it could also potentially be coupled to the movement of the acceptor stem of the P-site tRNA and the covalently bound nascent polypeptide that is induced by PtIM binding (**Figs. 2, 5, S7, & S12-S13**).

The turnover rate for the complete mechanochemical reaction cycle will therefore depend on the initial binding rate, the rate of transition into the ATPase active conformation of the active sites, and also the rate of ATP hydrolysis once that conformation is achieved (**Fig. 9**). All of these rates can be influenced by sequence variations in the canonical ATP-binding motifs as well as other amino acids in the interface between the tandem ABC domains, which controls active-site formation. For example, the observation of a substantial population of non-catalytic EQ_2_-YbiT/IC complexes (**Figs. 8 & S16**) seems likely to be influenced by the sequence variations in the strongly non-canonical Signature Sequence in its ABCα1 subdomain. The rates of the sequential steps in the mechanochemical reaction cycle of ABCF proteins could also be influenced by allosteric effects modulating movement of the ABCα1 subdomain relative to the ATP-binding CORE subdomain based on the observation that these movements mediate formation of non-catalytic conformations of YbiT with closed ATP-binding sites (**Figs. 8 & S16**). Future studies should address how sequence variations in these protein segments and especially the non-canonical amino acids found in the ABC Signature Sequences in many ABCF proteins influence these rates and the thermodynamic efficiency of ABCF proteins.

Multi-stage kinetic control of the conformational/catalytic reaction cycle provides additional opportunities for evolutionary diversification of ABCF protein function and especially for the development of regulatory functions. The conformations of YbiT with closed but catalytically inactive ATP-binding sites, which are probably related to the sequence variations in its strongly degenerate Signature Sequence, seem likely to couple elongation to the presence/absence of the bL33 protein on the ribosome (**Figs. 8 & S16**). EttA has been demonstrated to either promote or inhibit peptide bond synthesis dependent on ATP/ADP ratio^8,67^, and the structural mechanism underlying this functional kinetic control of elongation involves subdomain movements similar to those in the conformations of YbiT with closed but catalytically inactive active sites (manuscript in preparation^65^). In bacteria, variations in the kinetics of ABCF protein interaction could potentially have either a positive or negative effect on transcription level by decreasing or increasing the elongation rate in “translational speedometers”^9,89–93^; these sequences contain short upstream open reading frames encoding “leader peptides” whose translation rates control transcriptional antitermination by modulating the kinetics of mRNA folding between that short ORF and downstream transcriptional pause sites in the same transcript^94–97^. Finally, extending the residence time of an ABCF protein on the ribosome with closed but catalytically inactive ATP-binding sites could potentially recruit recycling factors to ribosomes with intractable stereochemical problems in the PTC/NPET (center in **Fig. 9**), in which case the ABCF would serve a related signaling function when it is unable to move the acceptor stem of the P-site tRNA to resolve the stereochemical problem that led to its recruitment. The flexibility in biochemical function that is possible by combining multi-stage kinetic control of ATPase activity with the diverse stereochemical effects that the PtIM can exert on the PTC/NPET of the ribosome likely explains the great evolutionary diversity of the ABCF protein family.

## Abbreviations

ABCF: ATP-binding cassette family F
A site: ribosomal aminoacylated-tRNA binding site
E site: ribosomal tRNA exit site
FSC: fourier shell correlation
H-bond: hydrogen bond
mRNA: messenger RNA
*Mtb*: *mycobacterium tuberculosis*
NPET: nascent peptide exit tunnel
P site: ribosomal peptidyl-tRNA binding site
PTC: peptidyltransferase center
rRNA: ribosomal RNA
RMSD: root-mean-square variation

## Acknowledgements

This work was supported by grants from the US NIH-NIGMS to JFH (GM127883) and RLG (GM084288 and GM137608). GB was supported by funds from the CNRS (UMR8261), Paris Cité University, the LABEX program (DYNAMO ANR-11-LABX-0011), and the French ANR (EZOtrad/ANR-14-ACHN-0027 and ABC-F_AB/ANR-18-CE35-0010). Cryo-EM data were collected at the Columbia Cryo-EM Core Facility and the New York Structural Biology Center (NYSBC). The authors thank the NYSBC staff for assistance with cryo-EM data collection and Israel Fernandez for advice.

## Data availability

The complete cryo-EM datasets, final maps, and refined coordinate models for all structures reported in this paper have been deposited in the EMPIAR, EMDB, and PDB databases, respectively. Identification codes for all of these depositions are in Tables S1-S3.

## Author contributions

JFH, RLG, JF, GB, and SS conceived the project. The experiments were designed by SS, RCG, NAB, RLG, and JFH, and they were performed by SS, RCG, ZF, CW, CGA, NAB, KH-W, RAG, ZZ, TC, and MJN. The data were analyzed by SS, RCG, ZF, KKR, CGA, GB, and JFH. SS and JFH drafted the manuscript and finalized it with help from all authors.

## Conflict of interest

JFH and JF are members of the Scientific Advisory Boards of Nexomics Biosciences Inc. and Viron Inc., respectively. The other authors declare no competing financial interests.

*Note: The ***Supplementary Information*** for this manuscript contains 17 figures and 4 tables.*

## Methods

### Sequence alignment and phylogentic tree inference

Protein sequences for the four ABCF paralogs from *E. coli* and their orthologs were compared to evaluate the sequence similarities and variations within the different subdomains and key functional motifs. The ARE ABCFs VmlR and MsrE and ABCFs from eukaryotes like Arb1 in *S. cerevisiae* and ABCF1 from *H. sapiens* were included to illustrate the extent of sequence similarities/variations in ABCF across the different genera. Clustal Ω^98^ was used with the default BLOSUM62 similarity matrix for the multiple sequence alignment, which was formatted using ESPript (https://espript.ibcp.fr)^99^. The guide tree in **Fig. 1a** was produced by Clustal Ω^98^ using the average distance method.

### ABCF protein purification

Plasmids containing the EQ_2_ variants of the four *E. coli* ABCFs under a pBAD promoter were transformed in to BL21-LobSTR strain^100^. Single colony was picked and added to LB medium under Ampicillin selection (100µg/ml) and the cells were grown at 37°C overnight with 4% glucose. The overnight culture was diluted 1/100 in Terrific Broth/Amp and grown at 37°C until the OD_600_ reached 2. At this point the cells were induced with 0.1% L- Arabinose and grown for another 3 hours. The cells were harvested, washed with PBS and stored at −80°C until the next step. The cells were lysed in buffer A (20 mM Tris (pH 7.5 at 4 °C), 300mM NaCl, 10mM imidazole, 2mM 2-Mercaptoethanol (BME), 10% glycerol) and the proteins purified using Ni-NTA affinity chromatography in buffer C (20 mM Tris (pH 7.5 at 4 °C), 150mM NaCl, 500mM imidazole, 2mM 2-Mercaptoethanol (BME) and 10% glycerol). The elution fractions were pooled, concentrated and purified further on a HiLoad 16/60 Superdex 75 prep grade gel filtration column (GE Biosciences) in buffer containing 20 mM Tris (pH 7.5 at 4 °C), 150mM NaCl, 1 mM Tris (2-carboxyethyl) phosphine (TCEP) and 10% glycerol. Elution fraction corresponding to the EQ_2_-ABCFs were pooled, concentrated and stored at −80 in small aliquots.

### Preparation of ribosomal subunits and initiation factors

The 30S and 50S ribosomal subunits were purified from *E. coli* MRE600 strain, as described previously^64^. Tight-coupled 70S ribosomes were isolated using two rounds of ultracentrifugation on a 10-40% sucrose density gradient prepared in Ribosome Storage Buffer (10 mM Tris-acetate (pH 7.5 at 4 °C), 60 mM NH_4_Cl, 7.5 mM MgCl_2_, 0.5 mM EDTA, 6 mM BME). Tight-coupled 70S ribosomes from the second round of ultracentrifugation was solubilized in Ribosome Dissociation Buffer (10 mM Tris-acetate (pH 7.5 at 4 °C), 60 mM NH_4_Cl, 1 mM MgCl_2_, 0.5 mM EDTA, 6 mM BME) and 30S and 50S subunits were isolated using ultracentrifugation on a 20-40% sucrose density gradient prepared in Ribosome Dissociation Buffer. The 30S and 50S subunits were pooled separately, pelleted down, solubilized and passed through a second round of ultracentrifugation on a 20-40% sucrose gradient in Ribosome Dissociation Buffer. The 30S and 50S subunits were finally pooled individually, pelleted, solubilized in Ribosome Storage Buffer, snap frozen in liquid Nitrogen, and stored at − 80 °C in small aliquots. Initiation factors were purified as described previously^64^.

### Preparation of mRNAs and tRNAs

The mRNAs with a strong Shine-Dalgarno sequence (underlined) were used to control positioning of the codon in the P site (bolded and underlined), which was the methionine codon AUG for the 70S ICs and the valine codon GUU for the 70S ECs (PRE^-A^_Val_ complexes)^68^:

~~~
5’-GCAACCUAAAACUCACACAGGGCCC<u>UAAGGA</u>CAUAAAA**<u>AUG</u>**ACACCUAAUCCUCCUGCUGCACUCGCUGCACAAAUCGCUCAACGGCAAUUAAGGA-3’
5’-GCAACCUAAAACUCACACAGGGCCC<u>UAAGGA</u>CAUAAAA**<u>GUU</u>**ACACCUAAUCCUCCUGCUGCACUCGCUGCACAAAUCGCUCAACGGCAAUUAAGGA-3’
~~~

The mRNAs were *in vitro* transcribed using T7 RNA polymerase from a linearized plasmid DNA template using previously described protocol^101,102^. tRNA^fMet^ and tRNA^Val^ were obtained from MP Biomedicals, tRNA^fMet^ was aminoacylated and formylated and the yield of fMet-tRNA^fMet^, as assessed by hydrophobic interaction chromatography^64,101^ on a TSKgel Phenyl-5PW column (Tosoh Bioscience), was ∼95%.

### Assembly of 70S ICs and ECs (PRE^-A^_Val_ complexes)

All the ICs were assembled using the same preparation of ribosome subunits, while a different preparation of ribosome subunits was used to assemble the two PRE^-A^_Val_ complexes. The EQ_2_-EttA- and EQ_2_-YbiT-containing PRE^-A^_Val_ complexes were assembled using direct assembly with slight modifications to the previously optimized protocol^68^. 30 pmol mRNA, ∼20 pmol deacylated tRNA^Val^, and 15 pmol 30S subunits were added to a total reaction volume of 10 µl Ribosome Assembly Buffer (50 mM Tris hydrochloride, pH_25 °C_ = 7.5, 70 mM ammonium chloride, 30 mM potassium chloride, 6 mM 2- mercaptoethanol and 7 mM magnesium chloride) and incubated at 37 °C for 10 min. ∼11 pmol of 50S subunit was added to the reaction mixture and incubated for another 20 min at 37 °C. A 1:1 mixture of Mg-ATP and EQ_2_-ABCF (EttA/YbiT) was added to the reaction mixture to a final concentration of 3 µM and incubated at room temperature for 10 min and stored on ice. Finally, the reaction mixture was diluted in cold Tris-Polymix buffer (50 mM Tris acetate, pH_25 °C_ = 7.0, 100 mM potassium chloride, 5 mM ammonium acetate, 0.5 mM calcium acetate, 0.1 mM ethylenediamine tetraacetic acid, 10 mM 2-mercaptoethanol, 5 mM putrescine dihydrochloride and 1 mM spermidine, free base) to bring the final concentration of 50S subunit to ∼0.3 µM. The samples were then deposited on Quantifoil copper R1.2/1.3 300 mesh grids, coated with thin carbon layer. The EQ_2_-EttA and EQ_2_-YbiT 70S ICs were also prepared using direct assembly of the components where the individual concentrations of the components were increased such that the final concentration of the 50S subunit before being deposited onto UltraAu R1.2/1.3 300 mesh grids was ∼ 0.8 µM.

The EQ_2_-Uup- and EQ_2_-YheS-bound 70S ICs were assembled using enzymatic assembly. A 30S IC was assembled by mixing 1.2 µM each of IF1 and IF2 (with IF3 excluded to ensure formation of stable 70S ICs^64,103,104^), 1.2 µM of fMet-tRNA^fMet^, 1.6 µM of mRNA, 1.2 mM of GTP, and 0.8 µM of 30S subunit in Tris-Polymix buffer, followed by incubation at 37 °C for 10 min. Another round of incubation was carried out for 20 min at 37 °C after addition of 0.8 µM of 50S subunit to the 30S IC. A 1:1 mixture of Mg-ATP and EQ_2_-ABCF (Uup/YheS) was added to the reaction mixture to a final concentration of 7 µM and incubated at room temperature for 2 min and stored on ice before depositing on UltraAu R1.2/1.3 300 mesh grids.

### Cryo-EM sample preparation and data collection

The grids were glow discharged (in H_2_ and O_2_ for 25 s), using a Gatan Solarus 950 plasma cleaning system set to a power of 25W prior to sample vitrification. 3µL of the samples kept on ice were deposited on the grids, excess sample was wicked off and the grids plunge frozen using VitroBot Mark IV (Thermo Fisher). The grids with frozen vitrified samples were saved in liquid nitrogen until imaging. Grids were imaged on a 300kV Tecnai Polara F30 TEM (FEI) with a K3 detector or on a Titan KRIOS with K2 detector, refer to **Tables S1, S2** for details on magnification, pixel size and total dose used for imaging each sample.

### Image Processing, 3D reconstruction and analysis

All the datasets were processed in RELION and cryoSPARC^73,74^ (**S1-S6, Tables S1, S2**). Movies were motion corrected for beam-induced motion using MotionCor2^105^ implemented in RELION using a patch of 5X5. Contrast transfer function (CTF) estimation for the motion corrected images were determined using GCTF^106^. ∼1000 particles were manually picked to generate templates for particle picking. The final set of particles were autopicked using these templates, subjected to two rounds of reference free 2D classification; 2D class averages with 30S particles and junk were discarded at this stage. 3D reconstruction of the selected particles resulted in a consensus map. A mask corresponding to the 30S subunit was used for focused classification with signal subtraction to separate the 50S and 70S particles. Particles corresponding to 70S were refined further yielding a consensus map. A loose mask corresponding to the uL1-stalk, E-site bound factor and P-site tRNA was used for focused classification with signal subtraction to classify the 70S particles into classes with homogenous particles with empty 70S, 70S with tRNA at P and/or E-site, 70S with P-site tRNA and ABCF at E-site^61,62^. The particles subset for ABCF bound ICs or ECs was unbinned and refined to get a consensus map. Bayesian Polishing^107^ followed by another round of focused classification with or without signal subtraction was applied to classify the particles into homogenous subsets. The final subset of particles were refined in cryoSPARC using multiple rounds of non-uniform refinement^71^ and CTF refinement^108^ to derive the final maps for each of the EQ_2_-ABCF bound ICs or ECs and the ICs reported in this study. ResMap implemented in RELION was used to estimate the local resolution of the final maps^109^. FSC at 0.143 is used to report the global resolution of the final maps.

### Molecular model building, refinement, and validation

Model building was initiated by visually docking PDB ID 5u9f^110^ into the cryo-EM density map using USCF Chimera^111^. The fitted model was manually adjusted, built and real space refined in COOT^112^. The model corresponding to the ABCFs was manually built using the crystal structure of EttA (PDB ID 4fin)^8^. The final model of the complex was real space refined in PHENIX^113,114^. Cryo-EM validation tools in PHENIX were used for model validation^115,116^. The map and model statistics are reported in **Tables S1-S3**.

### Structure analysis methods

The inter subunit rotations of the different initiation and the elongator complexes reported in this paper and the reference ribosomes with or without factors at the E-site were calculated using RadTool^77^. Subdomain rotations of the ABCFs from *E. coli* and ARE ABCFs were calculated using the PyMol plugin draw_rotation_axis. Erodaxis^117^ was used for calculating tRNA rotations of the E-site factor bound ribosomes relative to the P-site tRNA in classical state from the reference *E. coli* ribosome (PDB ID 4v9d).

### Molecular graphics

Figures were generated using ChimeraX^118,119^ and PyMol (The PyMOL Molecular Graphics System, Version 2.0 Schrödinger, LLC). Graphpad PRISM 9 was used for generating graphs.

## Supplementary Information

### Supplementary Discussion

#### ATP binding and hydrolysis motifs in ABC domains

The strongly conserved sequence features within ABC domains (**Fig. 1d**) are all involved in the binding or hydrolysis of ATP. The CORE subdomain comprises an N-terminal antiparallel β-sheet unique to ABC domains together with a mixed parallel/antiparallel β-sheet with flanking α-helices that is shared with the ATP-binding domains in the F1 ATPase^120^ and other ATPases in the AAA+ protein superfamily. This structural module contains both the Walker A motif^120^ (GxxGxGKST), which forms an α-helical capping structure that ligates the α/β-phosphates of ADP/ATP, and the Walker B motif (φ_4_DE, with φ any hydrophobic amino acid), which forms a buried β-strand that terminates in an aspartate residue that H-bonds to both the conserved serine in the Walker A and a water molecule ligating the Mg+ cofactor of ADP/ATP. In ABC domains, the aspartate in the Walker B motif is followed immediately by an invariant glutamate residue that functions as the catalytic base that activates water for hydrolytic attack on ATP. The parallel β-strands flanking both sides of the Walker B β- strand also terminate in catalytically important residues that are invariant in ABC domains. One terminates in the glutamine at the N-terminus of the γP_i_S (described in the main text), which also ligates the Mg^++^ cofactor of ATP, while the other terminates in a histidine residue that directly ligates the γ-phosphate of ATP. The ABCα subdomain contains a bundle of three α-helices, one of which contains the ABC Signature Sequence (canonically LSGGQ but usually LSGGE in ABCF proteins), which is considered the hallmark of ABC Superfamily ATPase domains. This motif forms an α-helical capping structure that ligates the γ-phosphate of the ATP molecule bound to the Walker A/B motifs in the CORE subdomain in the second ABC domain to form one of the two catalytically active composite ATPase sites in the interface of the ATP-sandwich complex that is described in the main text. The second ATPase site is formed by the reciprocal interaction of the other CORE and ABCα subdomains in the two ABC domains forming the ATP-sandwich complex (**Fig. 1d**).

#### Contacts between each ABCF subdomain and components of the 70S ribosome

The ATP- binding core of ABC1 (CORE1) forms a composite ATP-binding site (“Site 1”) in combination with the ABCα subdomain of ABC2 (ABCα2) (**Fig. 1d**); these mutually interacting regions of the ABCF proteins both consistently interact with the 50S subunit, while ABCα2 also consistently makes extensive interactions with the P-site tRNA (**Figs. 2f & S7g**). CORE1 specifically interacts with 23S rRNA, protein bL33, and the uL1 stalk in the 50S subunit, while ABCα2 specifically interacts with protein uL5 in all paralogs and also protein bL33 in all except YheS. The ATP- binding core of ABC2 (CORE2) forms a composite ATP-binding site (“Site 2”) in combination with the ABCα subdomain of ABC1 (ABCα1) (**Figs. 1d**). CORE2 consistently interacts with components of the 30S subunit, specifically 16S rRNA and protein uS7 (**Figs. 2f & S7g**). CORE2 of Uup also makes limited contacts to protein uL5, while CORE2 of YheS also makes limited contacts to both the uL1 stalk and the P-site tRNA. ABCα1 consistently interacts with the uL1 stalk of the 50S subunit (**Figs. 2f & S7g**), as does the Arm motif that extends from the ABCα1 subdomain, although the Arm in Uup also makes limited contacts to 23S rRNA.

#### Detailed analyses of ribosome conformations in E. coli ABCF-bound 70S ribosome complexes

As indicated in the main text, the EQ_2_ variants of the *E. coli* ABCF proteins all stabilize the ribosome in Global State 1 (GS1), alternatively called Macrostate-I, in which the ribosomal subunits are in their “non-rotated” relative orientation and the P-site tRNA is in its “classical” configuration^75,76^ (**Figs. 2b, S7b, & S10**). However, the different complexes show small rotations in the orientations of the body and head domains of the 16S rRNA compared to standard reference ribosome in GS1. We analyzed these rotations using an angular convention recently developed by Whitford and colleagues^77^. We compared the orientations of the “body” region of the 16S rRNA relative to the 23S rRNA in our *E. coli* ABCF-bound 70S complexes and also the orientations of the “head” region of the 16S rRNA relative to its body (**Figs. 2b, S7b, & S10**). The reference ribosome employed in these calculations, which is in the classical non-rotated conformation, comes from a crystal structure (PDB ID 4v9d) containing two ribosomes in the asymmetric unit, one in the <u>c</u>lassical conformation harboring a P-site tRNA in the P/P configuration (4v9d_C_) and the other in the <u>h</u>ybrid conformation harboring a P-site tRNA in the P/E configuration (4v9d_H_)^78^. The IC complexes of the EQ_2_ variants of the *E. coli* ABCF proteins show small counterclockwise 16S rRNA body rotations ranging from −2.2° to −1.3° and small 16S rRNA head rotations relative to the body ranging from +1.1° to +4.2° (**Figs. 2b, S7b, S10 & Table S4a**). The largest observed rotation is in the 16S rRNA head of the EQ_2_-YbiT_H1/PtIM(a)_/IC structure, which produces the largest phosphate coordinate shifts compared to the 4v9d_C_ reference ribosome following alignment of the cores of the 23S rRNAs in the EQ_2_-ABCF-bound 70S IC complex structures (**Fig. 2e**). The 16S rRNA rotations observed in these complexes all reside close to the dominant cluster of anticorrelated body/head rotation values observed in a recent analysis of a large ensemble of *E. coli* ribosome structures^77^, although EQ_2_-EttA_H1_/IC and EQ_2_-YheS_H_/IC show ∼+1° higher head rotations than most ribosomes with equivalent body rotations, while EQ_2_-YbiT_H1/PtIM(a)_/IC shows an ∼2.5° higher head rotation (**Figs. 2b & S10**). The observed 16S rRNA body tilt angles in the *E. coli* ABCF-bound ribosomes (+1.1° to +1.7°) match expectations for their observed 16S rRNA body rotations based on the Whitford analysis^77^, and their 16S rRNA head tilt angles (+0.5° to +1.9°) also match expectations for their observed 16S rRNA head rotations except for EQ_2_- YbiT_H1/PtIM(a)_/IC, which shows a head tilt angle ∼1.5° lower than expected.

A set of representative 70S ribosome complexes with ARE ABCFs show similar non- rotated ribosome conformations to the *E. coli* ABCF paralogs, although with somewhat more positive body rotations ranging from −1.0° to +4.6° (**Figs. 2b, S7, & S10**) and somewhat more positive head rotations (∼+2°) than most ribosomes with equivalent body rotations^77^. The body tilt angles in the ARE structures are also generally ∼+2° greater than expectations for ribosomes with equivalent body rotations, while their head tilt angles are consistent with expectations for their observed head tilt rotations. The more positive body rotations and body tilt angles stabilized by the ARE factors are likely caused by the larger P-site tRNA displacements induced by these factors compared to the *E. coli* ABCF paralogs (**Figs. 5 & S10, S12-S13**), as explained in the next section. The slightly more positive head rotations in both the *E. coli* ABCF and ARE structures compared to ribosomes with equivalent body rotations are likely controlled by the binding geometry of their ABC2 domains (**Figs. 2b, 2f, 4, S7b, & S7f**).

The stem and the head of the uL1 stalk rRNA are rotated by +9° to +17° and by +16° to +26°, respectively, in the *E. coli* ABCF-bound 70S-IC complexes relative to the uL1 stalk configuration in the 4v9d_C_ reference ribosome, while the ARE ABCF complexes show a wider range of rRNA stem rotations from +12° to +42° but a similar range of rRNA head rotations from +18° to +27° relative to the same reference structure (**Fig. S10**). Because the uL1 stalk in the standard 4v9d_C_ reference structure^78^ is stabilized in a likely non-physiological state by inter- ribosome crystal-packing contacts, these rotation values must be interpreted relative those observed in other functional states of the ribosome calculated using the same 4v9d_C_ reference structure. (Similarly, the coordinate shifts in uL1 stalk of the ABCF-bound 70S IC complexes compared to 4v9d_C_ shown in **Fig. 2e** do not represent absolute shifts from a reference functional state of the ribosome, and only the differences in those shifts between different 70S complexes are meaningful.) **Fig. S10** shows that the uL1 rRNA stem and head rotations relative to the 4v9d_C_ reference ribosome range from +21° to + 26° and from +34° to + 35°, respectively, in *E. coli* ribosomes with different tRNA and elongation factor occupancies at several key stages of the physiological translation elongation cycle^85,121^. A cryo-EM structure of a 70S complex of Elongation Factor P (EF-P) (PDB ID 6enu)^58^, a significantly smaller translation factor that also binds in the E site of the ribosome (21 kDa vs. ∼60 kDa for the ABCFs), shows much larger 72° and 39° uL1 stem and head rotations, respectively, relative to the 4v9d_C_ reference structure. The stem and the head of the uL1 stalk rRNA in the ABCF complexes thus show relatively modest outward rotations away from the P-site tRNA and PTC compared to the physiological translation elongation complexes, by ∼4-17° and ∼8-19°, respectively. In contrast, the stem of the uL1 stalk rRNA in the EF-P complex shows a substantial inward rotation of ∼50°, which is needed to enable the uL1 stalk to contact this much smaller E-site factor at its binding site proximal to the P-site tRNA and PTC.

Plotting the rotation of the stem of the uL1 stalk rRNA against the 16S rRNA body/head rotations/tilts in all of these ribosome complexes shows no correlation^61,63,68,79,80^ (**Fig. S10b**). Therefore, as pointed out in the main text, the structural features of the ABCFs and EF-P independently control the uL1 stalk configuration and the rotational macrostate of 70S ribosome complexes harboring a P-site tRNA.

#### Detailed analyses of the ATP interactions in E. coli ABCF-bound 70S ribosome complexes

The Walker A and B motifs in the ATP-binding cores have a canonical sequence in both ABC domains in all four of the *E. coli* ABCF proteins except for a conservative serine-to-threonine substitution in the Walker A motif in ABC2 of Uup (GxxGxGKTT *vs.* the canonical GxxGxGKST sequence). The γ-phosphate switch (Q-loop) and the H-loop also have canonical sequences in all of the paralogs. All four of these sequence motifs interact with the phosphates in the bound ATP molecules in the standard pre-catalytic geometry for ABC Superfamily ATPases (**Fig. 7c**) in all of the EQ_2_-EttA, EQ_2_-Uup, and EQ_2_-YheS 70S complex structures and also in the EQ_2_-YbiT_H_ conformations, with the glutamate-to-glutamine (EQ) mutation in the catalytic base in the Walker B motif preventing ATP hydrolysis^35,56,122^.

In contrast, the Signature Sequences in the ABCα subdomains of the *E. coli* ABCF proteins all have more significant variations compared to the canonical LSGGQ sequence characteristic of ABC Superfamily ATPases (**Fig. 7c**), as alluded to above. Both ABC domains in YheS show a somewhat conservative phenylalanine substitution for the initial leucine in the Signature Sequence, which is mostly buried and does not contact ATP. ABC2 in all of the factors as well as ABC1 in EttA show a relatively conservative glutamate substitution for the final glutamine in the Signature Sequence, which H-bonds to a hydroxyl group of the ribose moiety in ATP in the standard catalytic geometry for ABC Superfamily ATPases^38,59^. This H-bond is maintained in the composite ATP- binding sites harboring the glutamate substitution at this site in the Signature Sequence (**Fig. 7c**) in the ABCF paralogs. Notably, ABC1 in Uup, YbiT, and YheS all have a strongly non- conservative substitution of tryptophan for glutamine at this site, and this tryptophan does not make either van der Waals contacts or an H-bond to the ribose of the ATP, leaving the hydroxyls of the ribose solvent-exposed in the composite ATP-binding sites harboring the tryptophan substitution at this site (**Fig. 7c**). While the influence of these amino acid substitutions in the Signature Sequence on the rate constants for ATP hydrolysis is unknown, variations in the atomic contacts to the ribose of ATP seem unlikely to abrogate ATPase activity as long as all of the canonical H-bonds observed in ABC Superfamily ATPases are maintained to the β/γ-phosphates of ATP, which is the case for both of the composite ATP-binding sites in EttA, Uup, and YheS and also for ATP-binding Site 1 in the YbiT_H_ conformations described below (**Fig. 7c & Table S4b**).

We selected two fiducial distances, designated “Separation” and “Slide” (**Fig. 7b & Table S4b**) for qualitative characterization of the conformational state and catalytic competence of the two interfacial composite ATP-binding sites in ABC Superfamily ATPases. We define Separation as the distance between the Cα atoms of the glycine immediately preceding the lysine in the Walker A (canonically GxxGxGKST) and the second glycine in the ABC Signature Sequence (canonically LSGGQ), while we define Shift as the distance between the Cα atoms of the lysine residue in the Walker A motif and the serine residue in the ABC Signature Sequence (**Fig. 7b**). The Separation and Slide distances are between 11.2-12.1 Å and 9.4-10.0 Å, respectively, in both of the composite ATP-binding sites in all of our structures of 70S ribosome complexes of the EQ_2_ mutants of EttA, Uup, and YheS (**Fig. 7c & Table S4b**). The consistency of the Separation and Slide distances in these structures combined with their conservation of all amino acids making H-bonds to the β/γ- phosphate groups of ATP support our inference above that both composite ATP-binding sites are catalytically competent in these ABCF paralogs. However, as pointed out in the main text, enzymological studies will be needed to verify this inference and to establish the influence of the non-canonical residues in their Signature Sequences on their catalytic rate constants and the details of the chemical and conformational dynamics accompanying ATP hydrolysis.

#### Detailed analysis of ATP-binding site configurations in the YbiT_H_ -bound 70S ribosome complexes

As indicated in the main text, our six reconstructions of EQ_2_-YbiT bound to 70S ICs or ECs (**Figs. 8, S2, S6, S13, & S15-S17** and **Table S2**) show three different configurations of its tandem ATP-binding sites, which we have designated YbiT_H_ (<u>H</u>ydrolytic), YbiT_I_ (<u>I</u>ntermediate), and YbiT<u>_N_</u> (Non-hydrolytic) based on structural analyses explained here. The YbiT_H_ conformations show essentially canonical contacts to β/γ-phosphates of the ATP molecule bound in ATP-binding Site 1 (*i.e.*, the composite site formed between the Walker A/B motifs in CORE1 in ABC1 and the Signature Sequence in the ABCα2 subdomain in ABC2), and these interactions are very similar to those at both ATP-binding sites in our reconstructions of 70S ribosome complexes of the EQ_2_ mutants of the other *E. coli* ABCF paralogs (**Fig. 7c** and **Table S4b**). As explained in detail below, the YbiT_H_ conformations show nearly but not entirely canonical contacts to β/γ-phosphates of the ATP molecule bound in ATP-binding Site 2 (*i.e.*, the composite site formed between the Walker A/B motifs in CORE2 in ABC2 and the Signature Sequence in the ABCα1 subdomain in ABC1). While analyses of the ATP interactions in our cryo-EM structures support the inference that Site 1 in YbiT is likely to be catalytically competent, there is greater uncertainty concerning the catalytic properties of Site 2.

The Signature Sequence in the ABCα1 subdomain in ABC1 in YbiT (*i.e.*, in ATP-binding Site 2) shows three non-conservative amino acid substitutions in addition to the tryptophan substitution in the terminal glutamine described in the previous section. These substitutions convert the canonical LSGGQ sequence to the strongly degenerate sequence VAPGW. In the YbiT_H_ conformations, the backbone nitrogen atom of the penultimate glycine residue, which is the only canonical amino acid retained in this highly degenerate sequence, makes an H-bond to the γ- phosphate of ATP in an equivalent geometry to that observed in the standard catalytic geometry observed in ATP-bound complexes of the EQ_2_ mutants of other ABC Superfamily ATPases with canonical Signature Sequences^38,46^ (**Fig. 7c**). However, the replacement of the canonical serine by alanine at the second position in the degenerate Signature Sequence in ABC1 in YbiT disrupts a conserved H-bond made by the sidechain of that serine to the γ-phosphate of ATP in the standard catalytic geometry. Except for this single H-bond, the other conserved H-bonds to the β/γ- phosphates of ATP observed in the EQ_2_ complexes of many other ABC Superfamily ATPase are all conserved in ATP-binding Site 2 in the YbiT_H_ conformations. The impact of the loss of this single conserved H-bond between a serine sidechain and the γ-phosphate of ATP on the catalytic competence of this composite ATP-binding site is unclear and requires experimental investigation.

#### Detailed analysis of ATP-binding site configurations in the YbiT_I_ -bound 70S ribosome complex

The conformation of YbiT in this complex shows an ∼4° rotation of the ABCα1 subdomain compared to the YbiT_H_ conformations. This rotation produces coupled shifts in the position of its degenerate Signature Sequence that forms part of ATP-binding Site 2 and simultaneously in the position the conserved catalytic residue Q71 in the γ-phosphate switch in ABC1 that forms part of ATP-binding Site 1. These shifts yield configurations of both ATP- binding sites intermediate between those observed in the YbiT_H_ and Ybit_N_ conformations (**Figs. 7c, 8a-b, d, & S16a-b**). Site 2 in this conformation shows a Slide distance of 11.4 Å, which is between the 10.1-10.4 Å and 14.8-15.1 Å distances observed in the YbiT_H_ and YbiT_N_ conformations, respectively (**Table S4b**), but the ∼1 Å increase in this fiducial distance corresponds to a 1.6 Å shift in the position of gly-159 (**Figs. S16a-b**), the only canonical residue remaining in the highly degenerate Signature Sequence in ABCα1. This movement breaks conserved the H-bond from the backbone nitrogen of gly-159 to the γ-phosphate of ATP in Site 2, which is likely to slow or stop ATP hydrolysis at Site 2 in the YbiT_I_ conformation. The rotation of the ABCα1 subdomain driving this reconfiguration in Site 2 also changes the conformation of γ- phosphate switch in ABC1, which produces a 4.3 Å shift in the position of Q71 that breaks its conserved H-bond to the to the γ-phosphate of ATP in Site 1 (**Figs. S16a-b**). Therefore, rotation of the ABCα1 subdomain in YbiT likely renders Site 1 catalytically inactive simultaneously with rendering Site 2 catalytically inactive.

---

**Table S1.**
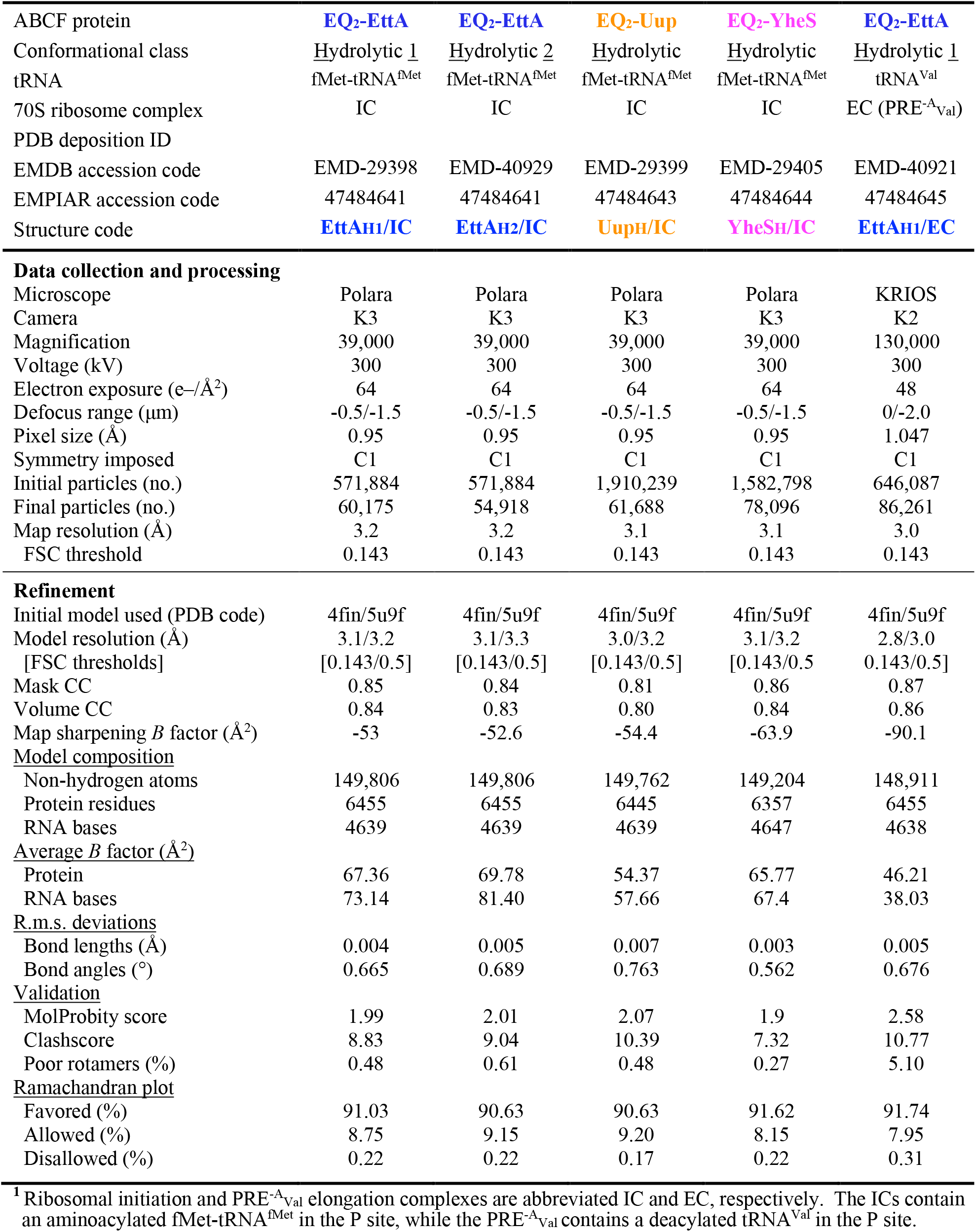
Cryo-EM data collection and refinement statistics for 70S complexes of E. coli EttA, Uup, and YheS. ^1^.

| ABCF protein | **EQ<sub>2</sub>-EttA** | **EQ<sub>2</sub>-EttA** | **EQ<sub>2</sub>-Uup** | **EQ<sub>2</sub>-YheS** | **EQ<sub>2</sub>-EttA** |
| --- | --- | --- | --- | --- | --- |
| Conformational class | <u>Hydrolytic 1</u> | <u>Hydrolytic 2</u> | <u>Hydrolytic</u> | <u>Hydrolytic</u> | <u>Hydrolytic 1</u> |
| tRNA | fMet-tRNA<sup>fMet</sup> | fMet-tRNA<sup>fMet</sup> | fMet-tRNA<sup>fMet</sup> | fMet-tRNA<sup>fMet</sup> | tRNA<sup>Val</sup> |
| 70S ribosome complex | IC | IC | IC | IC | EC (PRE<sup>-A<sub>Val</sub></sup>) |
| PDB deposition ID |  |  |  |  |  |
| EMDB accession code | EMD-29398 | EMD-40929 | EMD-29399 | EMD-29405 | EMD-40921 |
| EMPIAR accession code | 47484641 | 47484641 | 47484643 | 47484644 | 47484645 |
| Structure code | **EttAH1/IC** | **EttAH2/IC** | **UupH/IC** | **YheSH/IC** | **EttAH1/EC** |
| **Data collection and processing** |  |  |  |  |  |
| Microscope | Polara | Polara | Polara | Polara | KRIOS |
| Camera | K3 | K3 | K3 | K3 | K2 |
| Magnification | 39,000 | 39,000 | 39,000 | 39,000 | 130,000 |
| Voltage (kV) | 300 | 300 | 300 | 300 | 300 |
| Electron exposure (e<sup>-</sup>/Å<sup>2</sup>) | 64 | 64 | 64 | 64 | 48 |
| Defocus range (μm) | -0.5/-1.5 | -0.5/-1.5 | -0.5/-1.5 | -0.5/-1.5 | 0/-2.0 |
| Pixel size (Å) | 0.95 | 0.95 | 0.95 | 0.95 | 1.047 |
| Symmetry imposed | C1 | C1 | C1 | C1 | C1 |
| Initial particles (no.) | 571,884 | 571,884 | 1,910,239 | 1,582,798 | 646,087 |
| Final particles (no.) | 60,175 | 54,918 | 61,688 | 78,096 | 86,261 |
| Map resolution (Å) | 3.2 | 3.2 | 3.1 | 3.1 | 3.0 |
| FSC threshold | 0.143 | 0.143 | 0.143 | 0.143 | 0.143 |
| **Refinement** |  |  |  |  |  |
| Initial model used (PDB code) | 4fin/5u9f | 4fin/5u9f | 4fin/5u9f | 4fin/5u9f | 4fin/5u9f |
| Model resolution (Å) | 3.1/3.2 | 3.1/3.3 | 3.0/3.2 | 3.1/3.2 | 2.8/3.0 |
| [FSC thresholds] | [0.143/0.5] | [0.143/0.5] | [0.143/0.5] | [0.143/0.5] | [0.143/0.5] |
| Mask CC | 0.85 | 0.84 | 0.81 | 0.86 | 0.87 |
| Volume CC | 0.84 | 0.83 | 0.80 | 0.84 | 0.86 |
| Map sharpening *B* factor (Å<sup>2</sup>) | -53 | -52.6 | -54.4 | -63.9 | -90.1 |
| **Model composition** |  |  |  |  |  |
| Non-hydrogen atoms | 149,806 | 149,806 | 149,762 | 149,204 | 148,911 |
| Protein residues | 6455 | 6455 | 6445 | 6357 | 6455 |
| RNA bases | 4639 | 4639 | 4639 | 4647 | 4638 |
| **Average *B* factor (Å<sup>2</sup>)** |  |  |  |  |  |
| Protein | 67.36 | 69.78 | 54.37 | 65.77 | 46.21 |
| RNA bases | 73.14 | 81.40 | 57.66 | 67.4 | 38.03 |
| **R.m.s. deviations** |  |  |  |  |  |
| Bond lengths (Å) | 0.004 | 0.005 | 0.007 | 0.003 | 0.005 |
| Bond angles (°) | 0.665 | 0.689 | 0.763 | 0.562 | 0.676 |
| **Validation** |  |  |  |  |  |
| MolProbity score | 1.99 | 2.01 | 2.07 | 1.9 | 2.58 |
| Clashscore | 8.83 | 9.04 | 10.39 | 7.32 | 10.77 |
| Poor rotamers (%) | 0.48 | 0.61 | 0.48 | 0.27 | 5.10 |
| **Ramachandran plot** |  |  |  |  |  |
| Favored (%) | 91.03 | 90.63 | 90.63 | 91.62 | 91.74 |
| Allowed (%) | 8.75 | 9.15 | 9.20 | 8.15 | 7.95 |
| Disallowed (%) | 0.22 | 0.22 | 0.17 | 0.22 | 0.31 |

<sup>1</sup> Ribosomal initiation and PRE<sup>-A<sub>Val</sub></sup> elongation complexes are abbreviated IC and EC, respectively. The ICs contain an aminoacylated fMet-tRNA<sup>fMet</sup> in the P site, while the PRE<sup>-A<sub>Val</sub></sup> contains a deacylated tRNA<sup>Val</sup> in the P site.

**Table S2.** Cryo-EM data collection and refinement statistics for 70S ribosome complexes of E. coli YbiT. ^1^.

| ABCF protein | EQ2-YbiT | EQ2-YbiT | EQ2-YbiT | EQ2-YbiT | EQ2-YbiT | EQ2-YbiT |
| --- | --- | --- | --- | --- | --- | --- |
| ABC domain conformation | Hydrolytic 1 | Hydrolytic 2 | Intermediate | Non-hydrolytic 1 | Non-hydrolytic 2 | Non-hydrolytic 1 |
| PtIM configuration (a/b) | a | a | b | a | b | a |
| bL33 density | + | + | + | - | + | - |
| tRNA | fMet-tRNA<sup>fMet</sup> | fMet-tRNA<sup>fMet</sup> | fMet-tRNA<sup>fMet</sup> | fMet-tRNA<sup>fMet</sup> | fMet-tRNA<sup>fMet</sup> | tRNA<sup>Val</sup> |
| Ribosome complex | IC | IC | IC | IC | IC | EC (PRE<sup>-A</sup><sub>Val</sub>) |
| PDB deposition ID |  |  |  |  |  |  |
| EMDB accession code | EMD-29408 | EMD-40924 | EMD-40925 | EMD-40927 | EMD-40928 | EMD-40923 |
| EMPIAR accession code | 47484642 | 47484642 | 47484642 | 47484642 | 47484642 | 47484646 |
| Structure code | YbiT<sub>H1</sub>/PtIM(a)<br>/IC | YbiT<sub>H2</sub>/PtIM(a)<br>/IC | YbiT<sub>I</sub>/PtIM(b)<br>/IC | YbiT<sub>N1</sub>/PtIM(a)<br>/IC<sub>L33</sub>- | YbiT<sub>N2</sub>/PtIM(b)<br>/IC | YbiT<sub>N1</sub>/PtIM(a)<br>/EC<sub>L33</sub>- |
| **Data collection/processing** |  |  |  |  |  |  |
| Microscope | KRIOS | KRIOS | KRIOS | KRIOS | KRIOS | KRIOS |
| Camera | K2 | K2 | K2 | K2 | K2 | K2 |
| Magnification | 130,000 | 130,000 | 130,000 | 130,000 | 130,000 | 130,000 |
| Voltage (kV) | 300 | 300 | 300 | 300 | 300 | 300 |
| Electron exposure (e-/Å<sup>2</sup>) | 56.8 | 56.8 | 56.8 | 56.8 | 56.8 | 48 |
| Defocus range (μm) | -1.5/-2.5 | -1.5/-2.5 | -1.5/-2.5 | -1.5/-2.5 | -1.5/-2.5 | -0.5/-1.5 |
| Pixel size (Å) | 1.06 | 1.06 | 1.06 | 1.06 | 1.06 | 1.047 |
| Symmetry imposed | C1 | C1 | C1 | C1 | C1 | C1 |
| Initial particles (no.) | 577,295 | 577,295 | 577,295 | 577,295 | 577,295 | 1,005,322 |
| Final particles (no.) | 40,604 | 48,096 | 37,225 | 78,136 | 34,210 | 23,587 |
| Map resolution (Å) | 3.3 | 3.2 | 3.3 | 3.2 | 3.3 | 3.4 |
| FSC threshold | 0.143 | 0.143 | 0.143 | 0.143 | 0.143 | 0.143 |
| **Refinement** |  |  |  |  |  |  |
| Initial model used (PDB ID) | 4fin/5u9f | 4fin/5u9f | 4fin/5u9f | 4fin/5u9f | 4fin/5u9f | 4fin/5u9f |
| Model resolution (Å) | 3.2/3.3 | 3.2/3.3 | 3.2/3.4 | 3.1/3.3 | 3.2/3.4 | 3.1/3.5 |
| [FSC thresholds] | [0.143/0.5] | [0.143/0.5] | [0.143/0.5] | [0.143/0.5] | [0.143/0.5] | [0.143/0.5] |
| Mask CC | 0.85 | 0.84 | 0.83 | 0.87 | 0.85 | 0.83 |
| Volume CC | 0.83 | 0.82 | 0.81 | 0.84 | 0.84 | 0.82 |
| Map sharpening *B* (Å<sup>2</sup>) | -72.7 | -75.4 | -70.8 | -87.5 | -70.5 | -78.9 |
| **Model composition** |  |  |  |  |  |  |
| Non-hydrogen atoms | 149,741 | 149,741 | 149,864 | 149,408 | 149,825 | 151,790 |
| Protein residues | 6427 | 6427 | 6427 | 6383 | 6434 | 6698 |
| RNA bases | 4647 | 4647 | 4647 | 4647 | 4647 | 4642 |
| **Average *B* factor (Å<sup>2</sup>)** |  |  |  |  |  |  |
| Protein | 53.93 | 50.57 | 58.68 | 48.46 | 57.75 | 89.74 |
| RNA bases | 70.53 | 64.72 | 76.69 | 43.21 | 61.56 | 113.27 |
| **R.m.s. deviations** |  |  |  |  |  |  |
| Bond lengths (Å) | 0.007 | 0.006 | 0.008 | 0.005 | 0.007 | 0.009 |
| Bond angles (°) | 0.731 | 0.694 | 0.758 | 0.701 | 0.732 | 0.845 |
| **Validation** |  |  |  |  |  |  |
| MolProbity score | 1.99 | 1.98 | 2.02 | 1.9 | 1.98 | 2.32 |
| Clashscore | 7.79 | 8.17 | 8.71 | 7.25 | 9.05 | 18.68 |
| Poor rotamers (%) | 0.56 | 0.54 | 0.54 | 0.52 | 0.19 | 0.76 |
| **Ramachandran plot** |  |  |  |  |  |  |
| Favored (%) | 89.36 | 90.62 | 89.97 | 91.48 | 91.63 | 89.85 |
| Allowed (%) | 10.36 | 9.19 | 9.75 | 8.28 | 8.06 | 9.77 |
| Disallowed (%) | 0.28 | 0.19 | 0.28 | 0.24 | 0.32 | 0.38 |

<sup>1</sup> Ribosomal initiation and PRE<sup>-A</sup><sub>Val</sub> elongation complexes are abbreviated IC and EC, respectively. The ICs contain an aminoacylated fMet-tRNA<sup>fMet</sup> in the P site, while the PRE<sup>-A</sup><sub>Val</sub> contains a deacylated tRNA<sup>Val</sup> in the P site.

**Table S3.** Cryo-EM data collection and refinement statistics for 70S•PtRNA complexes. ^1^.

|  |  |  |
| --- | --- | --- |
| ABCF protein | - | - |
| bL33 density | + | - |
| tRNA | fMet-tRNA<sup>fMet</sup> | fMet-tRNA<sup>fMet</sup> |
| 70S ribosome complex | IC | IC |
| PDB deposition ID |  |  |
| EMDB accession code |  |  |
| EMPIAR accession code | 47484644 | 47484644 |
| Structure code | PtRNA/IC | PtRNA/IC<sub>L33-</sub> |
| **Data collection and processing** |  |  |
| Microscope | Polara | Polara |
| Camera | K3 | K3 |
| Magnification | 39,000 | 39,000 |
| Voltage (kV) | 300 | 300 |
| Electron exposure (e-/Å<sup>2</sup>) | 64 | 64 |
| Defocus range (μm) | -0.5/-1.5 | -0.5/-1.5 |
| Pixel size (Å) | 0.95 | 0.95 |
| Symmetry imposed | C1 | C1 |
| Initial particles (no.) | 1,582,798 | 1,582,798 |
| Final particles (no.) | 146,197 | 148,603 |
| Map resolution (Å) | 3.3 | 3.3 |
| FSC threshold | 0.143 | 0.143 |
| **Refinement** |  |  |
| Initial model used (PDB code) | EQ2-Uup<sub>H</sub>/IC | EQ2-Uup<sub>H</sub>/IC |
| Model resolution (Å) | 3.4/3.6 | 3.4/3.7 |
| [FSC thresholds] | [0.143/0.5] | [0.143/0.5] |
| Mask CC | 0.85 | 0.85 |
| Volume CC | 0.84 | 0.83 |
| Map sharpening *B* factor (Å<sup>2</sup>) | -124.7 | -125.4 |
| **Model composition** |  |  |
| Non-hydrogen atoms | 143,993 | 143,584 |
| Protein residues | 5,679 | 5,629 |
| RNA bases | 4,639 | 4,639 |
| **Average *B* factor (Å<sup>2</sup>)** |  |  |
| Protein | 76.95 | 75.33 |
| RNA bases | 88.88 | 92.64 |
| **R.m.s. deviations** |  |  |
| Bond lengths (Å) | 0.007 | 0.006 |
| Bond angles (°) | 0.775 | 0.744 |
| **Validation** |  |  |
| MolProbity score | 2.15 | 2.09 |
| Clashscore | 11.33 | 10.54 |
| Poor rotamers (%) | 1.08 | 0.80 |
| **Ramachandran plot** |  |  |
| Favored (%) | 89.77 | 90.13 |
| Allowed (%) | 10.07 | 9.69 |
| Disallowed (%) | 0.16 | 0.18 |

<sup>1</sup> Ribosomal initiation complexes containing an aminoacylated fMet-tRNA<sup>fMet</sup> in the P site are abbreviated IC.

**Table S4a.**
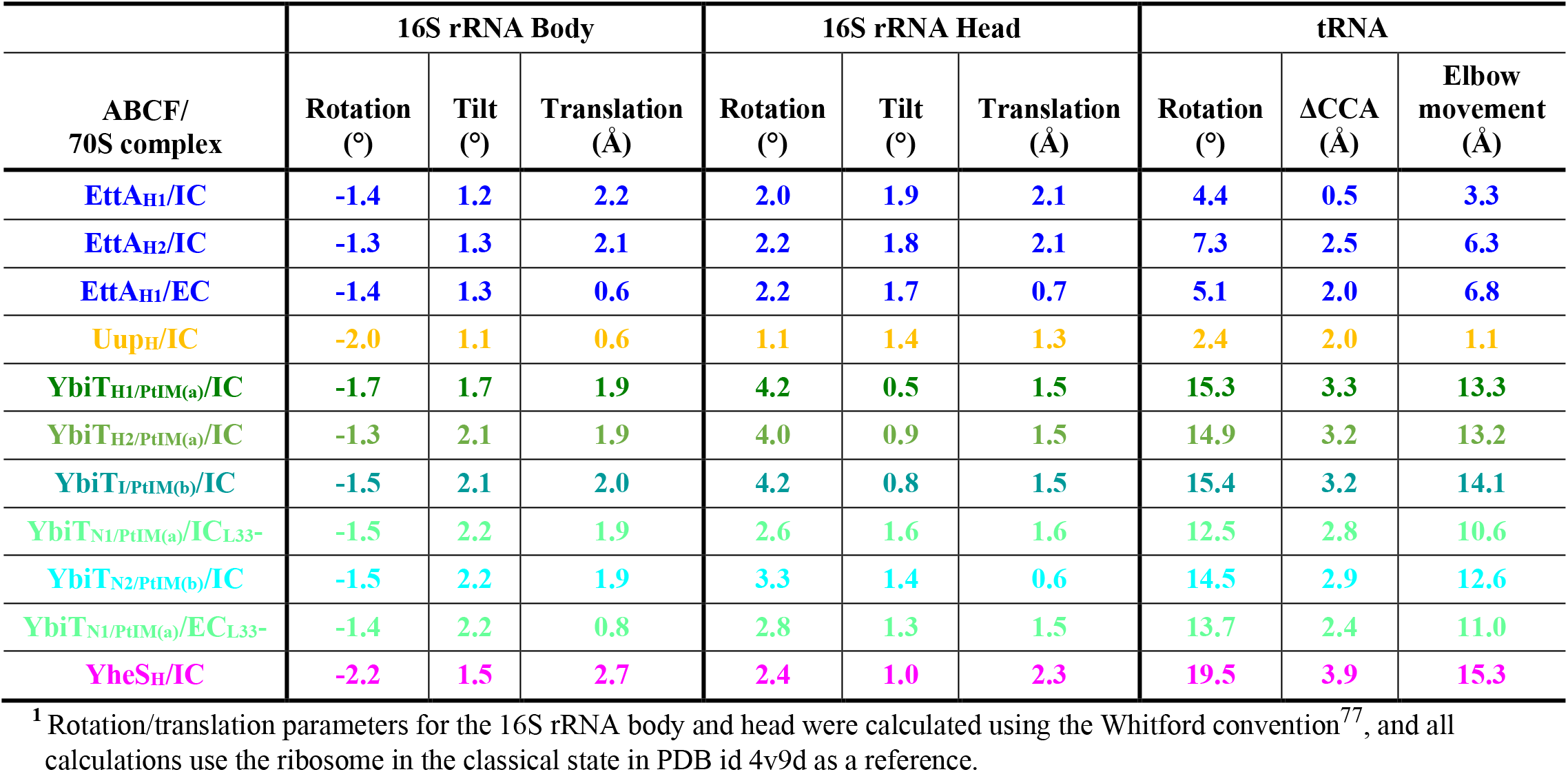
Fiducial distances at the ATPase active site in 70S-bound EQ_2_ variants of the E. coli ABCFs.

**Table S4b.**
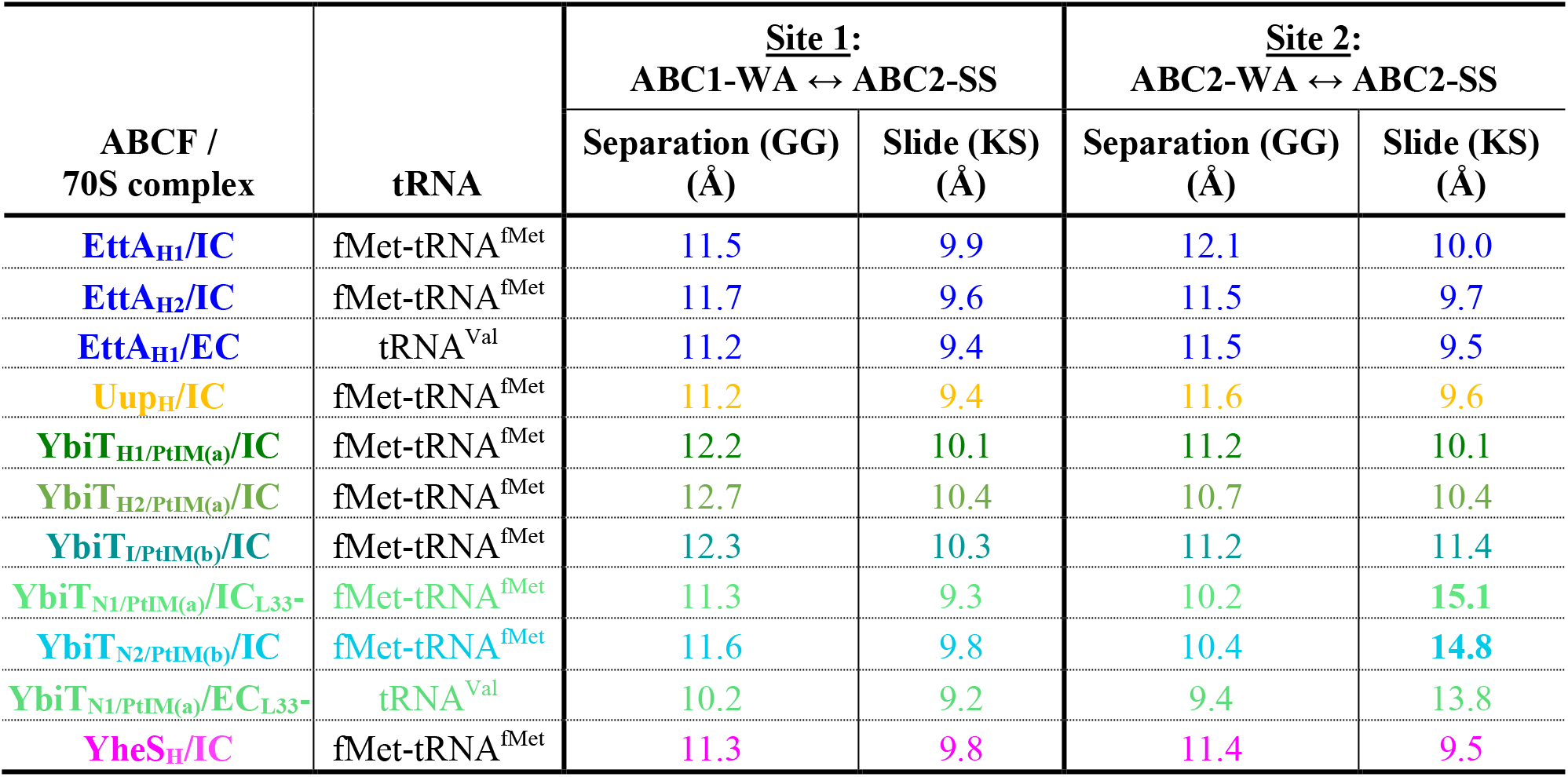
Fiducial distances at the ATPase active site in 70S-bound EQ_2_ variants of the E. coli ABCFs.

**Figure S1.**
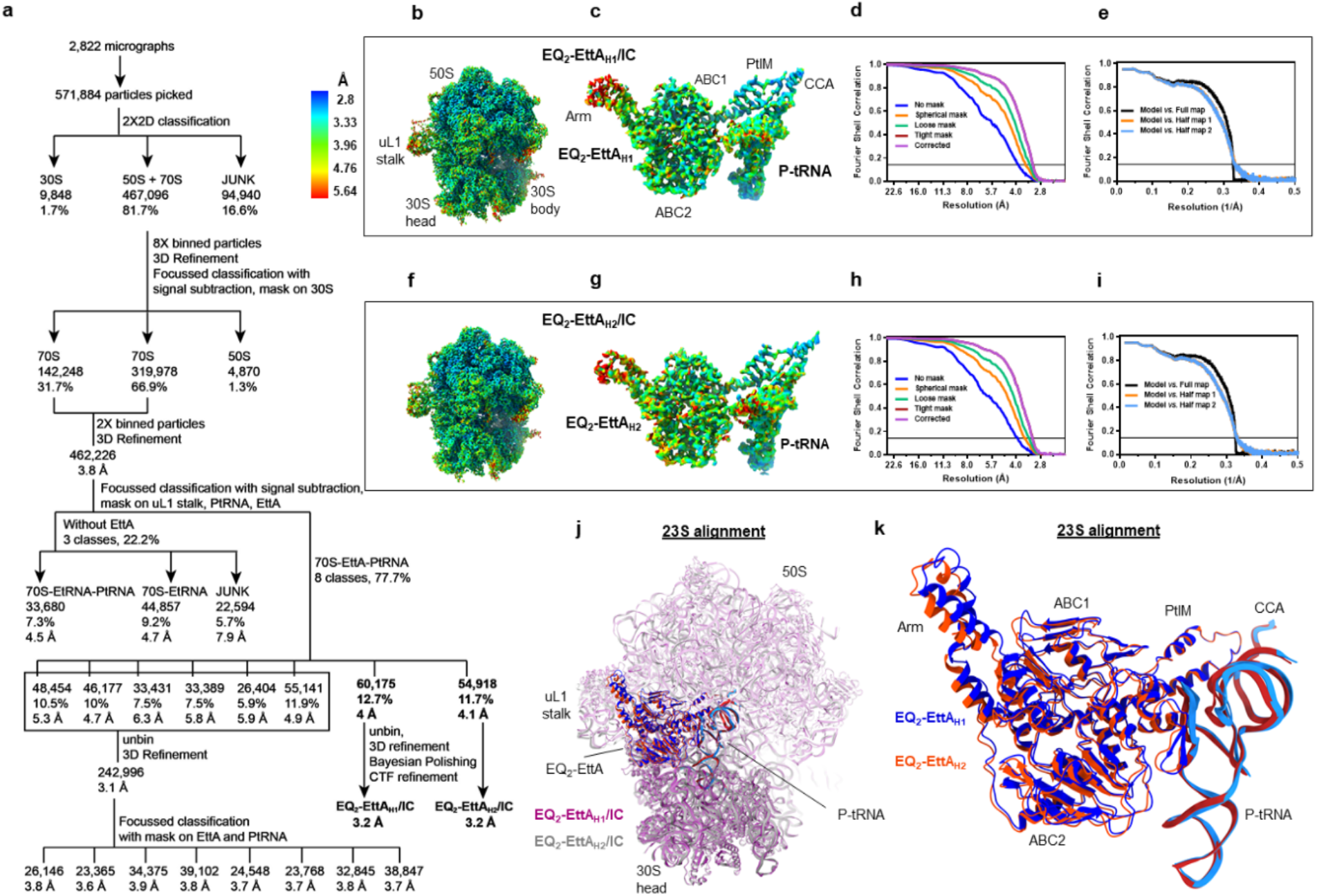
Cryo-EM imaging of EQ_2_-EttA added to a 70S IC. **(a)** Workflow of the cryo-EM data processing. The lower resolution classes in the branch at the bottom in panel **a** showed relatively poor density for EttA and the P-site tRNA. See the Methods for details. **(b, f)** Density maps for the two EQ_2_-EttA bound 70S IC complexes, colored according to the local resolution. **(c, g)** Isolated density maps for the EQ_2_-EttA and P-site tRNA in each complex, colored according to the local resolution. **(d, h)** FSC curves between the two half maps as a function of resolution, derived from cryoSPARC^74^ for the two 70S IC maps. Black line corresponds to an FSC of 0.143. **(e, i)** FSC curves between the refined coordinate model and the final sharpened map and the two half maps, derived from PHENIX^123^. **(j)** Superposition of the EQ_2_-EttA_H1_/IC and EQ_2_-EttA_H2_/IC complexes based on least-squares alignment in ChimeraX^111^ of the 23S rRNA in the 50S ribosomal subunit. **(k)** Zoomed-in view of the EQ_2_-EttA and the P-site tRNA in the superposition in panel **j**. The 70S IC classes containing EQ_2_-EttA that refined to high resolution (EQ_2_-EttA_H1_/IC and EQ_2_-EttA_H2_/IC) differ primarily in the orientation of the Arm motif and the adjoining helices in ABC1 subdomain of EttA and the uL1 stalk in 50S subunit (panels **j-k**).

**Figure S2.**
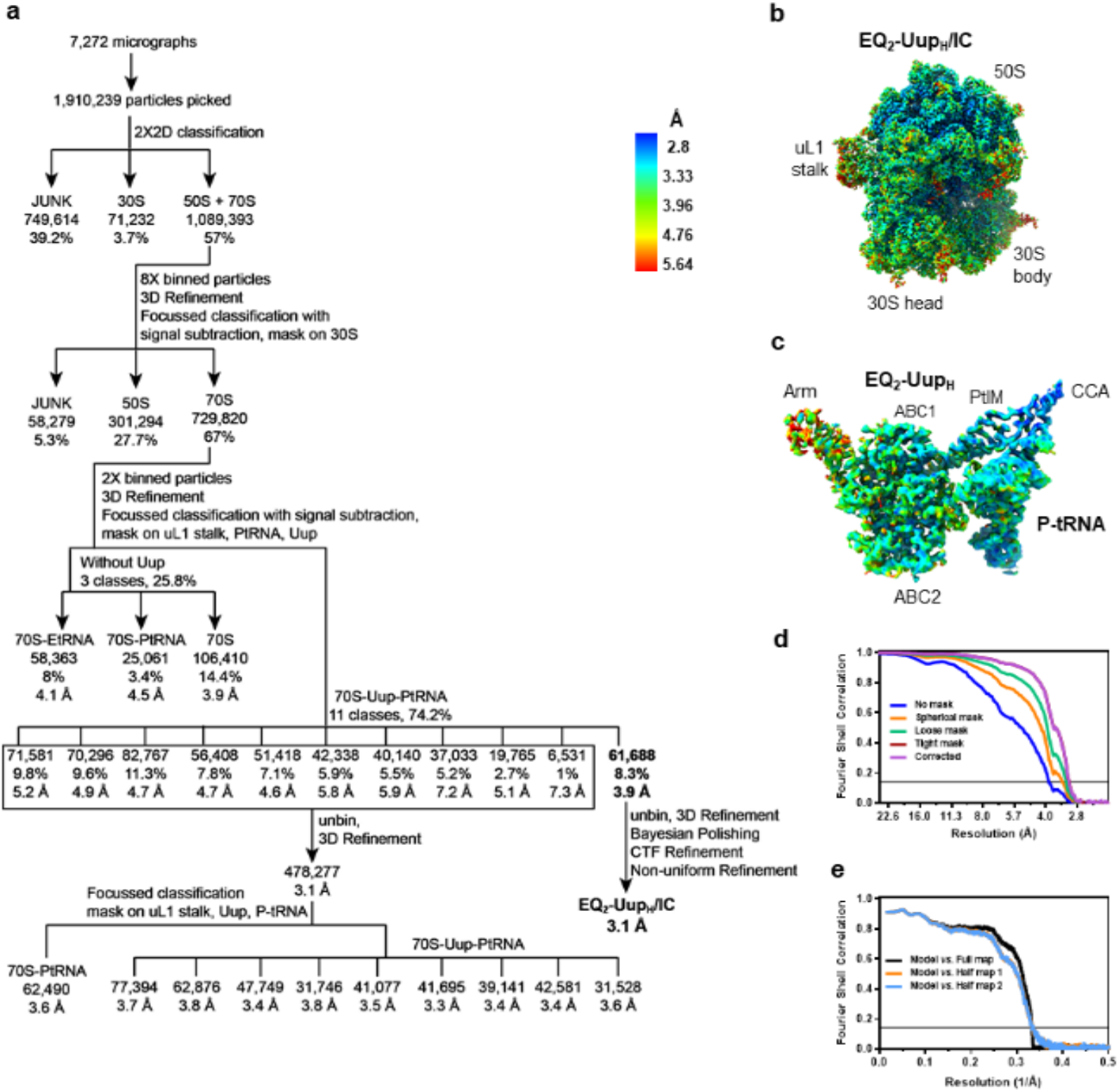
Cryo-EM imaging of EQ_2_-Uup added to 70S IC. **(a)** Workflow of the cryo-EM data processing for 70S-IC/ EQ_2_-Uup sample. The lower resolution classes in the branch at the bottom right showed relatively poor density for Uup and the P-site tRNA. **(b)** Density map for the EQ_2_- Uup bound 70S IC complex, colored based on local resolution. **(c)** Isolated density map for the EQ_2_-Uup and P-site tRNA from the complex, colored based on local resolution. **(d)** FSC curves between the two half maps as a function of resolution, derived from cryoSPARC^74^ for the 70S IC map. Black line corresponds to an FSC of 0.143. **(e)** FSC curves between the refined coordinate model and the final sharpened map and the two half maps, derived from PHENIX^123^.

**Figure S3.**
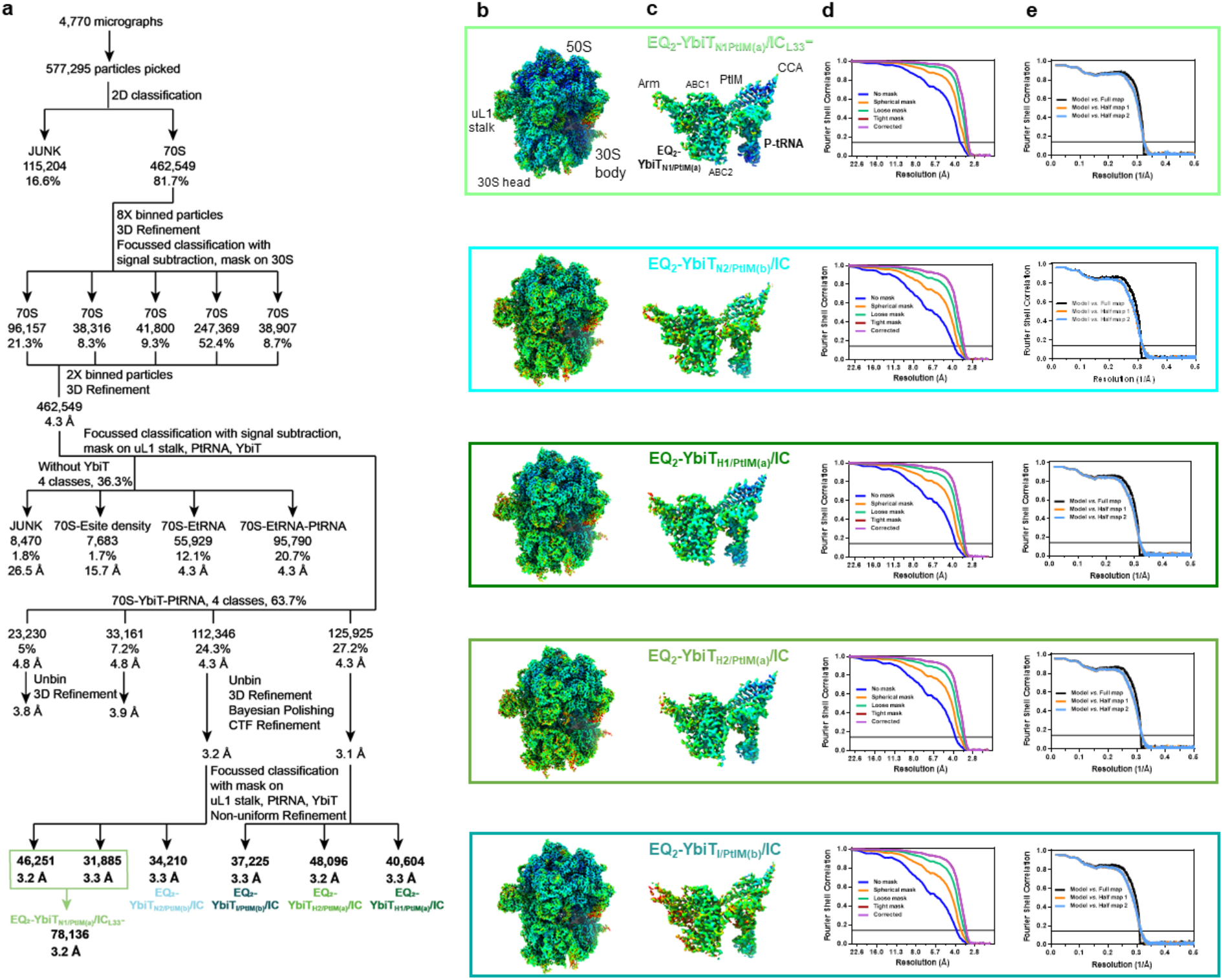
Cryo-EM imaging of EQ_2_-YbiT added to 70S IC. **(a)** Workflow of the cryo-EM data processing for 70S-IC/EQ_2_-YbiT sample. The lower resolution classes in the branch at the bottom right showed relatively poor density for YbiT and the P-site tRNA. The subscript defines the conformation at composite ATP binding site (H:hydrolytic, I: intermediate and N: non-hydrolytic); the different complexes displayed PtIMs in two conformations ((a) and (b)) on a 23S rRNA- alignment of the complexes. One of the EQ_2_-YbiT-bound 70S IC had the density for ribosomal protein bL33 missing (designated as L33-) **(b)** Density maps for the EQ_2_-YbiT bound 70S IC complexes for the five maps resolved to high resolution, colored based on local resolution. **(c)** Isolated density maps for the EQ_2_-YbiT and P-site tRNA from each of the five complexes, colored based on local resolution. **(d)** FSC curves between the two half maps as a function of resolution, derived from cryoSPARC^74^. Black line corresponds to an FSC of 0.143. **(e)** FSC curves between the refined coordinate model and the final sharpened map and the two half maps for each of the five maps, derived from PHENIX^123^.

**Figure S4.**
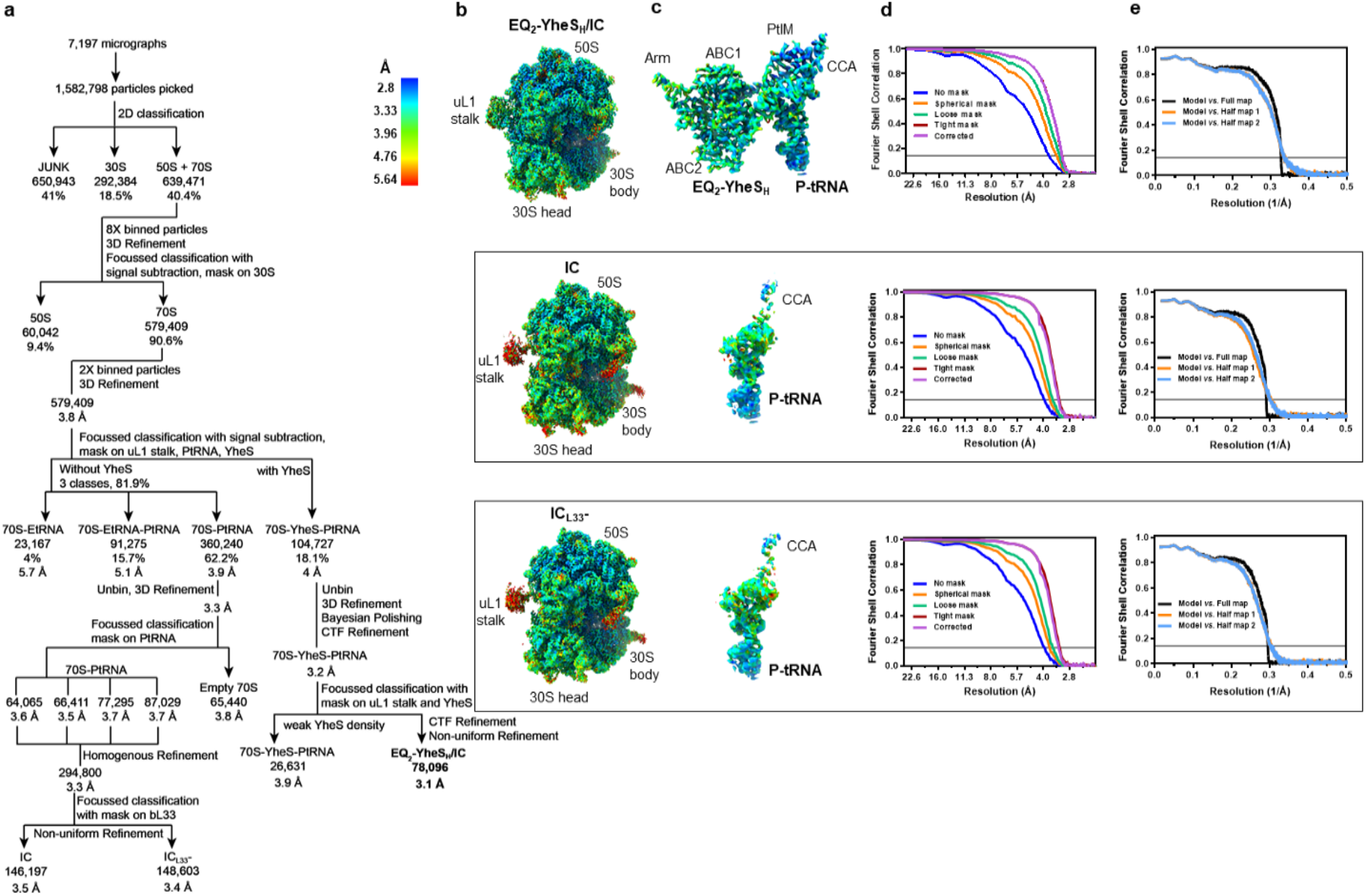
Cryo-EM imaging of EQ_2_-YheS added to 70S IC. **(a)** Workflow of the cryo-EM data processing for 70S-IC/EQ_2_-YheS sample. The lower resolution class in the branch at the bottom right showed relatively poor density for YheS. The 70S ICs without YheS were further classified into two groups IC and IC_L33_- (with no density for ribosomal protein bL33). **(b)** Density map for the EQ_2_-YheS-bound 70S IC and the 70S IC complexes, colored according to local resolution. **(c)** Isolated density map for the EQ_2_-YheS and P-tRNA and the P-site tRNA alone from the complex, colored according to local resolution. **(d)** FSC curves between the two half maps as a function of resolution, derived from cryoSPARC^74^. Black line corresponds to an FSC of 0.143. **(e)** FSC curves between the refined coordinate model and the final sharpened map and the two half maps, derived from PHENIX^123^.

**Figure S5.**
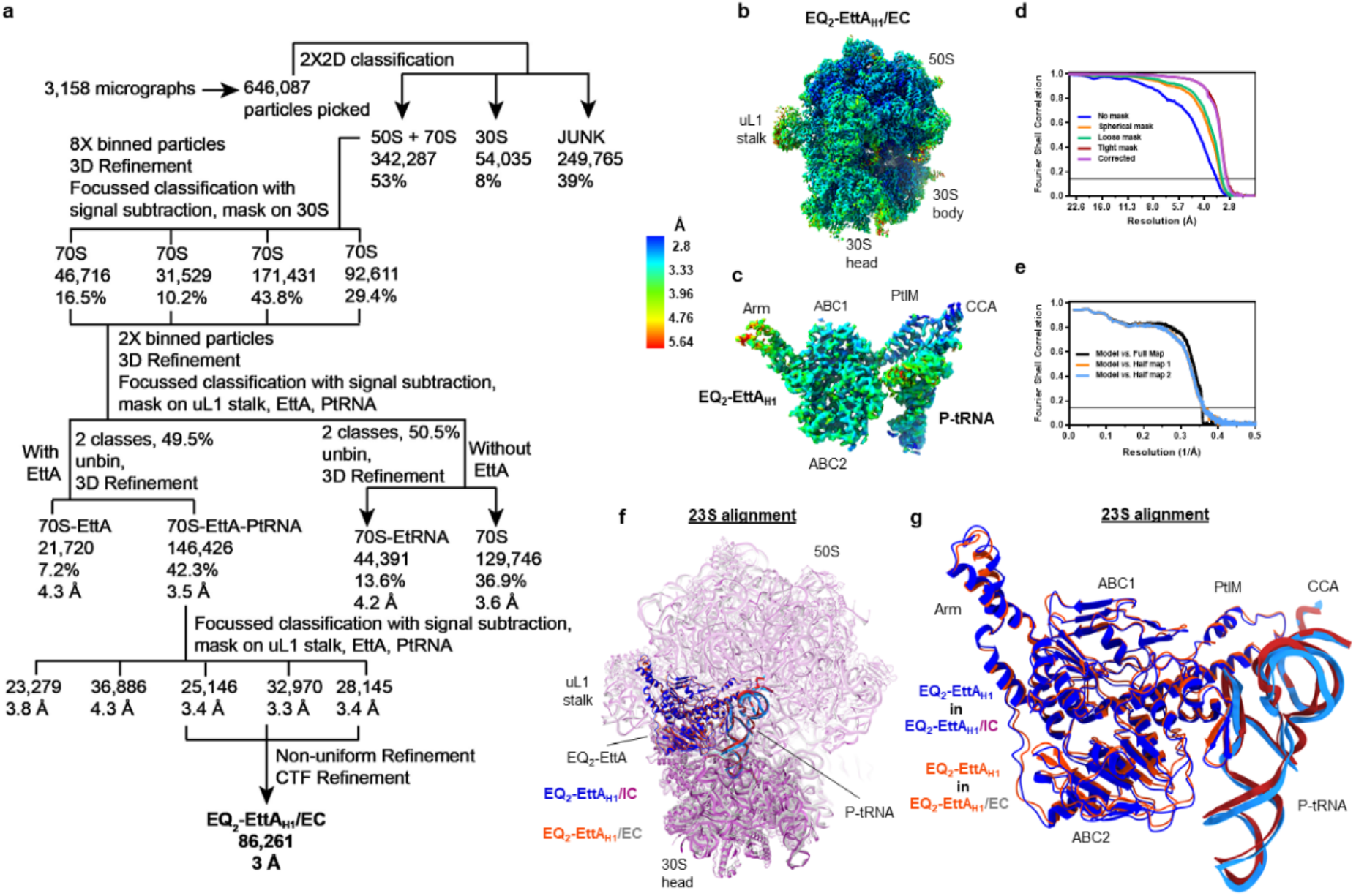
Cryo-EM imaging of EQ_2_-EttA added to 70S EC (PRE^-A^_Val_ complexes). **(a)** Workflow of the cryo-EM data processing. The lower resolution classes in the branch at the bottom left in panel **a** showed relatively poor density for EttA and the P-site tRNA. **(b)** Density map for the EQ_2_-EttA bound 70S EC, colored according to local resolution. **(c)** Isolated density map for the EQ_2_-EttA and P-site tRNA from the complex, colored according to local resolution. **(d)** FSC curves between the two half maps as a function of resolution, derived from cryoSPARC^74^ for the 70S EC map. Black line corresponds to an FSC of 0.143. **(e)** FSC curves between the refined coordinate model and the final sharpened map and the two half maps, derived from PHENIX^123^. **(f)** Superposition of EQ_2_-EttA_H1_/IC (ribosome in shades of purple and EttA/tRNA in shades of blue) and EQ_2_-EttA_H1_/EC (ribosome in shades of gray and EttA/tRNA in red) based on least- squares alignment in ChimeraX ^27,28^ of the 23S rRNA in the 50S subunit. **(g)** Zoomed-in view of the EQ_2_-EttA and the P-site tRNA in the superposition in panel **f**. The 70S IC and EC maps differ slightly in the positioning of the EttA and the tRNA in the two complexes (panels **f-g**)

**Figure S6.**
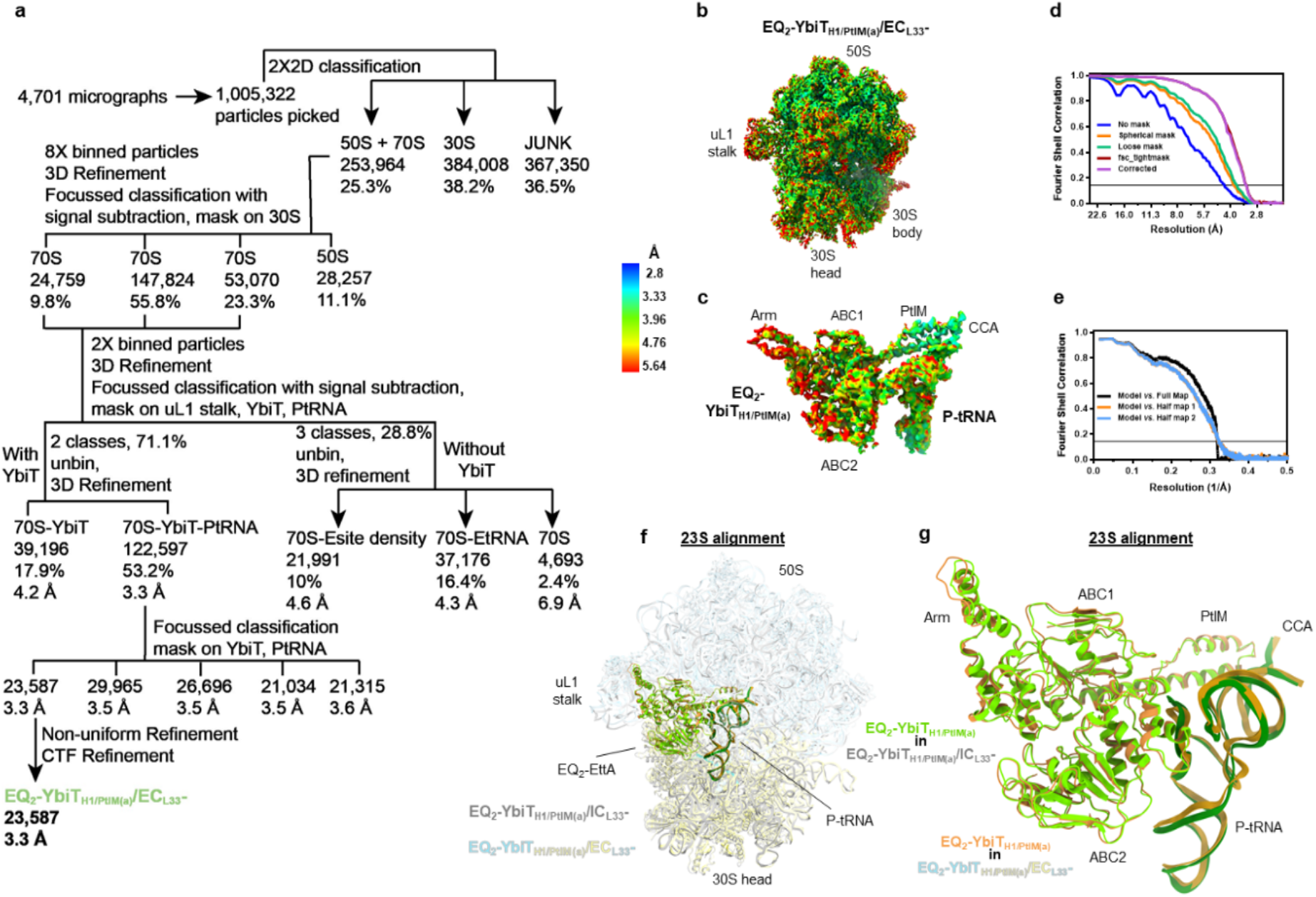
Cryo-EM imaging of EQ_2_-YbiT added to 70S EC (PRE^-A^_Val_ complexes). **(a)** Workflow of the cryo-EM data processing. The lower resolution classes in the branch at the bottom left in panel **a** showed very poor density for YbiT and the P-site tRNA. The subscript defines the conformation at composite ATP binding site (N: non-hydrolytic); with its PtIM in conformation “(a)” and the density for bL33 missing (designated as L33-). **(b)** Density map for the EQ_2_-YbiT- bound 70S EC complex, colored according to local resolution. **(c)** Isolated density map for the EQ_2_-YbiT and P-tRNA from the complex, colored according to local resolution. **(d)** FSC curves between the two half maps as a function of resolution, derived from cryoSPARC^74^ for the EQ_2_- YbiT-bound 70S EC map. Black line corresponds to an FSC of 0.143. **(e)** FSC curves between the refined coordinate model and the final sharpened map and the two half maps, derived from PHENIX^123^. **(f)** Superposition of the EQ_2_-YbiT_N1/PtIM(a)_/IC_L33_- and EQ_2_-YbiT_N1/PtIM(a)_/EC_L33_- complexes based on least-squares alignment in ChimeraX ^27,28^ of the 23S rRNA in the 50S subunit. **(g)** Zoomed-in view of the EQ_2_-YbiT and the P-site tRNA in the superposition in panel **f**.

**Figure S7.**
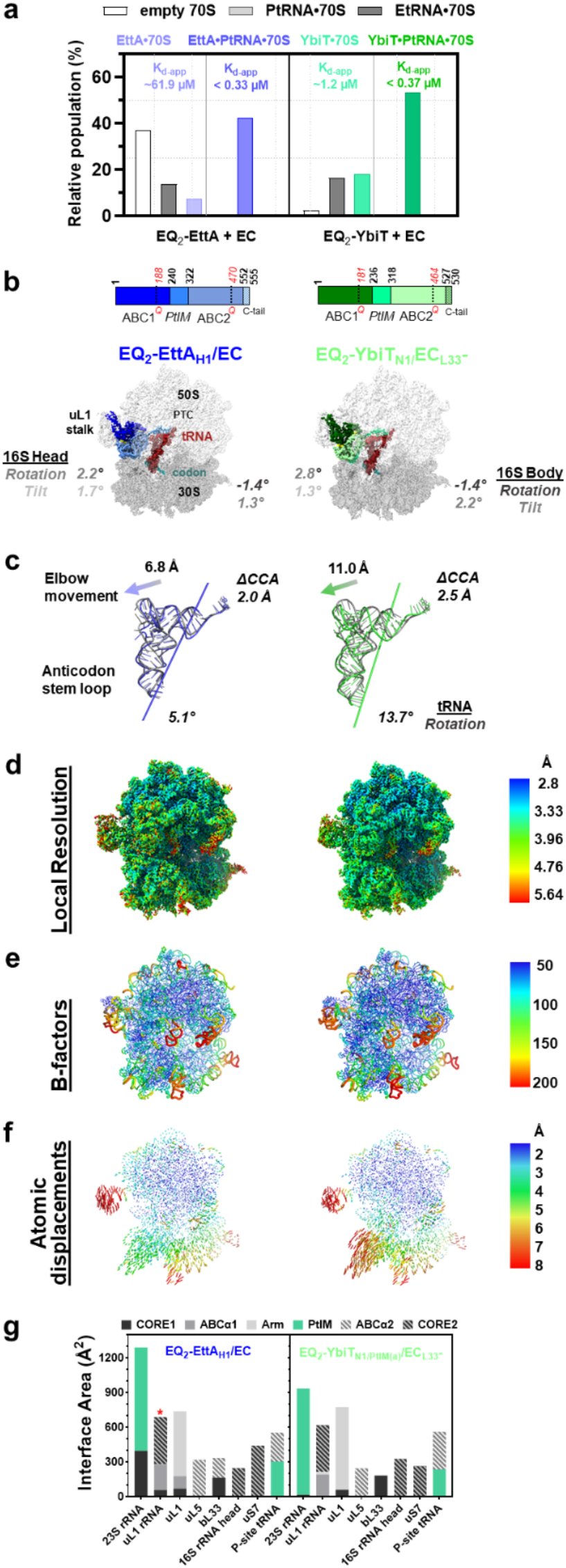
Cryo-EM structures of ATP-bound hydrolysis-deficient EQ_2_ mutants of E. coli EttA and YbiT bound to homologous 70S ECs (PRE^-A^_Val_) ribosome complexes. **(a)** Particle distribution in each dataset with *E. coli* ABCF paralogs EQ_2_-EttA and EQ_2_-YbiT added to EC, particles in each class presented here as bar graphs is a percentage of the total number of 70S ribosome-containing particles. **(b)** Color bars showing the general sub-domain organization of the ABCFs with the residue numbers demarcating each subdomain and the position harboring the E to Q mutation in each subdomain shown in red. Density maps of the *E. coli* EQ_2_-ABCF bound 70S ECs showing the 30S subunit in dark gray, 50S subunit in light gray, P-site tRNA in brick red, mRNA in teal, and two *E. coli* ABCFs color coded as in the color bar above. Values indicating intersubunit orientations of these complexes compared to the reference ribosome (4v9d_C_ PDB ID 4v9d), are indicated adjacent to each complex. **(c)** Rotation and shift of the terminal CCA of the acceptor stem and the elbow of the P-site tRNA relative to the positions that are observed in the classical configuration of the tRNA in the reference ribosome (4v9d_C_). **(d)** Density maps of the ABCF-bound ECs colored according to local resolution. (**e**) Cartoon putty representation of the ABCF-bound 70 ECs, thicker regions of the tube represent regions with high B-factor, the color coding varies blue to red for lower to higher B-factor values, as shown in the bar on the right. **(f)** Displacements (Å) of the phosphates in the 16S and 23S rRNAs from the ABCF-bound 70S ECs relative to those in the 4v9d_C_ complex (aligned on 23S rRNA), color-coded blue (smallest) to red (largest). **(g)** Bar graphs showing buried surface area between the subdomains in *E. coli* EQ_2_- ABCF paralogs and the ribosomal elements and the P-site tRNA, in the ABCF-bound 70S ECs. The asterisk (*) in the EQ_2_-EttA panel indicates that its CORE2 contact to the uL1 stalk involves residues in its C-terminal tail that extends from the CORE2 subdomain and is not conserved in the other paralogs.

**Figure S8.**
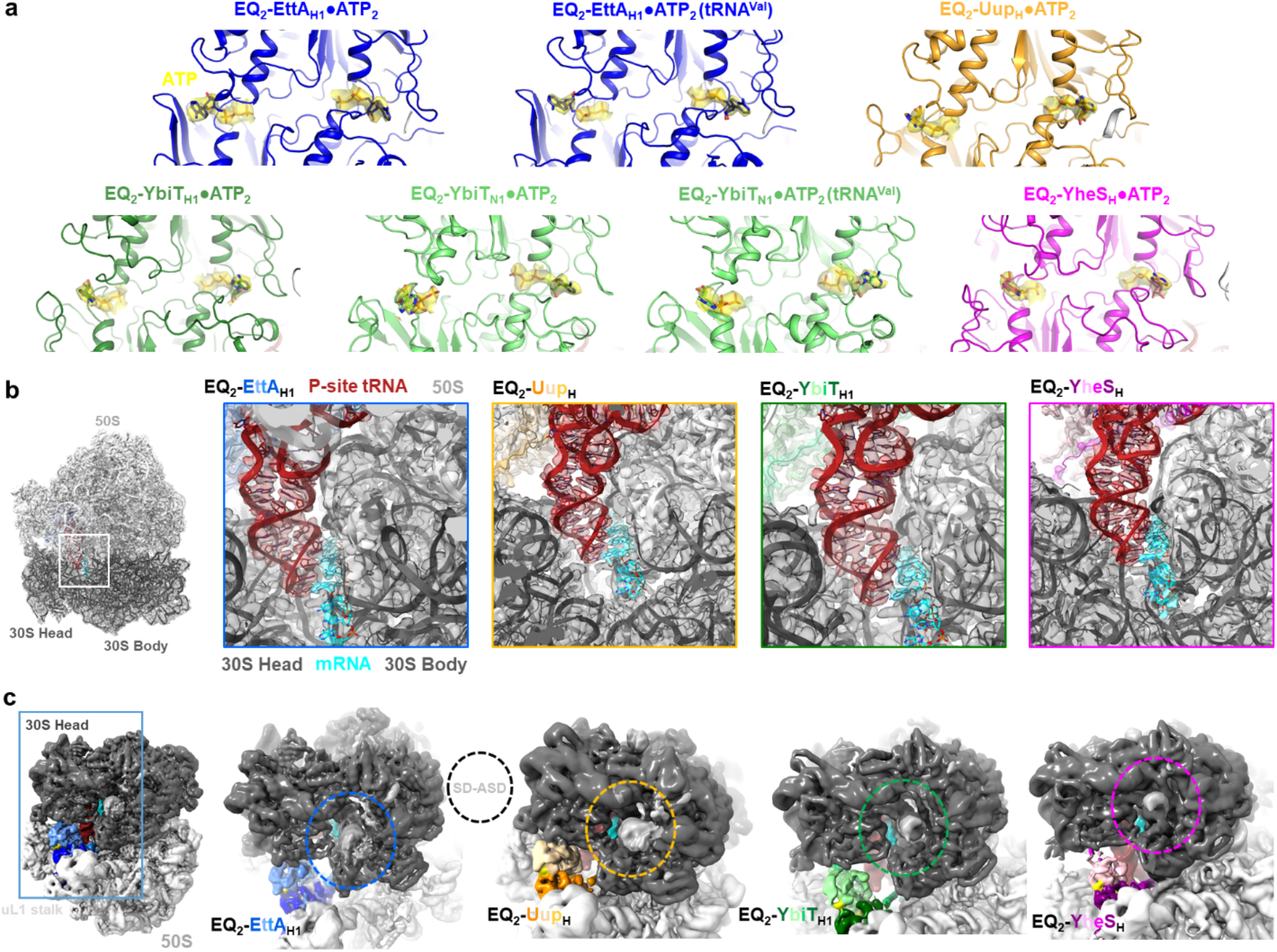
Cryo-EM maps show clear density for the two ATP molecules and minimal density in the mRNA exit channel in the E. coli EQ_2_ -ABCF bound 70S ribosome complexes. **(a)** *E. coli* EQ_2_-ABCFs showing density for the ATP (yellow) **(b)** *E. coli* EQ_2_-ABCF bound 70S ICs showing the density for the 30S subunit in dark gray, 50S subunit in light gray, and mRNA in cyan which is resolved clearly at the A and P sites; P-site tRNA is shown in scarlet red and the different EQ_2_-ABCFs in the 70S ICs are shown in three different shades (ABC2 – in dark, PtIM - medium and ABC1 - light) blue, orange, green, and magenta for EttA, Uup, Ybit and YheS, respectively. **(c)** Rotated view of the mRNA exit channel with the volume for Shine-Dalgarno anti-Shine- Dalgarno (SD-ASD, dashed circle) from 16S rRNA shown in lighter shade of gray relative to the rest of the 30S subunit.

**Figure S9.**
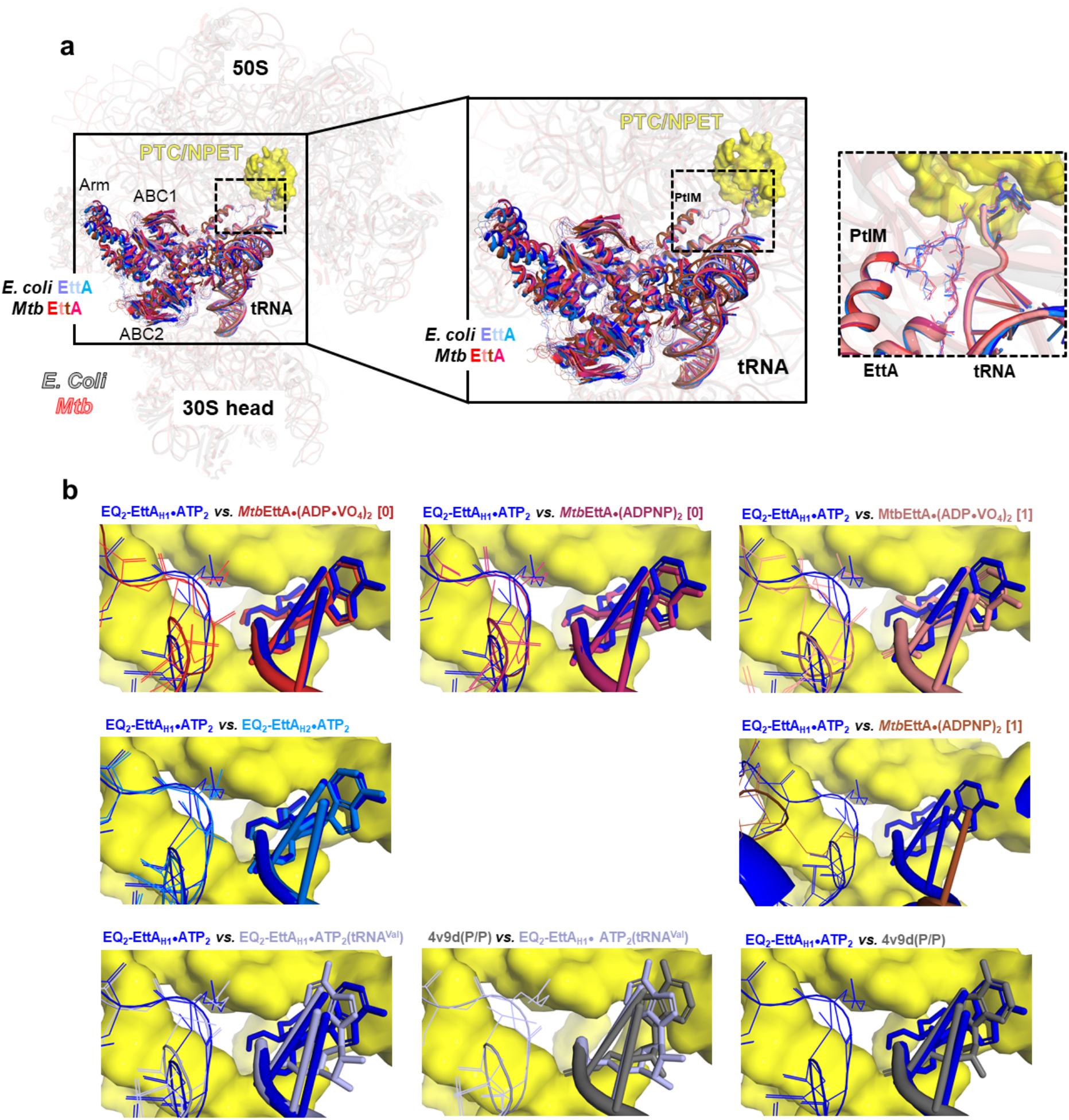
Comparison of PTC stereochemistry observed in EttA cryo-EM structures. **(a)** *E. coli* EQ_2_**-**EttA (shown in shades of blue)-bound 70S ribosome complexes (gray) compared to *Mtb*EttA with ADPNP (PDB IDs: 7msc (hot pink), 7msh (brown)) and ADP•VO_4_ (PDB IDs: 7msm (red), 7msz (salmon red)) bound 70S ribosome complexes (red), PTC/NPET residues in *E. coli* EttA-bound complex are shown in yellow surface. The middle panel shows a zoomed in view of the EttA and tRNA from the complexes. The right panel shows zoomed in view of the PTC showing the two EttA-bound 70S ICs from *E. coli* and the three *Mtb*EttA-bound complexes, the complex with ADPNP (PDB ID: 7msh) has been omitted as the tip of the PtIM has recoiled and the terminal CCA in the acceptor stem of tRNA is not modelled. The view in the right panel is rotated relative to left and middle to present a clearer view of the PTC comparison. **(b)** Zoomed in view of the PTC showing comparisons of the four *Mtb*EttA bound complexes to the reference *E. coli* EttA-bound 70S IC, in addition to the comparison of the two *E. coli* EttA-bound 70S ICs (rows 1 and 2). Row 3 shows comparisons of the deacylated tRNA (tRNA^Val^) containing EttA- bound PRE^-A^_Val_ compared to classical tRNA-containing reference (PDB ID: 4v9d) and to the reference EttA-bound 70S IC, and a comparison of the two reference structures.

**Figure S10.**
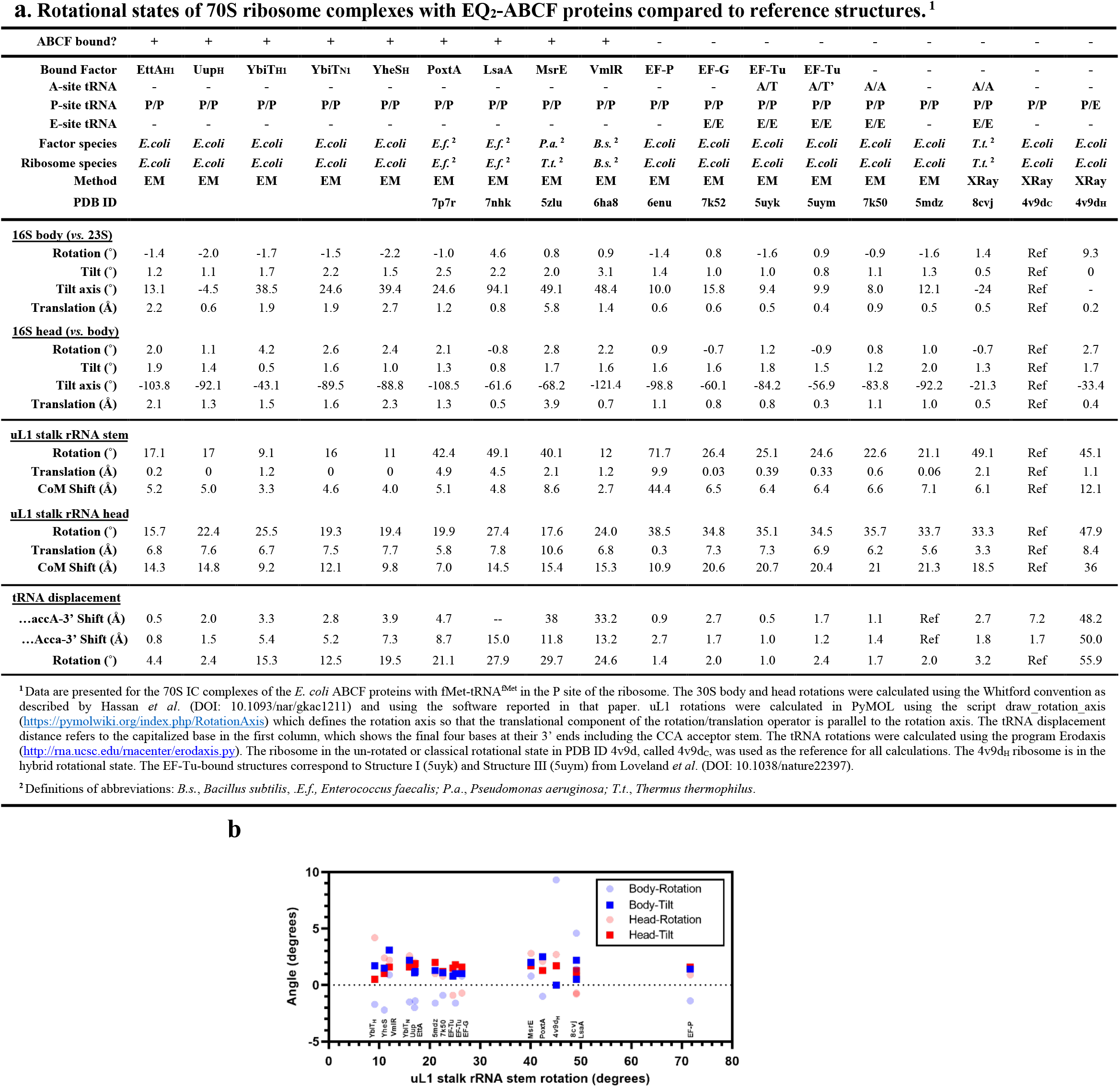
Rotational states of 70S ribosome complexes. **(a)** Table showing the intersubunit orientations, P-site tRNA configurations, and uL1 stalk conformations of the ribosome complexes with *E. coli* ABCFs, ARE ABCFs with PtIMs of varying length (PDB IDs: 7p7r, 7nhk, 5zlu, 6ha8), Elongation-Factor-bound complexes (PDB IDs: 6enu, 7k52, 5uyk, 5uym), and reference ribosomes (PDB IDs: 7k50, 5mdz, 8cvj, 4v9d). Data are shown for the *E. coli* EQ_2_-ABCF proteins in 70S IC complexes, but the values are similar for EQ_2_-EttA and EQ_2_-YbiT in 70S EC (PRE^-A^_Val_) complexes. **(b)** Graph showing the values of the intersubunit orientation and uL1 stalk conformation in the ribosome complexes listed in Table **a**.

**Figure S11.**
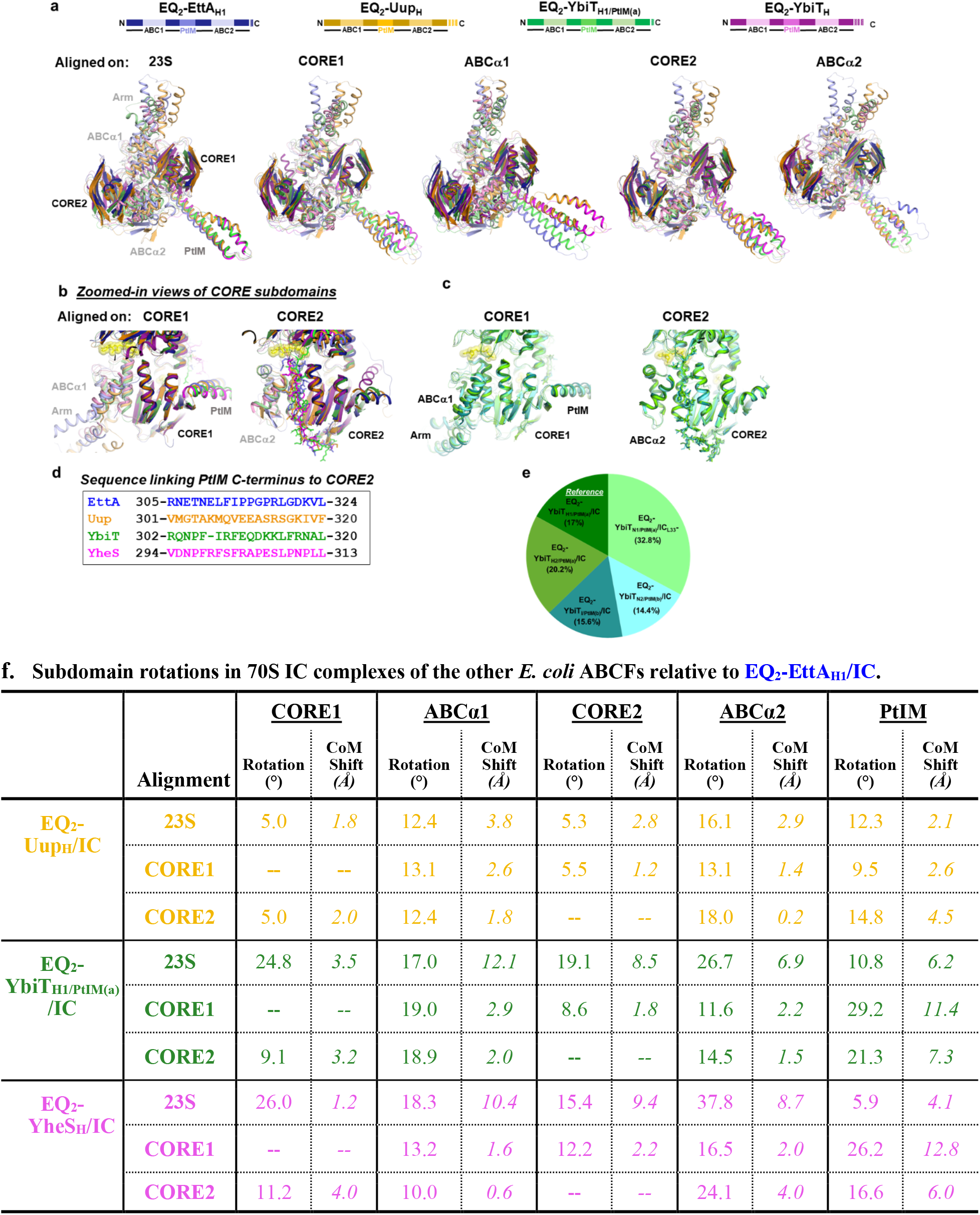
Binding geometries of the E. coli ABCF paralogs in the E site of 70S ICs relative to EQ_2_-EttA_H1_/IC. **(a)** Orientation of the subdomains in *E. coli* EQ2-ABCFs bound to 70S ICs relative to EQ_2_-EttA_H1_ in its 70S ribosome-bound form. The ABCFs used for comparisons are derived from the complexes as mentioned above the bars. The bars depict the color coding adopted for each of the ABCF subdomains, which is dark shade for the CORE, intermediate for PtIM and lighter shade for the ABCα subdomain and the Arm region. The alignments are done on (from left to right) 23S rRNA of the 70S IC and ABCF subdomains CORE1, ABCα1, CORE2, ABCα2. **(b)** Zoomed in view of the four *E. coli* ABCFs in their 70S IC-bound states, aligned on CORE1 and CORE2 subdomain of EQ_2_-EttA. Loop segment linking the base of the PtIM to CORE2 is shown in sticks format and the ATPs from the EQ_2_-EttA are shown as yellow spheres. **(c)** Zoomed in view of the five YbiT-bound 70S ICs aligned on CORE1 and CORE2 of YbiT from the reference complex, EQ_2_-YbiT_H1/PtIM(a)_/IC. **(d)** Sequence alignment in the loop region linking the C-terminus of the PtIM to CORE2 from the four *E. coli* ABCFs. **(e)** Pie chart from **Fig. 8** used here to highlight the color codes used in **c** for YbiTs from the different 70S IC-bound subsets. **(f)** Table has the magnitude of subdomain rotation and shift in center of mass (CoM) of the *E. coli* EQ_2_- ABCF paralogs bound to 70S ICs relative to EQ_2_-EttA in EQ_2_-EttA_H1_/IC. The EQ_2_-ABCFs are aligned on 23S rRNA of the 70S ICs and the CORE1 and CORE2 subdomains of the ABCFs.

**Figure S12.**
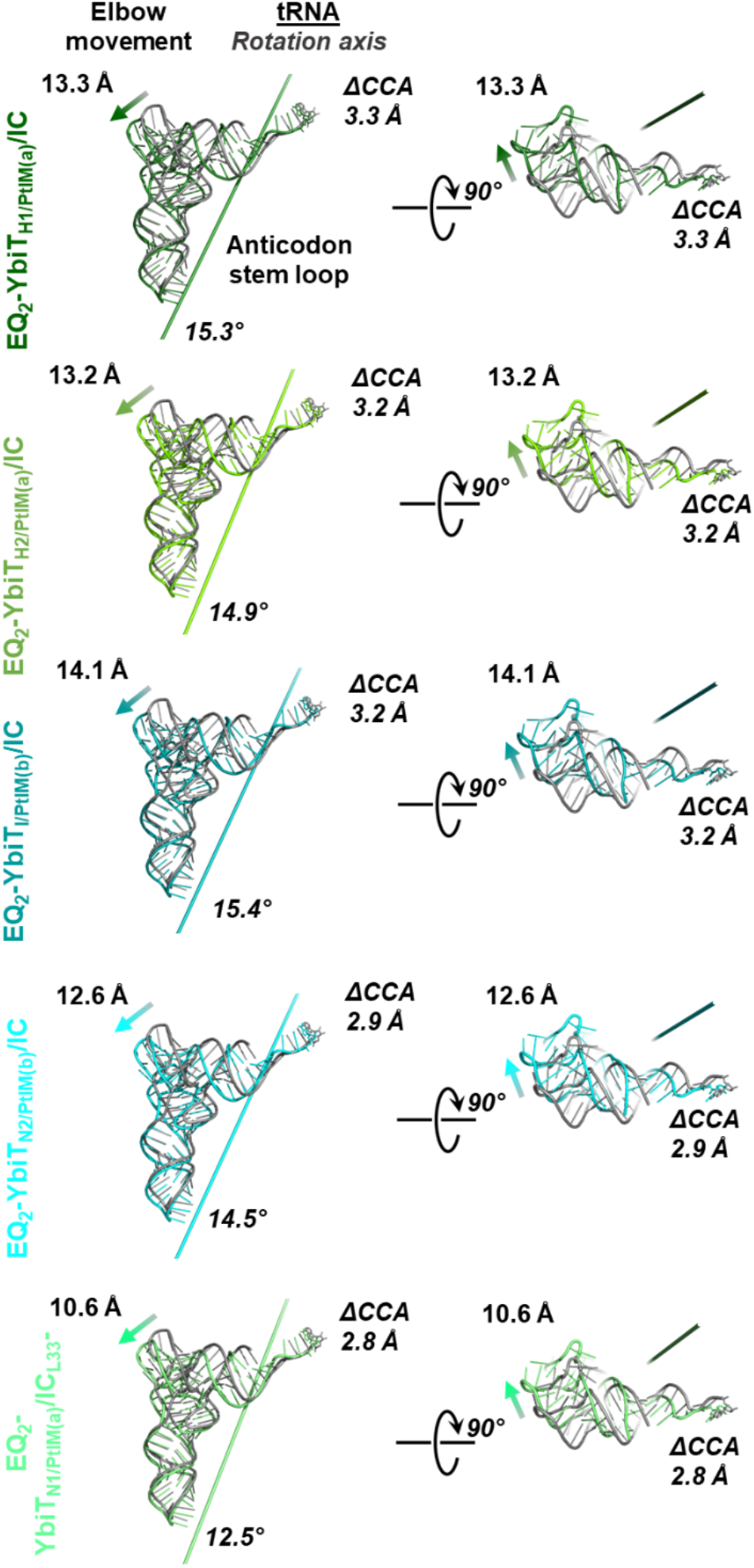
Comparison of the P-site tRNA binding geometry in the different YbiT-bound 70S IC reconstructions. Two orthogonal views showing movement of the P-site tRNA in the five classes of the EQ2-YbiT-bound 70S ICs relative to classical tRNA configuration from the reference ribosome (PDB ID 4v9d). The line running across the length of the tRNA represents the axis of rotation with the magnitude of rotation (°) at bottom right in the left panel. ΔCCA shows the shift in terminal A in the acceptor stem while the arrow at top left of each of the tRNAs shows the direction of elbow movement and value in (Å) shows its magnitude relative to initiator tRNA from the reference ribosome (PDB ID 5mdz). Panels on the right side are 90° rotated view of the images on the left.

**Figure S13.**
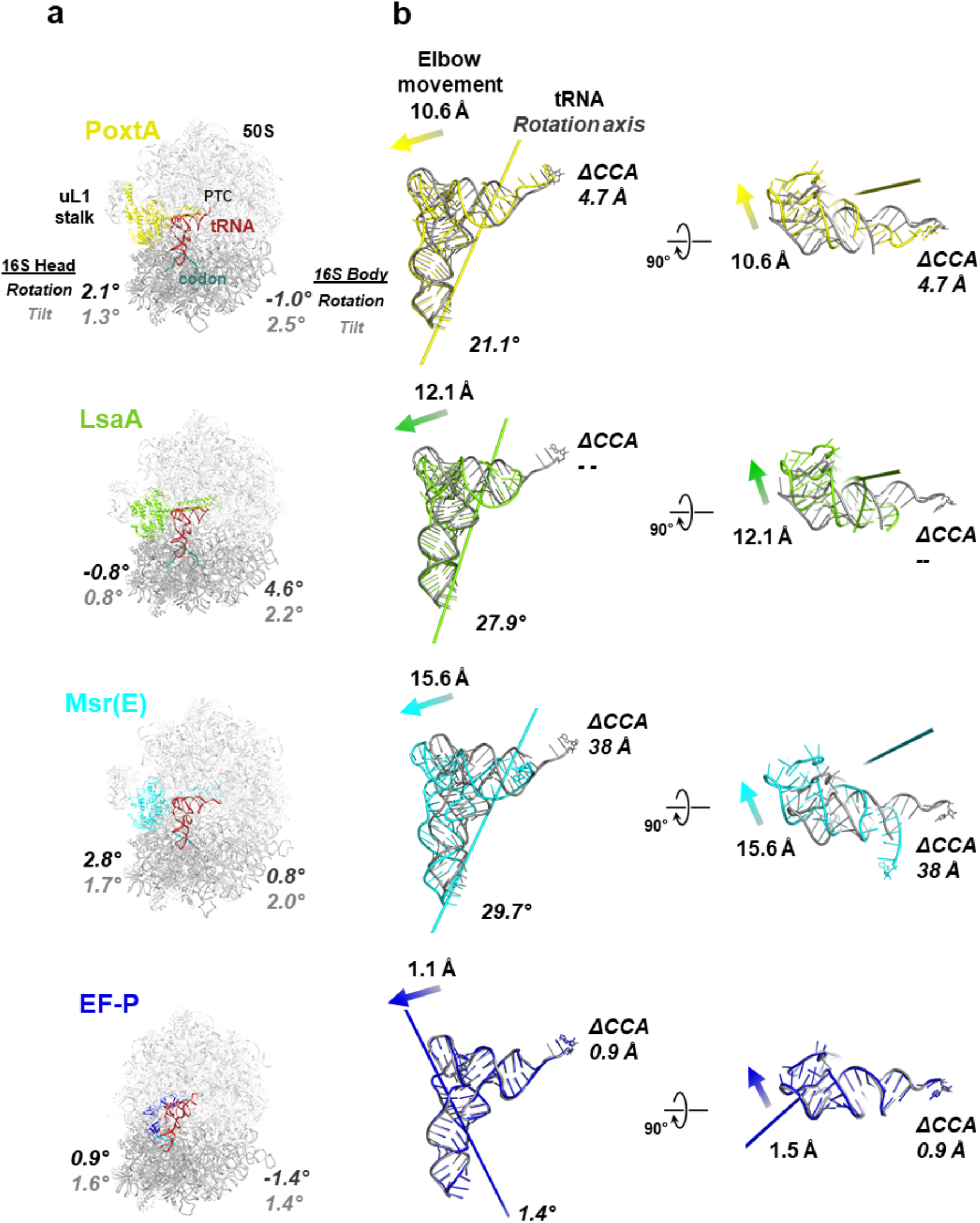
Comparison of the P-site tRNA binding geometry in Elongation Factor P (EF-P) and additional ARE ABCF complexes with 70S ribosomes. **(a)** Cartoon representation of ARE ABCFs PoxtA (PDB ID 7p7r), LsaA (PDB ID 7nhk), MsrE (PDB ID 5zlu), and EF-P (PDB ID 6enu)-bound complexes. The values for intersubunit orientations are mentioned adjacent to each of the complexes. **(b)** Two orthogonal views of the P-site tRNA from each of the complexes shown in the same color as the E-site factors in **a**. The rotation axis of the P-site tRNA relative to the classical tRNA configuration in the reference ribosome (PDB ID 4v9d) are depicted by the line running through the tRNA with the magnitude of rotation (°) at bottom right. The displacement in CCA and movement in elbow region of the P-site tRNA is determined relative to the P-site tRNA (shown in gray) containing 70S ICs of each without the factor at E site (PDB IDs: 7p7u for PoxtA and LsaA, 4v5c for MsrE, 6enf for the tRNA from EF-P bound complex).

**Figure S14.**
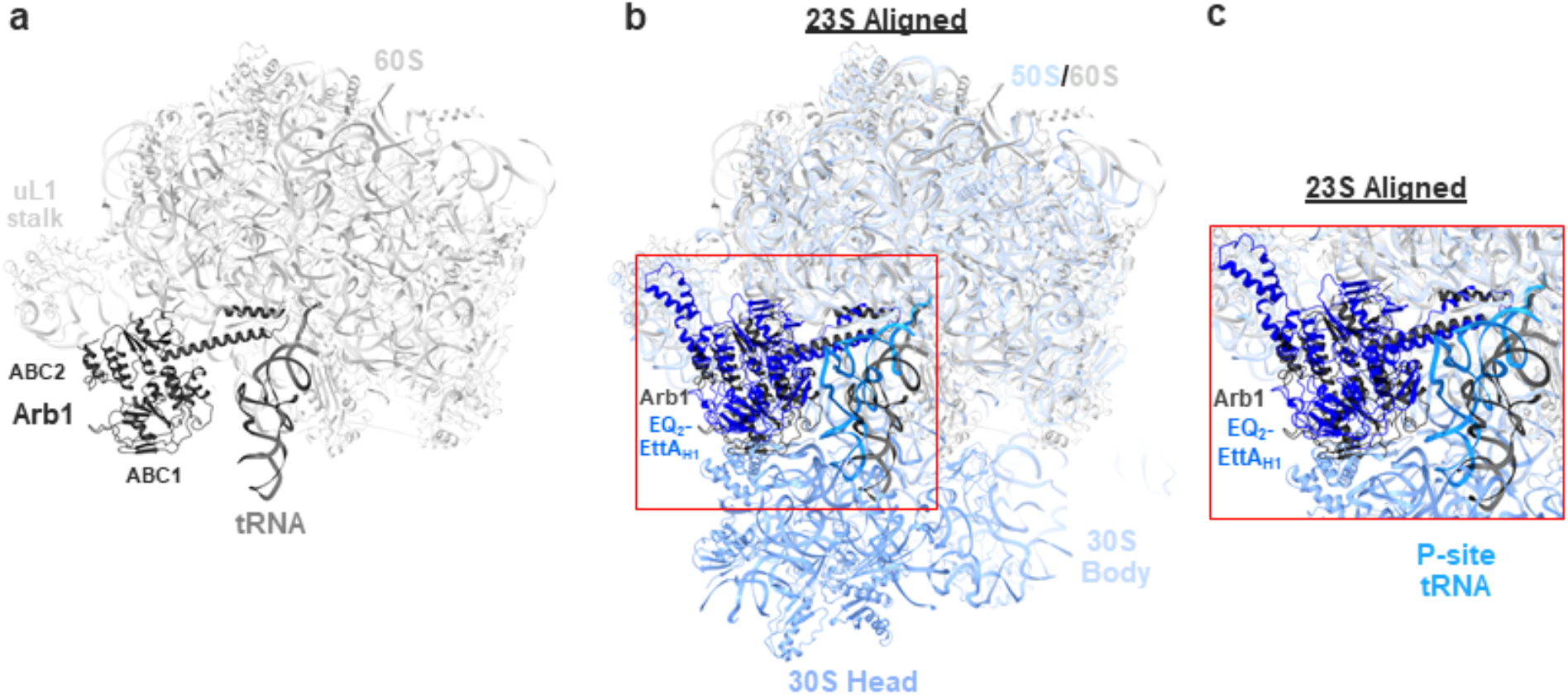
Cryo-EM structure of a yeast ribosome rescue complex shows the PtIM of the ABCF family member Arb1 makes equivalent large (60S) subunit and P-site tRNA contacts as the E. coli ABCF paralogs and the ARE ABCFs of known structure. **(a)** Cartoon representation of the Arb1, ABCF from *S. cerevisiae* in complex with 60S subunit and P-site tRNA (PDB ID 6r84), shown in dark, light and medium shades of gray respectively. **(b)** Arb1 bound P-site tRNA/60S subunit complex superimposed on the EQ_2_-EttA_H1_/IC (30S subunit is shown in dark and 50S subunit is shown in light blue, EttA is shown in blue and P-tRNA is shown in marine blue) aligned on its 23S rRNA. **(c)** Zoomed-in view of the region in inset in **b**.

**Figure S15.**
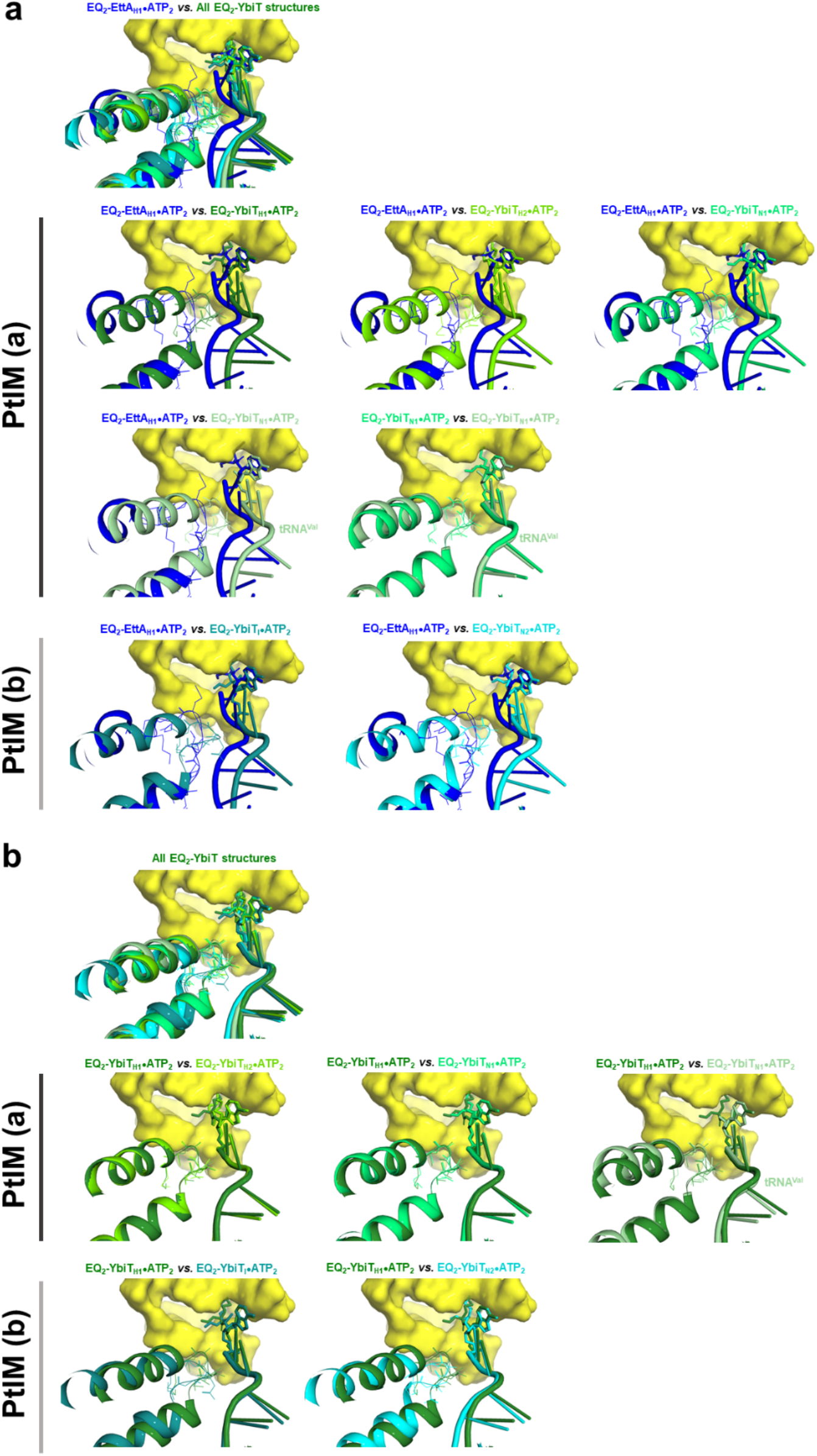
Comparison of PTC stereochemistry observed in YbiT-bound 70S ribosome complexes. **(a)** *E. coli* EQ_2_-YbiT bound ICs and ECs (PRE^-A^_Val_ complexes) compared to the EQ_2_-EttA_H1_/IC (first panel), the residues in the PTC from the EttA-bound 70S IC are shown as yellow surface. The PtIM and tRNA in EQ_2_-EttA are shown in blue and those in EQ_2_-YbiT are shown in shades of green, cyan, and teal. Lower panels show individual YbiT-bound ICs and ECs compared to the EQ_2_-EttA_H1_/IC. PtIM(a) and PtIM(b) indicate the two conformations of PtIM displayed by the YbiTs in the 70S ribosome-bound complexes. EQ_2_-YbiT_N1_/EC is also compared to the EQ_2_-YbiT_N1_/IC (row 3, right) since the YbiT in the two complexes exhibit similar features at conformational level and on the ribosome (missing bL33 density) and the tRNA rotation. **(b)** *E. coli* EQ_2_-YbiT bound ICs and ECs compared to the EQ_2_-YbiT_H1_/IC (first panel) and the lower panels show comparison of each of the complexes individually.

**Figure S16.**
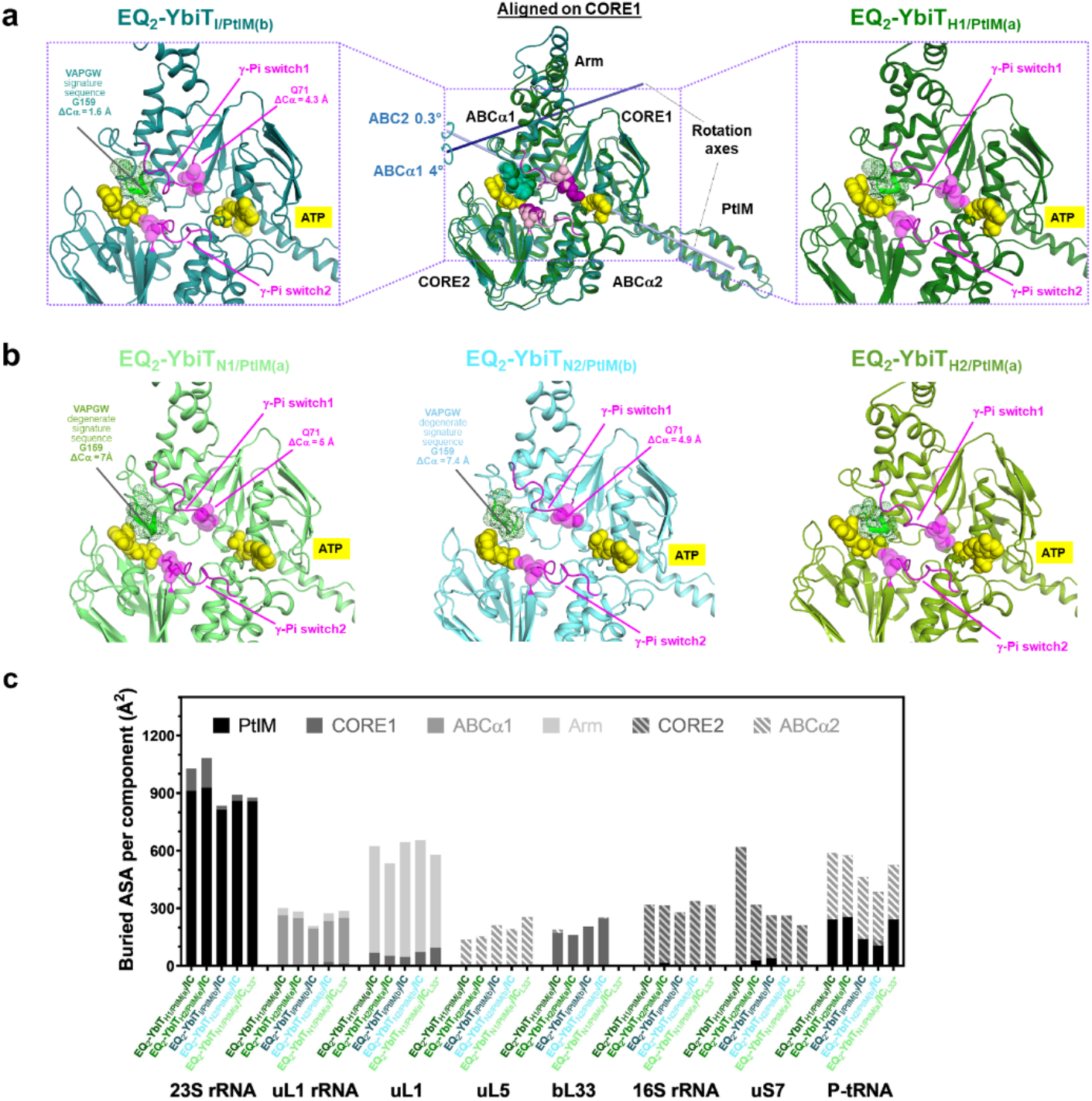
Cryo-EM density and binding geometry of additional YbiT conformational classes. **(a)** Cartoon representation of the YbiTs in Hydrolytic and Intermediate conformations aligned on CORE1. The axes of rotation of the ABCα1 and ABC2 subdomains in the intermediate state relative to the reference hydrolytic state are shown as navy blue and light blue rods. The ATP molecules are shown as yellow spheres, glutamate from γ-Pi switch in magenta and the degenerate signature sequence in ABC1 (VAPGW) are shown as green spheres, in the individual panels. In the superimposed image glutamate from Intermediate conformation is shown as light pink and residues in signature sequence are shown in teal spheres, which are colored in deep violet and green spheres for the Hydrolytic conformation. The left and the right panels show the zoomed in view of the individual YbiTs from the inset showing the glutamate in γ-Pi switch and the ATP molecules along with the signature sequence in ABC1 from the Non-hydrolytic and Hydrolytic conformations respectively. The ΔCα values are the linear displacement of G159 in the ABC1 signature sequence and the Q71 in ABC1 γ-Pi switch from the Intermediate conformation of YbiT in teal relative to the reference Hydrolytic conformation shown in dark green. **(b)** Zoomed-in view of the remaining YbiT conformations classified in the EQ_2_-YbiT/IC dataset, showing YbiT Non-hydrolytic conformation in light green, the second conformation with YbiT in Non-hydrolytic conformation in cyan and the second conformation for YbiT hydrolytic state in chartreuse green. The glutamate residues are shown in magenta spheres, ATP as yellow and the degenerate signature sequence at ATP-binding site 2 is shown as green dotted spheres. The shift in the Cα of G159 and Q71 in the two Non-hydrolytic conformations relative to the reference Hydrolytic conformation are indicated as well. **(c)** Bar graphs showing buried surface area between the subdomains in the different classes of *E. coli* EQ_2_-YbiT bound IC and the ribosomal elements and the P-site tRNA.

**Figure S17.**
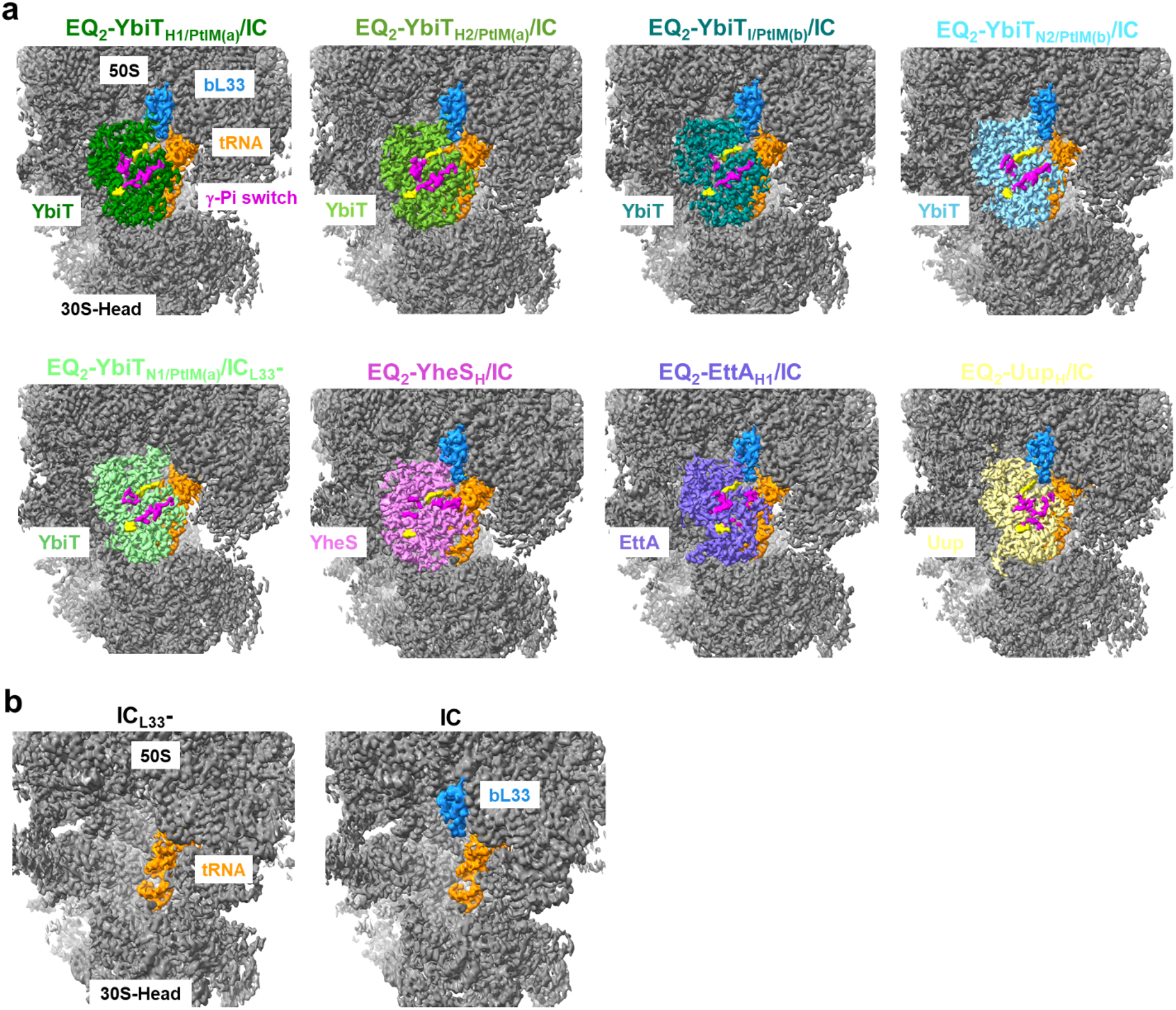
Cryo-EM density and binding geometry of additional YbiT conformational classes and the other E. coli ABCF paralogs in the vicinity of ribosomal protein bL33. **(a)** Density maps of all four *E. coli* ABCF bound ICs highlighting the density for the ABCFs in different shades, P-site tRNA in orange and the ribosomal protein from 50S subunit, bL33, in marine blue, **(b)** Density maps of the ICs; the left panel has the 50S subunit protein bL33 missing while the right panel shows the density map of the class with bL33 (marine blue).

## References

1 Higgins, C. F. ABC transporters: from microorganisms to man. Annu Rev Cell Biol 8, 67–113, doi:10.1146/annurev.cb.08.110192.000435 (1992).

2 Higgins, C. F., Hiles, I. D., Salmond, G. P., Gill, D. R., Downie, J. A., Evans, I. J., Holland, I. B., Gray, L., Buckel, S. D., Bell, A. W. &, et al. A family of related ATP-binding subunits coupled to many distinct biological processes in bacteria. Nature 323, 448–450, doi:10.1038/323448a0 (1986).

3 Jaciuk, M., Nowak, E., Skowronek, K., Tanska, A. & Nowotny, M. Structure of UvrA nucleotide excision repair protein in complex with modified DNA. Nat Struct Mol Biol 18, 191–197, doi:10.1038/nsmb.1973 (2011).

4 Lammens, K., Bemeleit, D. J., Mockel, C., Clausing, E., Schele, A., Hartung, S., Schiller, C. B., Lucas, M., Angermuller, C., Soding, J., Strasser, K. & Hopfner, K. P. The Mre11:Rad50 structure shows an ATP-dependent molecular clamp in DNA double-strand break repair. Cell 145, 54–66, doi:10.1016/j.cell.2011.02.038 (2011).

5 Hopfner, K. P., Karcher, A., Shin, D. S., Craig, L., Arthur, L. M., Carney, J. P. & Tainer, J. A. Structural biology of Rad50 ATPase: ATP-driven conformational control in DNA double- strand break repair and the ABC-ATPase superfamily. Cell 101, 789–800, doi:10.1016/s0092-8674(00)80890-9 (2000).

6 Nachin, L., Loiseau, L., Expert, D. & Barras, F. SufC: an unorthodox cytoplasmic ABC/ATPase required for [Fe-S] biogenesis under oxidative stress. Embo J 22, 427–437 (2003).

7 Amstrup, S. K., Ong, S. C., Sofos, N., Karlsen, J. L., Skjerning, R. B., Boesen, T., Enghild, J. J., Hove-Jensen, B. & Brodersen, D. E. Structural remodelling of the carbon-phosphorus lyase machinery by a dual ABC ATPase. Nat Commun 14, 1001, doi:10.1038/s41467-023-36604-y (2023).

8 Boël, G., Smith, P. C., Ning, W., Englander, M. T., Chen, B., Hashem, Y., Testa, A. J., Fischer, J. J., Wieden, H.-J., Frank, J., Gonzalez, R. L. & Hunt, J. F. The ABC-F protein EttA gates ribosome entry into the translation elongation cycle. Nature Structural & Molecular Biology 21, 143–151, doi:10.1038/nsmb.2740 (2014).

9 Koberska, M., Vesela, L., Vimberg, V., Lenart, J., Vesela, J., Kamenik, Z., Janata, J. & Balikova Novotna, G. Beyond Self-Resistance: ABCF ATPase LmrC Is a Signal-Transducing Component of an Antibiotic-Driven Signaling Cascade Accelerating the Onset of Lincomycin Biosynthesis. mBio 12, e0173121, doi:10.1128/mBio.01731-21 (2021).

10 Pisarev, A. V., Skabkin, M. A., Pisareva, V. P., Skabkina, O. V., Rakotondrafara, A. M., Hentze, M. W., Hellen, C. U. & Pestova, T. V. The role of ABCE1 in eukaryotic posttermination ribosomal recycling. Mol Cell 37, 196–210, doi:10.1016/j.molcel.2009.12.034 (2010).

11 Tyzack, J. K., Wang, X., Belsham, G. J. & Proud, C. G. ABC50 interacts with eukaryotic initiation factor 2 and associates with the ribosome in an ATP-dependent manner. J Biol Chem 275, 34131–34139 (2000).

12 Kasari, V., Pochopien, A. A., Margus, T., Murina, V., Turnbull, K., Zhou, Y., Nissan, T., Graf, M., Nováček, J., Atkinson, G. C., Johansson, M. J. O., Wilson, D. N. & Hauryliuk, V. A role for the Saccharomyces cerevisiae ABCF protein New1 in translation termination/recycling. Nucleic Acids Research 47, 8807–8820, doi:10.1093/nar/gkz600 (2019).

13 Sharkey, L. K. R. & O’Neill, A. J. Antibiotic Resistance ABC-F Proteins: Bringing Target Protection into the Limelight. ACS Infect Dis 4, 239–246, doi:10.1021/acsinfecdis.7b00251 (2018).

14 Sharkey, L. K., Edwards, T. A. & O’Neill, A. J. ABC-F Proteins Mediate Antibiotic Resistance through Ribosomal Protection. MBio 7, e01975, doi:10.1128/mBio.01975-15 (2016).

15 Wilson, D. N., Hauryliuk, V., Atkinson, G. C. & O’Neill, A. J. Target protection as a key antibiotic resistance mechanism. Nat Rev Microbiol 18, 637–648, doi:10.1038/s41579-020-0386-z (2020).

16 Murina, V., Kasari, M., Takada, H., Hinnu, M., Saha, C. K., Grimshaw, J. W., Seki, T., Reith, M., Putrins, M., Tenson, T., Strahl, H., Hauryliuk, V. & Atkinson, G. C. ABCF ATPases Involved in Protein Synthesis, Ribosome Assembly and Antibiotic Resistance: Structural and Functional Diversification across the Tree of Life. J Mol Biol 431, 3568–3590, doi:10.1016/j.jmb.2018.12.013 (2019).

17 Linton, K. J. & Higgins, C. F. The Escherichia coli ATP-binding cassette (ABC) proteins. Mol Microbiol 28, 5–13, doi:10.1046/j.1365-2958.1998.00764.x (1998).

18 Hardie, D. G., Scott, J. W., Pan, D. A. & Hudson, E. R. Management of cellular energy by the AMP-activated protein kinase system. FEBS Lett 546, 113–120, doi:10.1016/s0014-5793(03)00560-x (2003).

19 Buckstein, M. H., He, J. & Rubin, H. Characterization of Nucleotide Pools as a Function of Physiological State in Escherichia coli. Journal of Bacteriology 190, 718–726, doi:10.1128/jb.01020-07 (2008).

20 Crowe-McAuliffe, C., Murina, V., Turnbull, K. J., Kasari, M., Mohamad, M., Polte, C., Takada, H., Vaitkevicius, K., Johansson, J., Ignatova, Z., Atkinson, G. C., O’Neill, A. J., Hauryliuk, V. & Wilson, D. N. Structural basis of ABCF-mediated resistance to pleuromutilin, lincosamide, and streptogramin A antibiotics in Gram-positive pathogens. Nat Commun 12, 3577, doi:10.1038/s41467-021-23753-1 (2021).

21 Crowe-McAuliffe, C., Murina, V., Turnbull, K. J., Huch, S., Kasari, M., Takada, H., Nersisyan, L., Sundsfjord, A., Hegstad, K., Atkinson, G. C., Pelechano, V., Wilson, D. N. & Hauryliuk, V. Structural basis for PoxtA-mediated resistance to phenicol and oxazolidinone antibiotics. Nat Commun 13, 1860, doi:10.1038/s41467-022-29274-9 (2022).

22 Crowe-McAuliffe, C., Graf, M., Huter, P., Takada, H., Abdelshahid, M., Novacek, J., Murina, V., Atkinson, G. C., Hauryliuk, V. & Wilson, D. N. Structural basis for antibiotic resistance mediated by the Bacillus subtilis ABCF ATPase VmlR. Proc Natl Acad Sci U S A 115, 8978–8983, doi:10.1073/pnas.1808535115 (2018).

23 Mohamad, M., Nicholson, D., Saha, Chayan K., Hauryliuk, V., Edwards, Thomas A., Atkinson, Gemma C., Ranson, Neil A. & O’Neill, Alex J. Sal-type ABC-F proteins: intrinsic and common mediators of pleuromutilin resistance by target protection in staphylococci. Nucleic Acids Research 50, 2128–2142, doi:10.1093/nar/gkac058 %J Nucleic Acids Research (2022).

24 Obana, N., Takada, H., Crowe-McAuliffe, C., Iwamoto, M., Egorov, A. A., Wu, K. J. Y., Chiba, S., Murina, V., Paternoga, H., Tresco, B. I. C., Nomura, N., Myers, A. G., Atkinson, G. C., Wilson, D. N. & Hauryliuk, V. Genome-encoded ABCF factors implicated in intrinsic antibiotic resistance in Gram-positive bacteria: VmlR2, Ard1 and CplR. Nucleic Acids Res, doi:10.1093/nar/gkad193 (2023).

25 Chen, B., Boël, G., Hashem, Y., Ning, W., Fei, J., Wang, C., Gonzalez, R. L., Hunt, J. F. & Frank, J. EttA regulates translation by binding the ribosomal E site and restricting ribosome- tRNA dynamics. Nature Structural & Molecular Biology 21, 152–159, doi:10.1038/nsmb.2741 (2014).

26 Cui, Z., Li, X., Shin, J., Gamper, H., Hou, Y. M., Sacchettini, J. C. & Zhang, J. Interplay between an ATP-binding cassette F protein and the ribosome from Mycobacterium tuberculosis. Nat Commun 13, 432, doi:10.1038/s41467-022-28078-1 (2022).

27 Su, W., Kumar, V., Ding, Y., Ero, R., Serra, A., Lee, B. S. T., Wong, A. S. W., Shi, J., Sze, S. K., Yang, L. & Gao, Y. G. Ribosome protection by antibiotic resistance ATP-binding cassette protein. Proc Natl Acad Sci U S A 115, 5157–5162, doi:10.1073/pnas.1803313115 (2018).

28 Su, T., Izawa, T., Thoms, M., Yamashita, Y., Cheng, J., Berninghausen, O., Hartl, F. U., Inada, T., Neupert, W. & Beckmann, R. Structure and function of Vms1 and Arb1 in RQC and mitochondrial proteome homeostasis. Nature 570, 538–542, doi:10.1038/s41586-019-1307-z (2019).

29 Finn, R. D., Bateman, A., Clements, J., Coggill, P., Eberhardt, R. Y., Eddy, S. R., Heger, A., Hetherington, K., Holm, L., Mistry, J., Sonnhammer, E. L., Tate, J. & Punta, M. Pfam: the protein families database. Nucleic Acids Res 42, D222–230, doi:10.1093/nar/gkt1223 (2014).

30 Ero, R., Kumar, V., Su, W. & Gao, Y. G. Ribosome protection by ABC-F proteins-Molecular mechanism and potential drug design. Protein Sci 28, 684–693, doi:10.1002/pro.3589 (2019).

31 Zhang, C., Liu, L., Zhang, P., Cui, J., Qin, X., Ma, L., Han, K., Wang, Z., Wang, S., Ding, S. & Shen, Z. Characterization of a Novel Gene, srpA, Conferring Resistance to Streptogramin A, Pleuromutilins, and Lincosamides in Streptococcus suis. Engineering 9, 85–94, doi:10.1016/j.eng.2020.12.015 (2022).

32 Lenart, J., Vimberg, V., Vesela, L., Janata, J. & Balikova Novotna, G. Detailed mutational analysis of Vga(A) interdomain linker: implication for antibiotic resistance specificity and mechanism. Antimicrob Agents Chemother 59, 1360–1364, doi:10.1128/AAC.04468-14 (2015).

33 Fostier, C. R., Monlezun, L., Ousalem, F., Singh, S., Hunt, J. F. & Boël, G. ABC-F translation factors: from antibiotic resistance to immune response. FEBS Letters 595, 675–706, doi:10.1002/1873-3468.13984 (2020).

34 Ousalem, F., Singh, S., Chesneau, O., Hunt, J. F. & Boël, G. ABC-F proteins in mRNA translation and antibiotic resistance. Research in Microbiology 170, 435–447, doi:10.1016/j.resmic.2019.09.005 (2019).

35 Moody, J. E., Millen, L., Binns, D., Hunt, J. F. & Thomas, P. J. Cooperative, ATP-dependent association of the nucleotide binding cassettes during the catalytic cycle of ATP-binding cassette transporters. J Biol Chem 277, 21111–21114, doi:10.1074/jbc.C200228200 (2002).

36 Oldham, M. L. & Chen, J. Snapshots of the maltose transporter during ATP hydrolysis. Proc Natl Acad Sci U S A 108, 15152–15156, doi:10.1073/pnas.1108858108 (2011).

37 Rees, D. C., Johnson, E. & Lewinson, O. ABC transporters: the power to change. Nat Rev Mol Cell Biol 10, 218–227, doi:10.1038/nrm2646 (2009).

38 Smith, P. C., Karpowich, N., Millen, L., Moody, J. E., Rosen, J., Thomas, P. J. & Hunt, J. F. ATP binding to the motor domain from an ABC transporter drives formation of a nucleotide sandwich dimer. Mol Cell 10, 139–149, doi:10.1016/s1097-2765(02)00576-2 (2002).

39 Timachi, M. H., Hutter, C. A., Hohl, M., Assafa, T., Bohm, S., Mittal, A., Seeger, M. A. & Bordignon, E. Exploring conformational equilibria of a heterodimeric ABC transporter. Elife 6, doi:10.7554/eLife.20236 (2017).

40 Hohl, M., Hurlimann, L. M., Bohm, S., Schoppe, J., Grutter, M. G., Bordignon, E. & Seeger, M. A. Structural basis for allosteric cross-talk between the asymmetric nucleotide binding sites of a heterodimeric ABC exporter. Proc Natl Acad Sci U S A 111, 11025–11030, doi:10.1073/pnas.1400485111 (2014).

41 Schmitt, L. & Tampe, R. Structure and mechanism of ABC transporters. Current opinion in structural biology 12, 754–760 (2002).

42 Oldham, M. L. & Chen, J. Crystal structure of the maltose transporter in a pretranslocation intermediate state. Science (New York, N.Y 332, 1202–1205, doi:10.1126/science.1200767 (2011).

43 van der Does, C. & Tampe, R. How do ABC transporters drive transport? Biol Chem 385, 927–933 (2004).

44 Yuan, Y. R., Blecker, S., Martsinkevich, O., Millen, L., Thomas, P. J. & Hunt, J. F. The crystal structure of the MJ0796 ATP-binding cassette. Implications for the structural consequences of ATP hydrolysis in the active site of an ABC transporter. J Biol Chem 276, 32313–32321, doi:10.1074/jbc.M100758200 (2001).

45 Verdon, G., Albers, S. V., Dijkstra, B. W., Driessen, A. J. & Thunnissen, A. M. Crystal structures of the ATPase subunit of the glucose ABC transporter from Sulfolobus solfataricus: nucleotide-free and nucleotide-bound conformations. J Mol Biol 330, 343–358 (2003).

46 Chen, J., Lu, G., Lin, J., Davidson, A. L. & Quiocho, F. A. A tweezers-like motion of the ATP- binding cassette dimer in an ABC transport cycle. Mol Cell 12, 651–661, doi:10.1016/j.molcel.2003.08.004 (2003).

47 Ambudkar, S. V., Kim, I. W. & Sauna, Z. E. The power of the pump: mechanisms of action of P-glycoprotein (ABCB1). Eur J Pharm Sci 27, 392–400, doi:10.1016/j.ejps.2005.10.010 (2006).

48 Chen, J. Molecular mechanism of the Escherichia coli maltose transporter. Current opinion in structural biology 23, 492–498, doi:10.1016/j.sbi.2013.03.011 (2013).

49 Hwang, T. C. & Sheppard, D. N. Gating of the CFTR Cl- channel by ATP-driven nucleotide- binding domain dimerisation. J Physiol 587, 2151–2161 (2009).

50 Dawson, R. J., Hollenstein, K. & Locher, K. P. Uptake or extrusion: crystal structures of full ABC transporters suggest a common mechanism. Mol Microbiol 65, 250–257 (2007).

51 Kim, D. M., Zheng, H., Huang, Y. J., Montelione, G. T. & Hunt, J. F. ATPase active-site electrostatic interactions control the global conformation of the 100 kDa SecA translocase. J Am Chem Soc 135, 2999–3010, doi:10.1021/ja306361q (2013).

52 Zhu, L., Kim, J., Leng, K., Ramos, J. E., Kinz-Thompson, C. D., Karpowich, N. K., Ruben L. Gonzalez, J. & Hunt, J. F. Realtime observation of ATP-driven single B_12_ molecule translocation through BtuCD-F. bioRxiv, 2022.2012.2002.518935, doi:10.1101/2022.12.02.518935 (2022).

53 Zhu, L., Kim, J., Leng, K., Ramos, J. E., Kinz-Thompson, C. D., Karpowich, N. K., Ruben L. Gonzalez, J. & Hunt, J. F. Mechanistic implications of the interaction of the soluble substrate- binding protein with a type II ABC importer. bioRxiv, 2022.2012.2002.518933, doi:10.1101/2022.12.02.518933 (2022).

54 Arbing, M. A., Handelman, S. K., Kuzin, A. P., Verdon, G., Wang, C., Su, M., Rothenbacher, F. P., Abashidze, M., Liu, M., Hurley, J. M., Xiao, R., Acton, T., Inouye, M., Montelione, G. T., Woychik, N. A. & Hunt, J. F. Crystal Structures of Phd-Doc, HigA, and YeeU Establish Multiple Evolutionary Links between Microbial Growth-Regulating Toxin-Antitoxin Systems. Structure 18, 996–1010, doi:10.1016/j.str.2010.04.018 (2010).

55 van der Does, C., Presenti, C., Schulze, K., Dinkelaker, S. & Tampe, R. Kinetics of the ATP hydrolysis cycle of the nucleotide-binding domain of Mdl1 studied by a novel site-specific labeling technique. J Biol Chem 281, 5694–5701, doi:10.1074/jbc.M511730200 (2006).

56 Janas, E., Hofacker, M., Chen, M., Gompf, S., Van Der Does, C. & Tampe, R. The ATP Hydrolysis Cycle of the Nucleotide-binding Domain of the Mitochondrial ATP-binding Cassette Transporter Mdl1p. J Biol Chem 278, 26862–26869 (2003).

57 Orelle, C., Dalmas, O., Gros, P., Di Pietro, A. & Jault, J. M. The conserved glutamate residue adjacent to the Walker-B motif is the catalytic base for ATP hydrolysis in the ATP-binding cassette transporter BmrA. J Biol Chem 278, 47002–47008 (2003).

58 Huter, P., Arenz, S., Bock, L. V., Graf, M., Frister, J. O., Heuer, A., Peil, L., Starosta, A. L., Wohlgemuth, I., Peske, F., Novacek, J., Berninghausen, O., Grubmuller, H., Tenson, T., Beckmann, R., Rodnina, M. V., Vaiana, A. C. & Wilson, D. N. Structural Basis for Polyproline-Mediated Ribosome Stalling and Rescue by the Translation Elongation Factor EF- P. Mol Cell 68, 515–527 e516, doi:10.1016/j.molcel.2017.10.014 (2017).

59 Karpowich, N., Martsinkevich, O., Millen, L., Yuan, Y.-R., Dai, P. L., MacVey, K., Thomas, P. J. & Hunt, J. F. Crystal Structures of the MJ1267 ATP Binding Cassette Reveal an Induced- Fit Effect at the ATPase Active Site of an ABC Transporter. Structure 9, 571–586, doi:10.1016/s0969-2126(01)00617-7 (2001).

60 Gouridis, G., Hetzert, B., Kiosze-Becker, K., de Boer, M., Heinemann, H., Nurenberg-Goloub, E., Cordes, T. & Tampe, R. ABCE1 Controls Ribosome Recycling by an Asymmetric Dynamic Conformational Equilibrium. Cell Rep 28, 723–734 e726, doi:10.1016/j.celrep.2019.06.052 (2019).

61 Agirrezabala, X., Liao, H. Y., Schreiner, E., Fu, J., Ortiz-Meoz, R. F., Schulten, K., Green, R. & Frank, J. Structural characterization of mRNA-tRNA translocation intermediates. Proceedings of the National Academy of Sciences 109, 6094–6099, doi:10.1073/pnas.1201288109 (2012).

62 Frank, J. & Agrawal, R. K. A ratchet-like inter-subunit reorganization of the ribosome during translocation. Nature 406, 318–322, doi:10.1038/35018597 (2000).

63 Valle, M., Zavialov, A., Sengupta, J., Rawat, U., Ehrenberg, M. & Frank, J. Locking and unlocking of ribosomal motions. Cell 114, 123–134, doi:10.1016/s0092-8674(03)00476-8 (2003).

64 Kaledhonkar, S., Fu, Z., Caban, K., Li, W., Chen, B., Sun, M., Gonzalez, R. L., Jr. & Frank, J. Late steps in bacterial translation initiation visualized using time-resolved cryo-EM. Nature 570, 400–404, doi:10.1038/s41586-019-1249-5 (2019).

65 Singh, S., Wong, K., Gentry, R. C., Patel, S., Aalberts, D. P., Frank, J., Gonzalez, R. L., Jr., Boël, G., Kinz-Thompson, C. D., Hunt, J. F. The mechanism and physiological profile of translational regulation by the ABCF protein EttA. Manuscript in preparation.

66 Ousalem, F., Ngo, S., Oïffer, T., Nasser, A. O., Monlezun, L., Boël, G. EttA alleviates early ribosome stalling events induced by acidic-residues. Manuscript in preparation.

67 Ousalem, F., Singh, S., Bailey, N. A., Wong, K., Zhu, L., Neky, M. J., Sibindi, C., Fei, J., Gonzalez, R. L., Jr., Boël, G. and Hunt, J. F. Comparative genetic, biochemical and biophysical analyses of the four E. coli ABCF paralogs support multiple functions related to mRNA translation. bioRxiv, doi:10.1101/2023.06.11.543863 (2023).

68 Fei, J., Richard, A. C., Bronson, J. E. & Gonzalez, R. L. Transfer RNA–mediated regulation of ribosome dynamics during protein synthesis. Nature Structural & Molecular Biology 18, 1043–1051, doi:10.1038/nsmb.2098 (2011).

69 Ning, W., Fei, J. & Gonzalez, R. L., Jr. The ribosome uses cooperative conformational changes to maximize and regulate the efficiency of translation. Proc Natl Acad Sci U S A 111, 12073–12078, doi:10.1073/pnas.1401864111 (2014).

70 MacDougall, D. D., Fei, J. & Gonzalez, R. L. in Molecular Machines in Biology Ch. Chapter 6, 93–116 (2011).

71 Punjani, A., Zhang, H. & Fleet, D. J. Non-uniform refinement: adaptive regularization improves single-particle cryo-EM reconstruction. Nature Methods 17, 1214–1221, doi:10.1038/s41592-020-00990-8 (2020).

72 Afonine, P. V., Poon, B. K., Read, R. J., Sobolev, O. V., Terwilliger, T. C., Urzhumtsev, A. & Adams, P. D. Real-space refinement in PHENIX for cryo-EM and crystallography. Acta Crystallogr D Struct Biol 74, 531–544, doi:10.1107/S2059798318006551 (2018).

73 Scheres, S. H. W. RELION: Implementation of a Bayesian approach to cryo-EM structure determination. Journal of Structural Biology 180, 519–530, doi:10.1016/j.jsb.2012.09.006 (2012).

74 Punjani, A., Rubinstein, J. L., Fleet, D. J. & Brubaker, M. A. cryoSPARC: algorithms for rapid unsupervised cryo-EM structure determination. Nature Methods 14, 290–296, doi:10.1038/nmeth.4169 (2017).

75 Fei, J., Kosuri, P., MacDougall, D. D. & Gonzalez, R. L., Jr. Coupling of ribosomal L1 stalk and tRNA dynamics during translation elongation. Mol Cell 30, 348–359, doi:10.1016/j.molcel.2008.03.012 (2008).

76 Frank, J., Gao, H., Sengupta, J., Gao, N. & Taylor, D. J. The process of mRNA-tRNA translocation. Proc Natl Acad Sci U S A 104, 19671–19678, doi:10.1073/pnas.0708517104 (2007).

77 Hassan, A., Byju, S., Freitas, Frederico C., Roc, C., Pender, N., Nguyen, K., Kimbrough, Evelyn M., Mattingly, Jacob M., Gonzalez Jr, Ruben L., de Oliveira, Ronaldo J., Dunham, Christine M. & Whitford, Paul C. Ratchet, swivel, tilt and roll: a complete description of subunit rotation in the ribosome. Nucleic Acids Research 51, 919–934, doi:10.1093/nar/gkac1211 (2023).

78 Dunkle, J. A., Wang, L., Feldman, M. B., Pulk, A., Chen, V. B., Kapral, G. J., Noeske, J., Richardson, J. S., Blanchard, S. C. & Cate, J. H. D. Structures of the Bacterial Ribosome in Classical and Hybrid States of tRNA Binding. Science 332, 981–984, doi:10.1126/science.1202692 (2011).

79 Cornish, P. V., Ermolenko, D. N., Staple, D. W., Hoang, L., Hickerson, R. P., Noller, H. F. & Ha, T. Following movement of the L1 stalk between three functional states in single ribosomes. Proc Natl Acad Sci U S A 106, 2571–2576, doi:10.1073/pnas.0813180106 (2009).

80 Bock, L. V., Blau, C., Vaiana, A. C. & Grubmüller, H. Dynamic contact network between ribosomal subunits enables rapid large-scale rotation during spontaneous translocation. Nucleic Acids Research 43, 6747–6760, doi:10.1093/nar/gkv649 (2015).

81 James, N. R., Brown, A., Gordiyenko, Y. & Ramakrishnan, V. Translational termination without a stop codon. Science 354, 1437–1440, doi:10.1126/science.aai9127 (2016).

82 Cruickshank, D. W. Remarks about protein structure precision. Acta Crystallogr D Biol Crystallogr 55, 583–601, doi:10.1107/s0907444998012645 (1999).

83 Singh, S., Gentry, R. C., Gonzalez, R. L., Jr., Boël, G., Hunt, J. F. YheS binding induces an allosteric conformational change in the peptidyltransferase center of the ribosome that highlights underlying conformational disorder. Manuscript in preparation.

84 Watson, Z. L., Ward, F. R., Meheust, R., Ad, O., Schepartz, A., Banfield, J. F. & Cate, J. H. Structure of the bacterial ribosome at 2 A resolution. Elife 9, doi:10.7554/eLife.60482 (2020).

85 Demo, G., Gamper, H. B., Loveland, A. B., Masuda, I., Carbone, C. E., Svidritskiy, E., Hou, Y. M. & Korostelev, A. A. Structural basis for +1 ribosomal frameshifting during EF-G- catalyzed translocation. Nat Commun 12, 4644, doi:10.1038/s41467-021-24911-1 (2021).

86 Benkovic, S. J. & Hammes-Schiffer, S. A perspective on enzyme catalysis. Science 301, 1196–1202, doi:10.1126/science.1085515 (2003).

87 Robertson, W. R., Dowsett, S. J. & Hardy, S. J. S. Exchange of ribosomal proteins among the ribosomes of Escherichia coli. Molecular and General Genetics MGG 157, 205–214, doi:10.1007/bf00267399 (1977).

88 Murina, V., Kasari, M., Hauryliuk, V. & Atkinson, G. C. Antibiotic resistance ABCF proteins reset the peptidyl transferase centre of the ribosome to counter translational arrest. Nucleic Acids Res 46, 3753–3763, doi:10.1093/nar/gky050 (2018).

89 Tuller, T., Carmi, A., Vestsigian, K., Navon, S., Dorfan, Y., Zaborske, J., Pan, T., Dahan, O., Furman, I. & Pilpel, Y. An evolutionarily conserved mechanism for controlling the efficiency of protein translation. Cell 141, 344–354, doi:10.1016/j.cell.2010.03.031 (2010).

90 Chevance, F. F., Le Guyon, S. & Hughes, K. T. The effects of codon context on in vivo translation speed. PLoS Genet 10, e1004392, doi:10.1371/journal.pgen.1004392 (2014).

91 Vimberg, V., Cavanagh, J. P., Novotna, M., Lenart, J., Nguyen Thi Ngoc, B., Vesela, J., Pain, M., Koberska, M. & Balikova Novotna, G. Ribosome-Mediated Attenuation of vga(A) Expression Is Shaped by the Antibiotic Resistance Specificity of Vga(A) Protein Variants. Antimicrob Agents Chemother 64, doi:10.1128/AAC.00666-20 (2020).

92 Takada, H., Mandell, Z. F., Yakhnin, H., Glazyrina, A., Chiba, S., Kurata, T., Wu, K. J. Y., Tresco, B. I. C., Myers, A. G., Aktinson, G. C., Babitzke, P. & Hauryliuk, V. Expression of Bacillus subtilis ABCF antibiotic resistance factor VmlR is regulated by RNA polymerase pausing, transcription attenuation, translation attenuation and (p)ppGpp. Nucleic Acids Res 50, 6174–6189, doi:10.1093/nar/gkac497 (2022).

93 Fostier, C. R., Ousalem, F., Leroy, E. C., Ngo, S., Soufari, H., Innis, C. A., Hashem, Y. & Boёl, G. Regulation of the macrolide resistance ABC-F translation factor MsrD. bioRxiv, doi:10.1101/2021.11.29.470318 (2022).

94 Artsimovitch, I. & Landick, R. Pausing by bacterial RNA polymerase is mediated by mechanistically distinct classes of signals. Proc Natl Acad Sci U S A 97, 7090–7095, doi:10.1073/pnas.97.13.7090 (2000).

95 Larson, M. H., Mooney, R. A., Peters, J. M., Windgassen, T., Nayak, D., Gross, C. A., Block, S. M., Greenleaf, W. J., Landick, R. & Weissman, J. S. A pause sequence enriched at translation start sites drives transcription dynamics in vivo. Science (New York, N.Y 344, 1042–1047, doi:10.1126/science.1251871 (2014).

96 Arenz, S., Ramu, H., Gupta, P., Berninghausen, O., Beckmann, R., Vazquez-Laslop, N., Mankin, A. S. & Wilson, D. N. Molecular basis for erythromycin-dependent ribosome stalling during translation of the ErmBL leader peptide. Nat Commun 5, 3501, doi:10.1038/ncomms4501 (2014).

97 Min, Y. H., Kwon, A. R., Yoon, E. J., Shim, M. J. & Choi, E. C. Translational attenuation and mRNA stabilization as mechanisms of erm(B) induction by erythromycin. Antimicrob Agents Chemother 52, 1782–1789, doi:10.1128/AAC.01376-07 (2008).

## Methods References

98 Sievers, F., Wilm, A., Dineen, D., Gibson, T. J., Karplus, K., Li, W., Lopez, R., McWilliam, H., Remmert, M., Soding, J., Thompson, J. D. & Higgins, D. G. Fast, scalable generation of high-quality protein multiple sequence alignments using Clustal Omega. Mol Syst Biol 7, 539, doi:10.1038/msb.2011.75 (2011).

99 Robert, X. & Gouet, P. Deciphering key features in protein structures with the new ENDscript server. Nucleic Acids Research 42, W320–W324, doi:10.1093/nar/gku316 (2014).

100 Andersen, K. R., Leksa, N. C. & Schwartz, T. U. Optimized E. coli expression strain LOBSTR eliminates common contaminants from His-tag purification. Proteins 81, 1857–1861, doi:10.1002/prot.24364 (2013).

101 Fei, J., Wang, J., Sternberg, S. H., MacDougall, D. D., Elvekrog, M. M., Pulukkunat, D. K., Englander, M. T. & Gonzalez, R. L., Jr. A highly purified, fluorescently labeled in vitro translation system for single-molecule studies of protein synthesis. Methods in enzymology 472, 221–259, doi:10.1016/S0076-6879(10)72008-5 (2010).

102 Sternberg, S. H., Fei, J., Prywes, N., McGrath, K. A. & Gonzalez, R. L., Jr. Translation factors direct intrinsic ribosome dynamics during translation termination and ribosome recycling. Nat Struct Mol Biol 16, 861–868, doi:10.1038/nsmb.1622 (2009).

103 Antoun, A., Pavlov, M. Y., Lovmar, M. & Ehrenberg, M. How initiation factors tune the rate of initiation of protein synthesis in bacteria. EMBO J 25, 2539–2550, doi:10.1038/sj.emboj.7601140 (2006).

104 MacDougall, D. D. & Gonzalez, R. L., Jr. Translation initiation factor 3 regulates switching between different modes of ribosomal subunit joining. J Mol Biol 427, 1801–1818, doi:10.1016/j.jmb.2014.09.024 (2015).

105 Zheng, S. Q., Palovcak, E., Armache, J.-P., Verba, K. A., Cheng, Y. & Agard, D. A. MotionCor2: anisotropic correction of beam-induced motion for improved cryo-electron microscopy. Nature Methods 14, 331–332, doi:10.1038/nmeth.4193 (2017).

106 Zhang, K. Gctf: Real-time CTF determination and correction. Journal of Structural Biology 193, 1–12, doi:10.1016/j.jsb.2015.11.003 (2016).

107 Zivanov, J., Nakane, T. & Scheres, S. H. W. A Bayesian approach to beam-induced motion correction in cryo-EM single-particle analysis. IUCrJ 6, 5–17, doi:10.1107/S205225251801463X (2019).

108 Zivanov, J., Nakane, T. & Scheres, S. H. W. Estimation of high-order aberrations and anisotropic magnification from cryo-EM data sets in RELION-3.1. IUCrJ 7, 253-267, doi:10.1107/s2052252520000081 (2020).

109 Kucukelbir, A., Sigworth, F. J. & Tagare, H. D. Quantifying the local resolution of cryo-EM density maps. Nature Methods 11, 63–65, doi:10.1038/nmeth.2727 (2013).

110 Demo, G., Svidritskiy, E., Madireddy, R., Diaz-Avalos, R., Grant, T., Grigorieff, N., Sousa, D. & Korostelev, A. A. Mechanism of ribosome rescue by ArfA and RF2. Elife 6, doi:10.7554/eLife.23687 (2017).

111 Pettersen, E. F., Goddard, T. D., Huang, C. C., Couch, G. S., Greenblatt, D. M., Meng, E. C. & Ferrin, T. E. UCSF Chimera--A visualization system for exploratory research and analysis. Journal of Computational Chemistry 25, 1605–1612, doi:10.1002/jcc.20084 (2004).

112 Emsley, P. & Cowtan, K. Coot: model-building tools for molecular graphics. Acta crystallographica 60, 2126–2132, doi:10.1107/S0907444904019158 (2004).

113 Liebschner, D., Afonine, P. V., Baker, M. L., Bunkoczi, G., Chen, V. B., Croll, T. I., Hintze, B., Hung, L. W., Jain, S., McCoy, A. J., Moriarty, N. W., Oeffner, R. D., Poon, B. K., Prisant, M. G., Read, R. J., Richardson, J. S., Richardson, D. C., Sammito, M. D., Sobolev, O. V., Stockwell, D. H., Terwilliger, T. C., Urzhumtsev, A. G., Videau, L. L., Williams, C. J. & Adams, P. D. Macromolecular structure determination using X-rays, neutrons and electrons: recent developments in Phenix. Acta Crystallogr D Struct Biol 75, 861–877, doi:10.1107/S2059798319011471 (2019).

114 Moriarty, N. W., Grosse-Kunstleve, R. W. & Adams, P. D. electronic Ligand Builder and Optimization Workbench (eLBOW): a tool for ligand coordinate and restraint generation. Acta Crystallogr D Biol Crystallogr 65, 1074–1080, doi:10.1107/S0907444909029436 (2009).

115 Afonine, P. V., Klaholz, B. P., Moriarty, N. W., Poon, B. K., Sobolev, O. V., Terwilliger, T. C., Adams, P. D. & Urzhumtsev, A. New tools for the analysis and validation of cryo-EM maps and atomic models. Acta Crystallogr D Struct Biol 74, 814–840, doi:10.1107/S2059798318009324 (2018).

116 Williams, C. J., Headd, J. J., Moriarty, N. W., Prisant, M. G., Videau, L. L., Deis, L. N., Verma, V., Keedy, D. A., Hintze, B. J., Chen, V. B., Jain, S., Lewis, S. M., Arendall, W. B., 3rd, Snoeyink,J., Adams, P. D., Lovell, S. C., Richardson, J. S. & Richardson, D. C. MolProbity: More and better reference data for improved all-atom structure validation. Protein Sci 27, 293–315, doi:10.1002/pro.3330 (2018).

117 Mohan, S. & Noller, H. F. Recurring RNA structural motifs underlie the mechanics of L1 stalk movement. Nature Communications 8, doi:10.1038/ncomms14285 (2017).

118 Goddard, T. D., Huang, C. C., Meng, E. C., Pettersen, E. F., Couch, G. S., Morris, J. H. & Ferrin, T. E. UCSF ChimeraX: Meeting modern challenges in visualization and analysis. Protein Science 27, 14–25, doi:10.1002/pro.3235 (2018).

119 Pettersen, E. F., Goddard, T. D., Huang, C. C., Meng, E. C., Couch, G. S., Croll, T. I., Morris, J. H. & Ferrin, T. E. UCSF ChimeraX : Structure visualization for researchers, educators, and developers. Protein Science 30, 70–82, doi:10.1002/pro.3943 (2020).

## Supplementary References

120 Walker, J. E., Saraste, M., Runswick, M. J. & Gay, N. J. Distantly related sequences in the alpha- and beta-subunits of ATP synthase, myosin, kinases and other ATP-requiring enzymes and a common nucleotide binding fold. EMBO J 1, 945–951, doi:10.1002/j.1460-2075.1982.tb01276.x (1982).

121 Loveland, A. B., Demo, G., Grigorieff, N. & Korostelev, A. A. Ensemble cryo-EM elucidates the mechanism of translation fidelity. Nature 546, 113–117, doi:10.1038/nature22397 (2017).

122 Urbatsch, I. L., Gimi, K., Wilke-Mounts, S. & Senior, A. E. Investigation of the role of glutamine-471 and glutamine-1114 in the two catalytic sites of P-glycoprotein. Biochemistry 39, 11921–11927, doi:10.1021/bi001220s (2000).

123 Adams, P. D., Afonine, P. V., Bunkoczi, G., Chen, V. B., Davis, I. W., Echols, N., Headd, J. J., Hung, L. W., Kapral, G. J., Grosse-Kunstleve, R. W., McCoy, A. J., Moriarty, N. W., Oeffner, R., Read, R. J., Richardson, D. C., Richardson, J. S., Terwilliger, T. C. & Zwart, P. H. PHENIX: a comprehensive Python-based system for macromolecular structure solution. Acta Crystallogr D Biol Crystallogr 66, 213–221, doi:10.1107/S0907444909052925 (2010).

